# SPT6L, a newly discovered ancestral component of the plant RNA-directed DNA methylation pathway

**DOI:** 10.1101/2024.01.20.576478

**Authors:** Vojtěch Čermák, Tomáš Kašpar, Lukáš Fischer

## Abstract

RNA-directed DNA methylation (RdDM) is driven by small RNAs (sRNAs) complementary to the nascent transcript of RNA polymerase V (Pol V). sRNAs associated with ARGONAUTE (AGO) proteins are tethered to Pol V mainly by the AGO-hook domain of its subunit NRPE1. We found, by *in silico* analyses, that Pol V strongly colocalizes on chromatin with another AGO-hook protein, SPT6-like (SPT6L), which is a known essential transcription elongation factor of Pol II. Our phylogenetic analysis revealed that SPT6L acquired its AGO-binding capacity already in the most basal streptophyte algae, even before the emergence of Pol V, suggesting that SPT6L might be a driving force behind the RdDM evolution. Since its emergence, SPT6L with the AGO-hook represents the only conserved SPT6 homolog in *Viridiplantae*, implying that the same protein is involved in both Pol II and Pol V complexes. To better understand the role of SPT6L in the Pol V complex, we characterized genomic loci where these two colocalize and uncovered that DNA methylation there is more dynamic, driven by higher levels of sRNAs often from non-canonical RdDM pathways and more dependent on chromatin modifying and remodeling proteins like MORC. Pol V loci with SPT6L are highly depleted in helitrons but enriched in gene promoters for which locally and temporally precise methylation is necessary. In view of these results, we discuss potential roles of multiple AGO-hook domains present in the Pol V complex and speculate that SPT6L mediates *de novo* methylation of naïve loci by interconnecting Pol II and Pol V activities.

## Introduction

The methylation of cytosines in DNA represents an important epigenetic mark, which mostly labels stably inactivated heterochromatin regions containing a high density of transposable elements (TEs). After DNA replication, several mechanisms ensure that the methylation pattern present at the parental DNA double helix is maintained in the newly synthesized strands. Hemimethylated symmetric CG contexts are methylated by MET1. Methylation in CHH and CHG contexts is maintained by CMT2 and CMT3, respectively, which are directed by histone H3K9me2 marks. Vice versa, this histone methylation is driven by SUVH4/5/6 enzymes that bind to methylated DNA, forming a positive self-reinforcing loop (Zhang et al., 2018a).

In addition, DNA methylation at many constitutively methylated loci, often situated at heterochromatin borders, is maintained by the activity of the RNA-directed DNA methylation pathway (RdDM; Böhmdorfer et al., 2016; Zhang et al., 2018a). Targets of RdDM are determined by sRNAs bound by Argonaute (AGO) proteins that search for complementarity on nascent transcripts of a specialized multisubunit DNA-dependent RNA Polymerase V. Pol V, which evolved from Pol II, is considered as an essential core component of the RdDM pathway because binding of AGO-sRNA to the Pol V transcript is necessary for the action of the RdDM-associated methyltransferase DRM1/2 (Wierzbicki et al., 2008; Zhong et al., 2014; Böhmdorfer et al., 2016). sRNAs involved in RdDM are mostly 24nt long and are produced by another plant-specific RNA polymerase, Pol IV, and associated enzymes (Daxinger et al., 2009). The pathway based on Pol IV-derived sRNAs is called canonical RdDM and mainly serves to maintain methylation after replication (Cuerda-Gil and Slotkin, 2016).

The RdDM pathway also allows for rapid and precise *de novo* introduction of DNA methylation marks even into previously active chromatin regions without preexisting DNA methylation. Thereby, RdDM can switch off transcription of suspicious genomic regions (e.g., newly integrated TEs) and also plays a key role in the dynamic methylation of promoter elements in a subset of developmentally and environmentally responsive genes (Fultz and Slotkin, 2017; Zhang et al., 2018a; He et al., 2022).

Maintenance methylation (after DNA replication) at the RdDM loci is likely mediated mainly by the attraction of Pol V to (hemi)methylated DNA by SUVH2/9 (Johnson et al., 2014). The recruitment of Pol V to naïve loci is less understood. Recently, two different explanations emerged, the first showing that Pol V sporadically transcribes even non-methylated regions of the *A. thaliana* genome (Tsuzuki et al., 2020) and the second indicating the role of RNA Pol II in attracting Pol V by an unknown mechanism that involves AGO proteins (Sigman et al., 2021).

AGO4 protein (and presumably all AGOs involved in RdDM) is tethered in the vicinity of the nascent transcript of Pol V by intrinsically disordered AGO-hook domains that typically contain variable repeats with conserved tryptophans in specific sequence contexts (mostly GW and WG; El-Shami et al., 2007; Karlowski et al., 2010). In *Arabidopsis thaliana*, such AGO-hook domains are present at the C-terminus of the largest subunit of Pol V (NRPE1) and also at its transcription elongation factor SPT5L, which participates in methylation of a large portion of loci methylated by the canonical RdDM pathway (Bies-Etheve et al., 2009). Also, two other AGO-hook proteins, NERD and SDE3, were suggested to influence RdDM at a small subset of loci (Garcia et al., 2012; Pontier et al., 2012).

Here we present *in silico* data analysis on SPT6-like (SPT6L), which carries one of the longest AGO-hook domains in *A. thaliana* (Karlowski et al., 2010). SPT6L is a plant-specific homolog of eukaryotic SPT6 proteins, known to act as the transcription elongation factors and histone chaperons in the Pol II complex (Antosz et al., 2017). In animals and other organisms, there is only the “canonical” SPT6 that does not contain the AGO-hook domain, while both SPT6 and SPT6L are present in *A. thaliana*. To reveal how the AGO-hook of SPT6L is involved in the regulation of gene expression, we reanalyzed available ChIP seq data (Chen et al., 2019) and discovered that SPT6L is not only a component of the Pol II complex but also colocalizes with Pol V at a large portion of its loci. Our phylogenetic analysis showed that SPT6L is the most ancient AGO-hook-bearing component of any RNA polymerase complex in the *Viridiplantae* clade and that, unlike *A. thaliana*, the vast majority of analyzed *Viridiplantae* species encode just SPT6L, while the “canonical” SPT6 without the AGO-hook is missing. Based on our thorough *in silico* analyses of genomic regions in which Pol V occurs together with SPT6L, we speculate about the possible roles of SPT6L in the RdDM pathway.

## Materials and methods

### Analyzed datasets

All NGS data analyzed in this study were downloaded from GEO (https://www.ncbi.nlm.nih.gov/geo/). Since the SPT6L ChIP-seq dataset that was key to our study came from 10-day-old seedlings, when searching for additional datasets, we aimed to have them as close to this stage as possible and selected the one that best matched with the SPT6L ChIP-seq data. All details about the analyzed datasets are listed in (Supplemental Table S1; Mi et al., 2008; Havecker et al., 2010; Luo et al., 2013; Johnson et al., 2014; Yelagandula et al., 2014; Zhang et al., 2016; Yang et al., 2017; Liu et al., 2018; Ma et al., 2018; Chen et al., 2019; Gallego-Bartolomé et al., 2019; Shu et al., 2019; He et al., 2021).

### ChIP-seq data analyses

Downloaded raw reads (Supplemental Table S1) were aligned to the TAIR10 reference genome using bowtie2 (v2.4.5; Langmead and Salzberg, 2012; https://github.com/BenLangmead/bowtie2; with parameters: -L 22 -i ‘S,1,1.15’ -- local). Peaks were called using macs2 callpeak (v2.2.7.1; https://github.com/macs3-project/MACS; with parameters: -- gsize ‘135000000’ --broad --broad-cutoff=’0.01’ --keep-dup ‘auto’ --qvalue ‘0.05’; with additional parameters: --nomodel --extsize #size --shift ‘0’ for single-end reads; the #size value was determined by macs2 predictd). In the case of more replicates, peaks were first called on pooled replicates and then on individual replicates, peaks from pooled replicates were then retained only when present also in individual replicates. The final set of significant peaks was filtered for those with two-fold enrichment or higher. Alignment files were filtered by samtools (v1.15.1; Danecek et al., 2021; https://github.com/samtools/samtools) to remove duplicate reads and coverage tracks were generated using deepTools bamCoverage (v3.5.1; Ramírez et al., 2016; https://github.com/deeptools/deepTools; with parameters: -bs 20 -- effectiveGenomeSize 135000000 --normalizeUsing RPGC --extendReads #size).

To obtain coverage over a set of loci, deepTools computeMatrix was used (with parameters: - m 500 -b 1000 -a 1000 -bs 20 --missingDataAsZero). The resulting data were then used to create metaplots (showing average and standard error) and heatmaps using deepTools, or the coverage over each locus was averaged and used to generate boxplots. ATAC-seq data were downloaded as coverage tracks (Zhong et al., 2021) and onward processed the same way as ChIP-seq data.

### DNA methylation analyses

Lists of DMRs were retrieved from Zhang et al., 2018b (260 datasets in total). To make the DMR data comparable with ChIP-seq peaks, the 100 bp long DMRs were merged to a single region if they were within 1 kbp from each other (such DMR regions overlap the ChIP-seq peaks in approximately 1:1 ratio). Bedtools intersect (v2.30.0; Quinlan and Hall, 2010; https://github.com/arq5x/bedtools2) was used to identify the DMRs/peaks with overlap. After the DMRs/peaks were counted for all the categories, the enrichment was calculated using the following formula: enrichment = log_2_(observed/expected); where expected = (total # of all P5 peaks with DMR overlap)*(P5 peaks of interest / all P5 peaks). The statistical significance of overlap was evaluated using hypergeometric distribution and the resulting p-value was corrected for multiple testing using Benjamini-Hochberg procedure. Samples were sorted into categories manually based on available literature, all the samples and their categories are listed in Supplemental Table S3.

The methylation tracks were retrieved from GEO (GSE39901; Stroud et al., 2013). Absolute values of methylation levels were calculated (ignoring the strandedness of methylation data). The procedure to obtain methylation level over a set of regions was similar as described in ChIP-seq analyses (with computeMatrix parameters: -m 500 -b 1000 -a 1000 -bs 20).

### sRNA-seq analyses

Downloaded raw reads (Supplemental Table S1) were trimmed with gsat (v0.1; https://github.com/MikeAxtell/gsat) using default settings. The read alignment, sRNA cluster identification and read counting was done by ShortStack (v3.8.5; Axtell, 2013; https://github.com/MikeAxtell/ShortStack; with parameters: --genome file TAIR10.fasta -- dicermin 20 --dicermax 25 --nohp for discovering new loci and with an additional parameter: --locifile loci_of_interest.bed for counting reads over analyzed loci). The resulting readcounts were normalized to the dataset size as reads per million (RPM).

### Loci analysis and annotation

To analyze loci where SPT6L and sRNAs overlap, bedtools intersect (default settings) was used to identify the intersection between the sRNA loci and SPT6L loci (i.e. SPT6L∩sRNA loci are parts of the original loci with exact overlap; Supplemental Fig. S1A). With this definition of loci, the analysis is done only at regions containing SPT6L and directly targeted by sRNAs, ignoring the surrounding non-overlapping parts (usually parts of SPT6L peaks, which are bigger), where we could not expect the sRNAs-SPT6L interaction to have an effect. These loci were then randomly shuffled with bedtools shuffle to provide loci for plotting background.

To classify Pol V loci based on the overlap with either SPT6L or Pol II, bedtools intersect (parameters -wa or -v) was used (i.e. in this case we looked at whole Pol V loci, not just their overlapping parts; Supplemental Fig. S2B). This approach was used because the goal was to compare different categories of the Pol V loci.

Pol V loci were annotated with associated genomic features (genes, pomoters, TEs etc.) using uropa (v3.3.4; Kondili et al., 2017; https://github.com/loosolab/UROPA; with parameters: “feature.anchor”:[“start”, “center”, “end”], “distance”:[0, 0], “internals”:“0.5”). The loci were annotated with the features independently, so each Pol V locus could be classified with multiple features. Promoter was defined as region 1 kbp upstream of TSS of the gene. Terminator was defined as 3’UTR plus 500 bp downstream. Araport11 was used for gene annotations and TE Transcripts v1.0 (Panda et al., 2016; https://github.com/KaushikPanda1/AthalianaTETranscripts) was used to annotate TEs. This TE annotation was also a source for all presented TE characteristics (length, distance to nearest gene, type of silencing etc.). The enrichment of certain features at the loci of interest was calculated using formula: enrichment = log_2_(observed/expected); where expected = (total # of all P5 peaks with genomic feature overlap)*(P5 peaks of interest / all P5 peaks). The statistical significance of enrichment was evaluated using hypergeometric distribution. For statistical comparison of quantitative characteristics (e.g. ChIP-seq enrichment, loci length…) between two sets of loci Mann–Whitney U test was used.

### Protein binding predictions

Protein sequences were retrieved from UniProt (https://www.uniprot.org/), chain IDs are listed in Supplemental Table S2. The protein complex predictions were done with AlphaFold-Multimer (v2.2; Evans et al., 2022; https://github.com/deepmind/alphafold; the model was run in full_dbs mode with default parameters). AlphaFold-Multimer produces ipTM score, for which in range between 0.7 and 0.8 over 60 % of complexes exceed DocQ 0.49 (medium quality) and over 80 % of complexes exceed DockQ 0.23 (acceptable quality; Basu and Wallner, 2016; Evans et al., 2022). The protein structures were visualized by ChimeraX (v1.5; Pettersen et al., 2021; https://www.rbvi.ucsf.edu/chimerax/). The presence of conserved SH2-binding motives in the protein sequences in question was checked by blastp (https://blast.ncbi.nlm.nih.gov/Blast.cgi).

### Orthologs identification

Published sequences were retrieved from NCBI (https://www.ncbi.nlm.nih.gov/), Phytozome v12.0, (https://phytozome.jgi.doe.gov), and Phycocosm (https://phycocosm.jgi.doe.gov).

*Arabidopsis thaliana* protein sequences of AT1G65440 (SPT6L), AT5G04290 (SPT5L), AT1G05460 (SDE3), AT2G16485 (NERD), AT4G35800 (NRPB1 - Pol II), AT2G40030 (NRPE1 - Pol V), AT1G63020 (NRPD1 - Pol IV), AT5G49160 (MET1), AT5G14620 (DRM2), and AT1G69770 (CMT3) were used as queries for searches against genomes spanning all *Viridiplantae* with main focus on basal streptophyte algae. As DNMT3 is lost in angiosperms the protein sequence of DNMT3 from *Ceratopteris richardii* was identified and used instead. Manual checks against protein annotation were made where needed to identify correct proteins.

When needed FGENESH+ (Softberry Inc., New York, NY USA; www.softberry.com) was used to predict gene coding sequences from genomic sequences using the *A. thaliana* protein model when specific annotations were unavailable.

### AGO-hook domain annotation

After obtaining sequences of desired paralogs, their AGO-hook was identified by using Wsearch (v1.0; Zielezinski and Karlowski, 2015; http://150.254.123.165/whub/) using a plant matrix. For a protein to be annotated as AGO-hook carrying we set the threshold for motifs score ≥ 6 and the number of such motifs must be ≥ 3 (Sheu-Gruttadauria and MacRae, 2018). For marginal cases manual check was made to see if the composed domain is biologically relevant and for annotation of *Klebsormidiophyceae* SPT6L see Supplemental Fig. S4H,I.

### Building a phylogenetic tree

Manual curation was necessary to combine overlapping BLAST hits or transcript ORFs in some cases. Orthology was assessed through reciprocal BLAST searches and PFAM protein domain prediction. The multiple sequence alignment of protein sequences identified in the previous step was done by MAFFT v7.455 using the parameters E-INS-I or L-INS-I in Geneious Prime 2023.1.2 (https://www.geneious.com). We manually checked the alignments to correct the misalignment of homologous regions.

Maximum-likelihood phylogenetic trees were constructed by RAxML-HPC (Stamatakis, 2014) tools version 8.2.12 using a General Time Reversible (GTR) model with a gamma distribution of rate heterogeneity, and 100 bootstrap replicas were used to obtain support values. To visualize the acquired trees FigTree v1.4.4 (http://tree.bio.ed.ac.uk/software/figtree/) was used.

## Results

### SPT6L localizes into Pol V loci

In *Arabidopsis thaliana* genome 22 genes are predicted to contain the AGO-hook domain (Karlowski et al., 2010), yet only a few of them have been assigned to some sRNA-related pathway so far; NRPE1, SPT5L, NERD, SDE3 and SUO; Onodera et al., 2005; Bies-Etheve et al., 2009; Garcia et al., 2012; Pontier et al., 2012; Yang et al., 2012). SPT6L (AT1G65440), which until today was known only as a transcription elongation factor of Pol II, belongs to proteins with the AGO-hook domain of unknown function. Because SPT6L binds to chromatin and all other chromatin binding AGO-hook proteins discovered so far are involved in RdDM, we hypothesized that it could be involved in this pathway as well. To address this question, we made use of published ChIP-seq data on SPT6L in *A. thaliana* (Chen et al., 2019) and combined them with sRNA-seq data (Ma et al., 2018) to identify genomic loci where SPT6L and sRNAs overlap (SPT6L∩sRNA; Fig. 1A, Supplemental Fig. S1A). Thereafter, we analyzed DNA methylation at the overlapping parts of these loci (Fig. 1B). Indeed, we saw there high cytosine methylation in all three sequence contexts as would be expected for the RdDM pathway, which contrasted with very low methylation level at SPT6L loci not overlapping with sRNAs. We looked also at other chromatin marks at these loci, but unlike DNA methylation, most of these had more or less intermediate levels at the overlapping loci compared to loci where SPT6L and sRNAs do not overlap (Supplemental Fig. S1B-D). To find the reason for increased DNA methylation at the overlapping loci, we checked available ChIP-seq data for Pol II (the known interactor of SPT6L) and key RdDM components (Pol IV, Pol V and AGO4). All the RdDM proteins are strongly enriched at the overlapping loci (Fig. 1C, Supplemental Fig. S1E,F), which brought up the possibility that besides Pol II, SPT6L also functions with Pol V. Indeed a cluster analysis of the SPT6L loci for SPT6L, Pol II and Pol V ChIP-seq data shows a clear formation of two clusters: in the first one, SPT6L colocalizes with Pol V and in the second with Pol II. Moreover, there is only a minor overlap between the Pol II and Pol V loci (Fig. 1D). Since Pol IV is also present at most of Pol V loci, both Pol IV and Pol V were possible candidates for the colocalization with SPT6L. To differentiate between these two possibilities, we looked at the Pol V loci, which are not enriched for Pol IV and vice versa. SPT6L was strongly enriched only at loci where Pol V was present but not at loci where Pol IV was without Pol V (Supplemental Fig. S1G,H). This together strongly suggests that SPT6L could be a component of the Pol V complex.

**Fig. 1:**
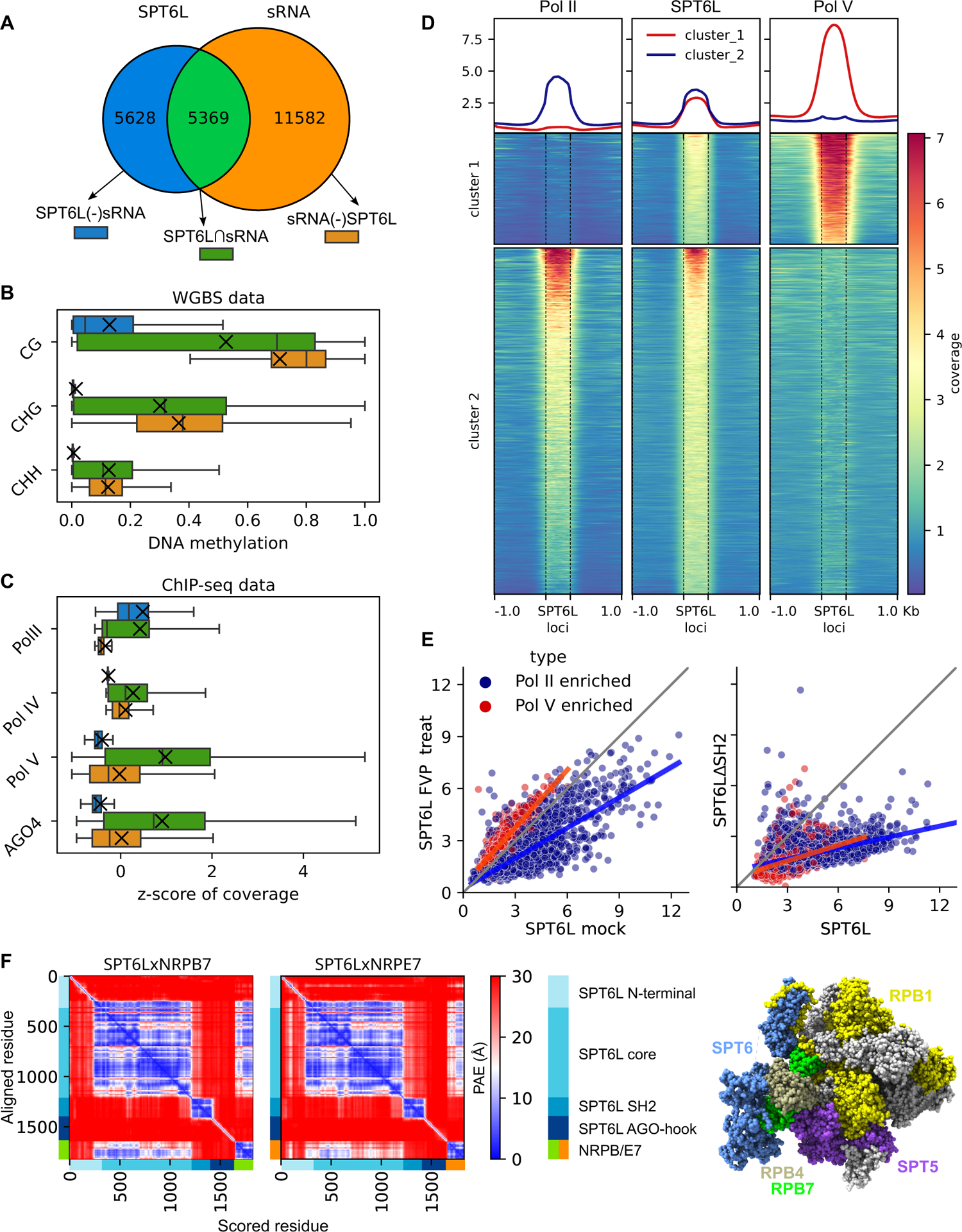
SPT6L is part of the Pol V complex (A-C) Analyses of loci where SPT6L and sRNAs overlap. (A) Venn diagram shows the numbers of the three analyzed groups of loci: SPT6L overlapping sRNA (SPT6L∩sRNA), non-overlapping SPT6L (SPT6L(-)sRNA) and non-overlapping sRNA (sRNA(-)SPT6L). (B,C) The boxplots show levels of DNA methylation (B) and z-score of ChIP-seq coverage for Pol II together with key RdDM components (Pol IV, Pol V and AGO4; C) at loci presented in panel (A). Medians are indicated by vertical lines and means by crosses. (D) Heatmap showing ChIP-seq data for Pol II (NRPB1), SPT6L and Pol V (NRPE1) at SPT6L loci (± 1 kbp of adjacent regions). The loci were clustered into two clusters using k-means clustering. (E) Scatterplot showing enrichment for SPT6L at SPT6L loci that overlap either only Pol II loci (in blue) or only Pol V loci (in red). Left graph: SPT6L in mock treated plants (x-axis) and flavopiridol (an inhibitor of Pol II kinase P-TEFb) treated plants (y-axis). Right graph: SPT6L in WT plants (x-axis) and in plants with SPT6L mutated in its SH2 domain (SPT6LΔSH2, y-axis). (F) Left: Predicted Aligned Error (PAE) for AlphaFold structure prediction of the SPT6LxNRPB7 and SPT6LxNRPE7 complexes (showing the best scoring models). Peptide chains and their relevant domains are shown at the x- and y-axes. Right: SPT6 and interacting proteins in the RNA polymerase complex depicted on the 3D structure of mammalian Pol II (PDB ID: 6GMH; Vos et al. 2018). SPT6 in blue, non-interacting subunits in gray and interacting subunits in remaining colors.

The interaction between SPT6/SPT6L (called SPT6(L) onwards) and Pol II is facilitated by the SH2 domain of SPT6(L) that binds to the phosphorylated serine, threonine and tyrosine residues at the linker to the C-terminal domain (CTD) of NRPB1, the largest subunit of the Pol II complex (Sdano et al., 2017; Vos et al., 2018; Chen et al., 2019). We checked for conserved SH2-binding motifs (residues) at the paralogous Pol V subunit NRPE1 but we did not find any. However, we were also unable to find these motives in the largest subunit of yeast Pol I (data not shown), which has also been indicated to bind SPT6 (Engel et al., 2015). To further address the means of possible SPT6L binding to Pol V, we reanalyzed ChIP-seq data for *A. thaliana* treated with flavopiridol (an inhibitor of Pol II kinase P-TEFb, responsible for phosphorylation of the NRPB1 CTD linker; Vos et al., 2018; Chen et al., 2019) and an *A. thaliana* mutant lacking the SH2 domain of SPT6L (SPT6LΔSH2; Chen et al., 2019). As expected, we could see a decrease in SPT6L binding to loci occupied by Pol II when the samples were treated with the kinase inhibitor. In contrast, at loci occupied by Pol V, the treatment caused a slight increase in SPT6L binding (Fig. 1E). Surprisingly, the missing SH2 domain reduced the binding of SPT6L to both Pol V and Pol II occupied loci (Fig. 1E). This shows that the SH2 domain affects SPT6L binding even to loci where only Pol V is present but this binding is not dependent on the phosphorylation machinery that normally deposits phosphate groups recognized by the SH2 domain.

Although NRPB1 CTD linker is important for the binding of SPT6(L) to Pol II, a much larger interacting interface between SPT6(L) and Pol II is at NRPB4/7 subunits and also at its elongation factor SPT5 (Fig. 1F; Vos et al., 2018; Chen et al., 2019). To assess whether SPT6L may bind these interfaces also at the Pol V complex we used the AlphaFold-Multimer predictions (Evans et al., 2022). Besides predicting the secondary structure of proteins or complexes, AlphaFold also calculates the accuracy of such structures. We used this score to estimate the likeliness of SPT6L interaction with Pol V subunits. Because assembling the whole polymerase complex using AlphaFold would be extremely demanding on computational resources due to its size, we resorted to testing pairwise interactions. We first ran the prediction for SPT6L and selected interacting subunits of the Pol II complex (NRPB1/4/6/7/SPT5) to see how AlphaFold will perform (Supplemental Table S2). Of these, NRPB7 showed the best-supported interaction (Fig. 1F). Therefore, we ran further predictions for all NRPB7 paralogues (Supplemental Table S2). In this analysis NRPE7 reached the highest score, very similar to that of NRPB7 (Fig. 1F; Supplemental Fig. S1I). The fact that NRPE7 probably retained the ability to interact with SPT6L throughout the evolution further supports our hypothesis that SPT6L is a component of the Pol V complex.

We also tried to search for any co-immunoprecipitation experiments (co-IP LC-MS/MS) that would indicate Pol V-SPT6L interaction. Unfortunately, from many co-IP studies involving Pol V only one presented the full dataset (Li et al., 2018). However, this study failed to detect SPT6L even in the Pol II complex, indicating that probably their co-IP conditions were not favorable to retain SPT6L in the immunoprecipitated complex.

### SPT6L is connected with euchromatic RdDM loci characterized by dynamic DNA methylation often targeted by non-canonical pathways

SPT6L may participate in the Pol V complex as a transcription elongation factor with histone chaperone activity and besides that, it brings a third AGO-hook to the complex in addition to those already present on NRPE1 and SPT5L. SPT6L is significantly enriched only at some Pol V loci (Fig. 2A), so determining the characteristic features of these loci would help to better understand the role of SPT6L in the RdDM pathway. For this purpose, we have defined Pol V loci as peaks from ChIP-seq data (Liu et al., 2018) and classified them based on their overlap with similarly defined SPT6L and also Pol II loci (Chen et al., 2019). In total, we have identified 8574 Pol V loci, of these 3472 loci (40%) are overlapped by SPT6L loci. We call these Pol V loci P5(+)SPT6L, in contrast to Pol V loci without SPT6L, which we call P5(-)SPT6L. Within the P5(+)SPT6L loci, 997 loci (29%; 11.6% of all Pol V loci) also overlap with Pol II loci (Fig. 2A, Supplemental Fig. S2A,B). At these loci, SPT6L can also be present as a component of the Pol II complex, so to filter this out in all the subsequent analyses, we will also separately provide results for P5(+)SPT6L loci without the overlap with Pol II, i.e. P5(+)SPT6L(-)P2 loci as supplementary data.

**Fig. 2:**
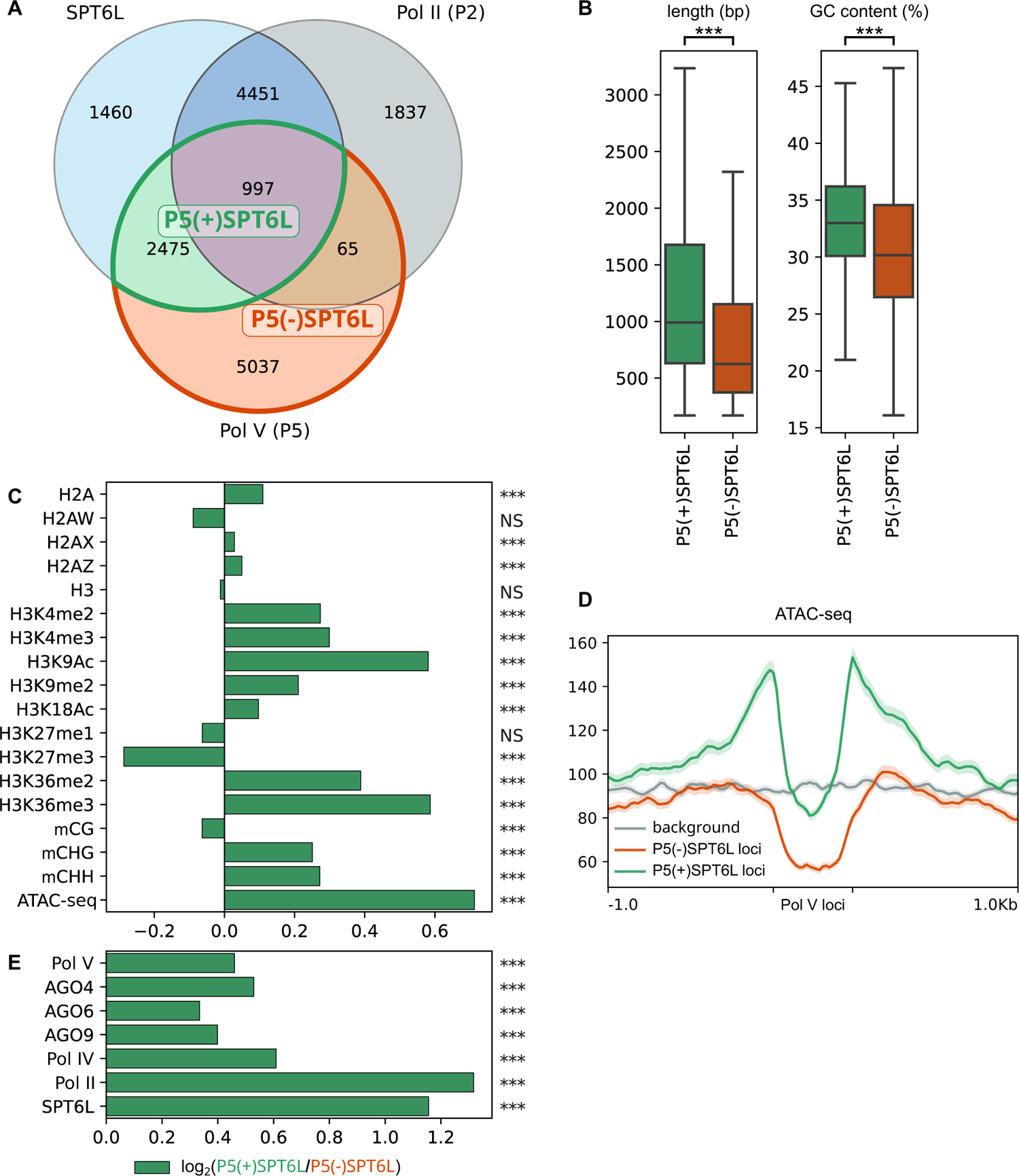
Characterization of Pol V loci occupied by SPT6L (A) Venn diagram with SPT6L, Pol II and Pol V loci with highlighted overlaps that are compared in the following panels. (B) Length and GC content of P5(+)SPT6L and P5(-)SPT6L loci. (C) Enrichment/depletion of selected chromatin marks at P5(+)SPT6L loci calculated as log_2_(P5(+)SPT6L/P5(-)SPT6L). (D) Metaplot comparing ATAC-seq signal at P5(+)SPT6L and P5(-)SPT6L loci and in their vicinity, randomized P5 loci are used for background. (E) Enrichment of RdDM components, SPT6L and Pol II at P5(+)SPT6L loci (calculated as in panel C). Significance is indicated by asterisks: p < 0.0005 = ***; NS = not significant; in Fig. 2C,E the statistics were calculated from the normalized coverages at the P5(+)SPT6L and P5(-)SPT6L loci from which the log_2_ ratio was calculated.

The Pol V loci with and without SPT6L were compared in four different aspects; i) chromatin state (components and modifications), ii) the mode of DNA methylation (proteins involved in its establishment/maintenance); iii) small RNAs mapping to the loci, and iv) functional classification of genomic elements overlapping with the loci.

### Chromatin characterization

The P5(+)SPT6L loci are longer when compared to P5(-)SPT6L, and they also have somewhat higher GC content but still much lower than what is typical for Pol II loci (Fig. 2B, Supplemental Fig. S2C). Both types of Pol V loci show depletion of transcription activation marks compared to the average level in the genome but the P5(+)SPT6L loci have higher levels of these marks, particularly in the adjacent regions where they are even over the average genome level, which is most evident for the chromatin accessibility (ATAC-seq) data (Fig. 2C,D, Supplemental Fig. S2D,E). In the case of transcription repressive marks, the P5(+)SPT6L loci are more enriched for H3K9me2 while P5(-)SPT6L are more enriched for H3K27me3 (Fig. 2C, Supplemental Fig. S2D,E).

At the P5(+)SPT6L loci, all the RdDM components are also more enriched compared to P5(-)SPT6L loci, probably because Pol V itself is more enriched at these loci (Fig. 2E, Supplemental Fig. S2D,F). Accordingly, the P5(+)SPT6L loci also show higher levels of DNA methylation in the CHG and CHH context but no difference in the CG context (Fig. 2C, Supplemental Fig. S2D,E). Both higher enrichment in RdDM components and increased methylation at P5(+)SPT6L loci, which are surrounded by accessible chromatin, might indicate a role of SPT6L in either stabilizing or recruiting the Pol V complex into its target loci embedded in more active chromatin.

### Mode of DNA methylation

To better understand the presumed role of SPT6L in DNA methylation, we asked which other proteins are involved in establishment or maintenance of DNA methylation at Pol V loci with and without SPT6L. To answer this, we looked at how DNA methylation is affected by mutations in various genes related to chromatin modifications at these loci. More specifically we looked if differentially methylated regions (DMRs) from the given sample (mutant) are more enriched at either P5(+)SPT6L or P5(-)SPT6L loci (by reusing data from Zhang et al., 2018b). In all samples (mutants), decreases/losses of DNA methylation (hypo-DMRs) were significantly more frequent at the P5(+)SPT6L loci. Surprisingly, gains/increases in DNA methylation (hyper-DMRs) were also significantly enriched for most of the samples at the P5(+)SPT6L loci. Only for the CG hyper-DMRs most samples did not show any significant enrichment at all. In summary, no sample showed significant depletion for hypo-DMRs and only few for hyper-DMRs at the P5(+)SPT6L loci (Fig. 3A,B, Supplemental Fig. S3A,B, Supplemental Table S3) which indicates that proper methylation at these loci is highly sensitive and practically any disruption in the methylation machinery causes hypo- or hypermethylation of the loci.

**Fig. 3:**
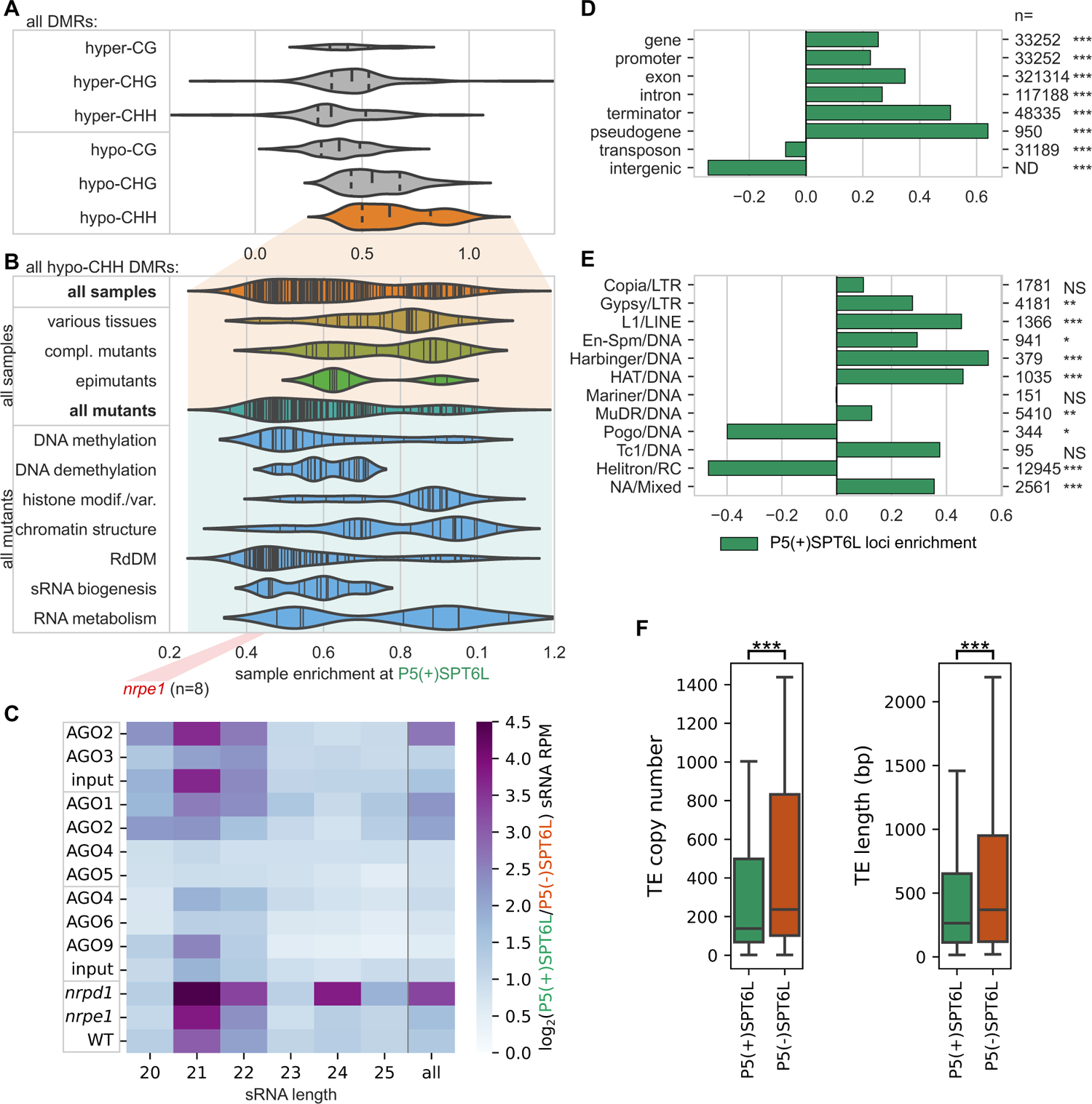
Mode of DNA methylation, types of sRNAs and genomic elements at Pol V loci occupied by SPT6L (A,B) Enrichment/depletion of differentially methylated regions (DMRs) at P5(+)SPT6L loci in datasets/samples from Zhang et al., 2018b containing whole genome methylation data for various mutants and some tissues. Violin plots show how the individual samples are enriched at P5(+)SPT6L loci (i.e. whether DMRs from the sample tend to overlap the P5(+)SPT6L loci more than the P5(-)SPT6L loci): (A) sample assessment based on DMR type (cytosine context and hyper-/hypomethylation) and by (B) the sample category that is shown only for hypo-CHH DMRs in the sample. Only samples significantly enriched/depleted for the P5(+)SPT6L loci are shown. In panel A the width of the violin plot is scaled to the number of samples and vertical lines indicate median and quartiles, in panel B the plots are not scaled and vertical lines indicate individual samples; position of *nrpe1* mutant samples is outlined below by the red rectangle. Data for individual samples and their classification can be found in Supplemental Table S3. (C) Heatmap with enrichment of sRNA at P5(+)SPT6L loci for indicated samples and sRNA lengths calculated as log_2_(P5(+)SPT6L/P5(-)SPT6L). Analyzed sRNA-seq datasets originate either from AGO immunoprecipitation or from total sRNA samples in case of “input” and WT, and *nrpe1* and *nrpd1* mutants. Samples outlined within the same rectangle come from the same experiment (these are well comparable with each other). Individual replicates are shown in Supplemental Table S4. (D,E) Enrichment/depletion of the overlap of P5(+)SPT6L loci at (D) different types of genomic elements and (E) various TE families. The “n” indicates the total number of genomic elements or TEs in a given category. (F) Copy numbers and lengths of TEs overlapping either P5(+)SPT6L or P5(-)SPT6L. Significance is indicated by asterisks: p < 0.05 = *; p < 0.005 = **; p < 0.0005 = ***; NS = not significant. The significance indicates either enrichment of P5(+)SPT6L in panels D and E, or significant difference between P5(+)SPT6L and P5(-)SPT6L in panel F.

In the *nrpe1* mutant with completely inactivated RdDM pathway, enrichment of CHH hypo-DMRs at the P5(+)SPT6L loci reached 0.41-0.46 (the enrichment is expressed as log_2_ fold-change, i.e. 1.33-1.38 times more CHH hypo-DMRs overlap the P5(+)SPT6L loci than expected from proportional distribution; Fig. 3B, Supplemental Fig. S3B, Supplemental Table S3). As expected, mutations in the essential components of the core canonical RdDM pathway scored about the same as the *nrpe1* mutant, with the lowest enrichment in *suvh2* and *fve* mutants, both genes involved in tethering Pol V to already methylated DNA (Supplemental Table S3; Johnson et al., 2014; Zhou et al., 2021). The remaining mutants from Zhang’s dataset scored higher than core RdDM mutants. Somewhat higher enrichment was observed for mutants in RdDM auxiliary factors, genes involved in DNA methylation and demethylation, and genes involved in sRNA generation and processing (besides the canonical RdDM pathway). The highest enrichment of CHH hypo-DMRs at the P5(+)SPT6L loci was observed for many mutants in factors involved in chromatin remodeling and histone modifications/histone variants (Fig. 3B, Supplemental Fig. S3B). To be more specific, mutants in MOM1/MORC complex, H3K4 demethylation (JMJ14, SDG8), histone deacetylation (HDA6) and CMT3/KYP self-reinforcing loop typical for maintenance methylation, had CHH hypo-DMRs most enriched at P5(+)SPT6L loci compared to the P5(-)SPT6L loci. Most of these mutants, however, affected only a small fraction of *nrpe1* hypo-DMRs (Supplemental Table S3). Besides enrichment for various mutants (and complemented mutants), we also observed very high enrichment for CHH hypo-DMRs at the P5(+)SPT6L loci in samples from various plant tissues, organs, and developmental stages (Fig. 3B, Supplemental Fig. S3B). All this indicates that SPT6L is present at the RdDM loci with chromatin state dynamically changing (like during development) and that such loci are very sensitive to any disturbances in the machinery responsible for their methylation.

### Small RNAs

Since SPT6L has an AGO-hook domain and sRNAs are key components of the RdDM pathway, we also looked at what type of sRNAs do map to these loci, into which AGOs are they loaded, and how these sRNAs are influenced in mutants of Pol IV and Pol V (*nrpd1* and *nrpe1*, respectively; Fig. 3C, Supplemental Table S4). The P5(+)SPT6L loci are overall targeted by higher levels of sRNAs. This difference is even more pronounced in the case of sRNAs loaded into AGO1 and AGO2 and less so in AGO6 and AGO9-loaded sRNAs. In agreement with this, P5(+)SPT6L loci are more enriched in 21nt and 22nt sRNAs typical for the non-canonical RdDM pathways (Cuerda-Gil and Slotkin, 2016). We also observe higher sRNAs enrichment at the P5(+)SPT6L loci in the *nrpd1* mutant with non-functional Pol IV (Fig. 3C, Supplemental Table S4). All of this suggests that although P5(+)SPT6L loci are more occupied by Pol IV compared to P5(-)SPT6L loci (Fig. 2E), they are even more strongly targeted by Pol II-derived sRNAs. A lot of the above-mentioned differences are less prominent (but still visible) if we exclude P5(+)SPT6L loci which overlap with the Pol II loci, i.e. P5(+)SPT6L(-)P2 from the analyses (Supplemental Fig. S3C, Supplemental Table S4). It indicates that the P5(+)SPT6L loci are not only more specifically targeted by Pol II-derived sRNAs but some of them may also serve as their sources.

### Genomic elements

Pol V is known to target mostly various types of transposable elements (TEs) and also gene promoters (Zheng et al., 2012), so we further analyzed if the presence of SPT6L is enriched at some of these targets. Indeed, P5(+)SPT6L loci show higher overlap with all gene-related features compared to P5(-)SPT6L loci (Fig. 3D). This overlap is lower for the P5(+)SPT6L(-)P2 loci with the exception of promoters and pseudogenes, where the absence of Pol II has no negative effect on the enrichment (Supplemental Fig. S3D). On the other hand P5(-)SPT6L loci show higher overlap with intergenic regions and TEs. A closer look at the TEs, however, shows higher enrichment for many DNA and LTR elements at the P5(+)SPT6L loci, but strong depletion for Helitrons, the largest TE family in *A. thaliana* (Fig. 3E, Supplemental Fig. S3D). Generally TEs at P5(+)SPT6L loci have lower copy numbers, are shorter and are more frequently silenced by various non-canonical RdDM pathways (Fig. 3F, Supplemental Fig. S3D,E). The TE length bias does not seem to be related to the preferred TE families. In summary, SPT6L is more frequently present at Pol V loci connected with shorter and rarer TEs silenced by non-canonical RdDM, and also with genes and gene promoters for which locally and temporally precise methylation is of particular importance.

### SPT6L was the first AGO-binding component of any plant RNA polymerase complex

Protein SPT6 has two paralogues in *A. thaliana*: SPT6L (AT1G65440) with the AGO-hook domain and SPT6 (AT1G63210) without this domain. The first one is likely part of the Pol V complex as shown in the previous chapters and also of the Pol II complex (Chen et al., 2019), while the second paralogue is not very well described but it might be a part of the Pol II complex (Antosz et al., 2017). It somewhat resembles the situation with a pair of paralogues of SPT5(L), another RNA polymerase elongation factor (non-homologous to SPT6L). SPT5 without the AGO-hook functions in the Pol II complex (a role conserved to all eukaryotes), while SPT5L with the AGO-hook acts in the Pol V complex (a function specific to plants; Bies-Etheve et al., 2009). To gain a better understanding of the evolution of SPT6 paralogues, we compared detailed phylogenetic trees for SPT5(L) and SPT6(L) (Fig. 4, Supplemental Fig. S4A,B).

**Fig. 4:**
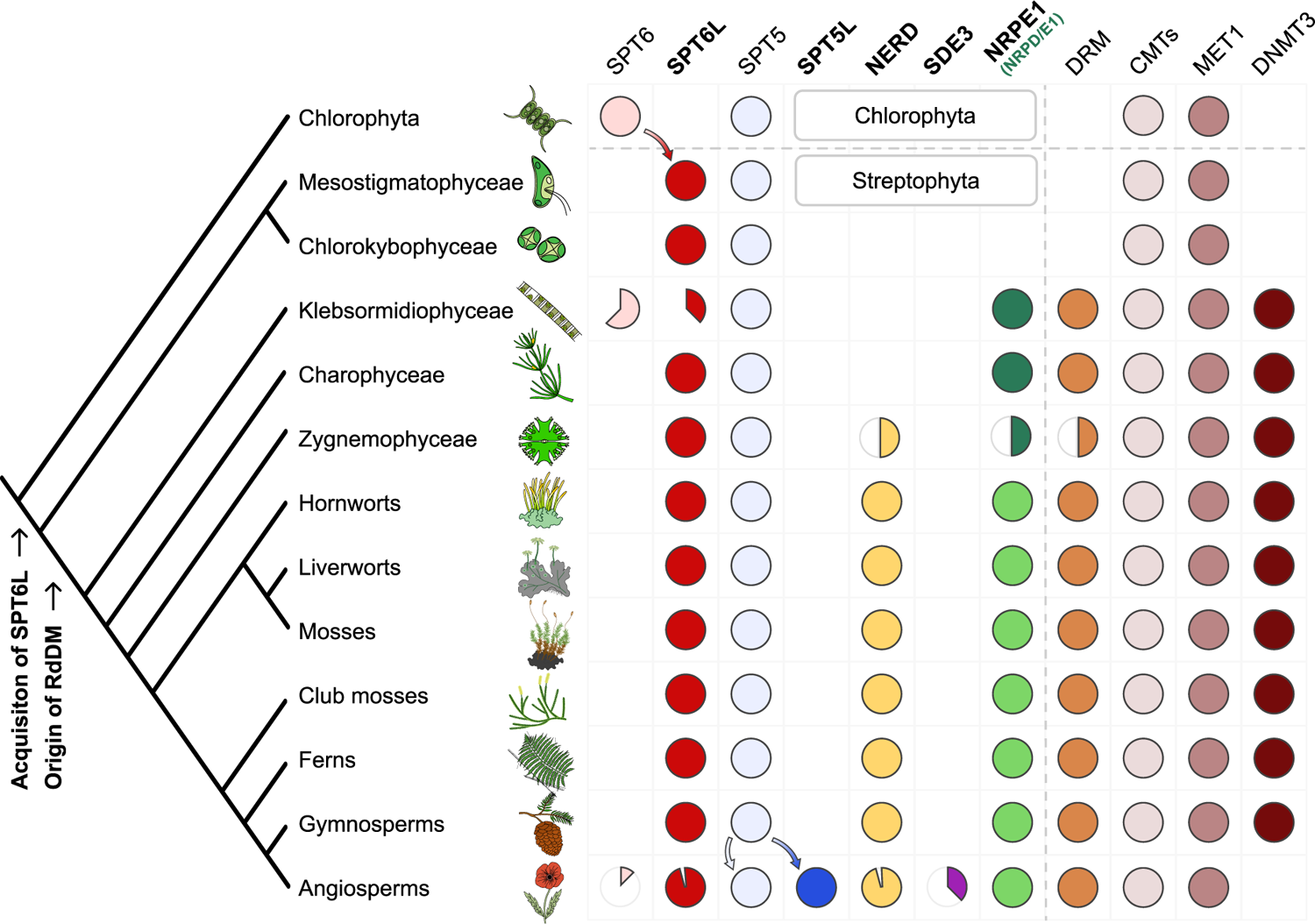
Evolution of SPT6L, other relevant AGO-hook proteins and DNA methyltransferases in *Viridiplantae* A schematic tree depicting stepwise acquisitions and presumed secondary losses of AGO-hook proteins and cytosine methyltransferases in *Viridiplantae* with focus on the early evolution of RdDM components in the streptophyte algae. The phylogenetic tree is adapted from (Leliaert et al., 2012). Filled circles indicate that the given protein was found in the taxa, and the colored sector depicts the approximate ratio of losses or gains in members of the taxa. The SDE3 sector shows the proportion of SDE3 proteins with AGO-hook and does not include SDE3 homologues without the AGO-hook (SDE3-like proteins), similarly the empty sector in NERD indicates only loss of the NERD’s AGO-hook not of the whole gene. The asterisk in the class *Klebsormidiophyceae* denotes possible secondary loss of the AGO-hook in the *Klebsormidium* genus; this special case is described in detail in the Supplemental Fig. S4H,I. It should be also noted that although the DNMT3 is present in many eukaryotes, it is absent in many algae, including the outgroup used in this tree (Goll and Bestor, 2005). Detailed trees for the individual proteins that were the basis for this schematic summary tree are available as Supplemental Fig. S4.

These trees indicate that SPT5 diverged in angiosperms (Supplemental Fig. S4B; Matzke et al., 2015) into two separate clades characterized by the presence/absence of the AGO-hook, with both paralogues detected in all analyzed angiosperm species. On the contrary, there is no such deep dichotomy in the phylogenetic tree of SPT6(L). The only conserved paralogue in plants is SPT6L with the AGO-hook domain (Supplemental Fig. S4A). Occasional occurrences of SPT6 without the AGO-hook reflect recent parallel duplications (in some *Brassicaceae* and a few other clades), likely followed by a secondary loss of the AGO-hook domain in one of the paralogues. SPT6L with the AGO-hook is missing only in *Carica papaya*, where surprisingly, also NERD, another AGO-hook protein related to the RdDM pathway, is present without the AGO-hook domain, and even SPT5L has its AGO-hook significantly truncated (only 13 motives compared to usual 50+ motives in other species; Supplemental Fig. S4A,B,C).

As already mentioned, SPT6L is just one of several AGO-hook-carrying proteins possibly affecting the RdDM pathway (NRPE1, SPT5L, NERD, SDE3). To better understand the RdDM evolution in the context of its newly identified component (SPT6L), we searched for the first emergence of all these proteins in the *Viridiplantae* clade (Fig. 4).

Protein SDE3 obtained the AGO-hook most recently and possesses this domain only in some angiosperm families, specifically in *Asteraceae* and *Brassicaceae* and its function is only partially connected with DNA methylation (Garcia et al., 2012; Supplemental Fig. S4D).

The previously mentioned SPT5L likely originated from SPT5 duplication in the ancestor of angiosperms (Supplemental Fig. S4B). Protein NERD is older and likely emerged in the common ancestor of *Zygnematophyceae* (detected in *Mesotaenium*), mosses and vascular plants. Interestingly, in gymnosperms that do not have SPT5L, NERDs are characterized by longer AGO-hook domains compared with other clades (Supplemental Fig. 4B,C).

The most ancient AGO-hook domain, which has been connected with RdDM, belongs to the largest subunit of Pol V, NRPE1 (more exactly the common ancestor of NRPD1/E1 of Pol IV and Pol V). It arose by duplication of the gene encoding the largest subunit of Pol II (NRPB1; Huang et al., 2015) and can first be found in the basal orders of streptophyte algae *Klebsormidiophyceae* and *Charophyceae* (Supplemental Fig. S4E; for details see Trujillo, 2019). The emergence of NRP(D)E1 in these algae is accompanied by the emergence of DRM methyltransferase, which is considered necessary for the RdDM pathway (Supplemental Fig. S4F). Although the duplication event giving rise to NRP(D)E1 likely happened shortly after the divergence of streptophyte algae from chlorophyte algae, the most basal clades of streptophyte plants represented by *Mesostigma* and *Chlorokybus* do not possess the NRP(D)E1 gene. Surprisingly, our phylogenetic analysis detected SPT6L with the AGO-hook motives already in these two algae (Supplemental Fig. S4A) in which the emergence of SPT6L coincides with the appearance of chromomethylases (Supplemental Fig. S4G), so the acquisition of the SPT6L’s AGO hook into the Pol II complex preceded the emergence of Pol V. In chlorophyte algae, only the canonical SPT6 was detected, making SPT6L a synapomorphy of *Streptophyta*. In summary, the four AGO-hook binding proteins have appeared successively at different stages of plant evolution and were often accompanied with gains or loss of some cytosine methyltransferase genes (Fig. 4). SPT6L appeared as the most ancient component of any RNA polymerase complex that acquired the ability to bind AGOs in *Viridiplantae*.

### A possible role for SPT6L in Pol V targeting

The absence of SPT6 in most plant species indicates that the same SPT6L with the AGO-hook is present in both Pol II and Pol V complexes. The fact that SPT6L did not diverge in evolution into specialized paralogues that would support these complexes separately suggests that the function of SPT6L in both complexes may be interconnected. SPT6L has been shown to have a role in chromatin opening, which is required for the initiation of Pol II transcription (Shu et al., 2022). The same may be true for Pol V at some of its loci. Pol V is predominantly attracted to its loci by SUVH2/9 that recognize methylated DNA (Johnson et al., 2014), likely primarily “hemimethylated” DNA after replication. In the case of unmethylated naïve loci, Sigman et al., 2021 suggested that Pol V could be guided by AGO4/6/9, which interact with Pol II transcripts. SPT6L, with its AGO-binding domain, histone-binding domain, and presumed ability to interact with both polymerases, seems like a good candidate to facilitate this process. Because Sigman et al., 2021 suggested that this AGO-dependent targeting should occur at loci where Pol V targeting is independent (or only partially dependent) on SUVH2/9, we took a closer look at these loci (Fig. 5A). Here we could observe much higher levels of Pol II, but only small differences in SPT6L binding when compared to loci fully dependent on SUVH2/9. SPT6L present at such loci is mostly not a part of the Pol II elongation complex, as the presence of SPT6L there is not reduced by the flavopiridol treatment. As expected, *suvh2/9* mutation causes depletion of NRPE1 (Pol V) from both types of loci, but the effect is much stronger in the case of SUVH2/9-dependent loci. Deletion of the SPT6L’s SH2 domain (SPT6LΔSH2) strongly reduced its binding at loci dependent on SUVH2/9. This was expected based on the previous analysis which showed that the SH2 domain is important for SPT6L binding to Pol V loci (Fig. 1E). Surprisingly, SPT6L level was almost unaffected at loci where Pol V binding can be independent of SUVH2/9. It is unclear how SPT6L can bind to these loci and if its binding there involves Pol II (even in absence of the SH2 domain). But since this binding is likely independent of Pol V, it opens the possibility to switch the roles so that SPT6L guides Pol V to these loci (Fig. 5B).

**Fig. 5.**
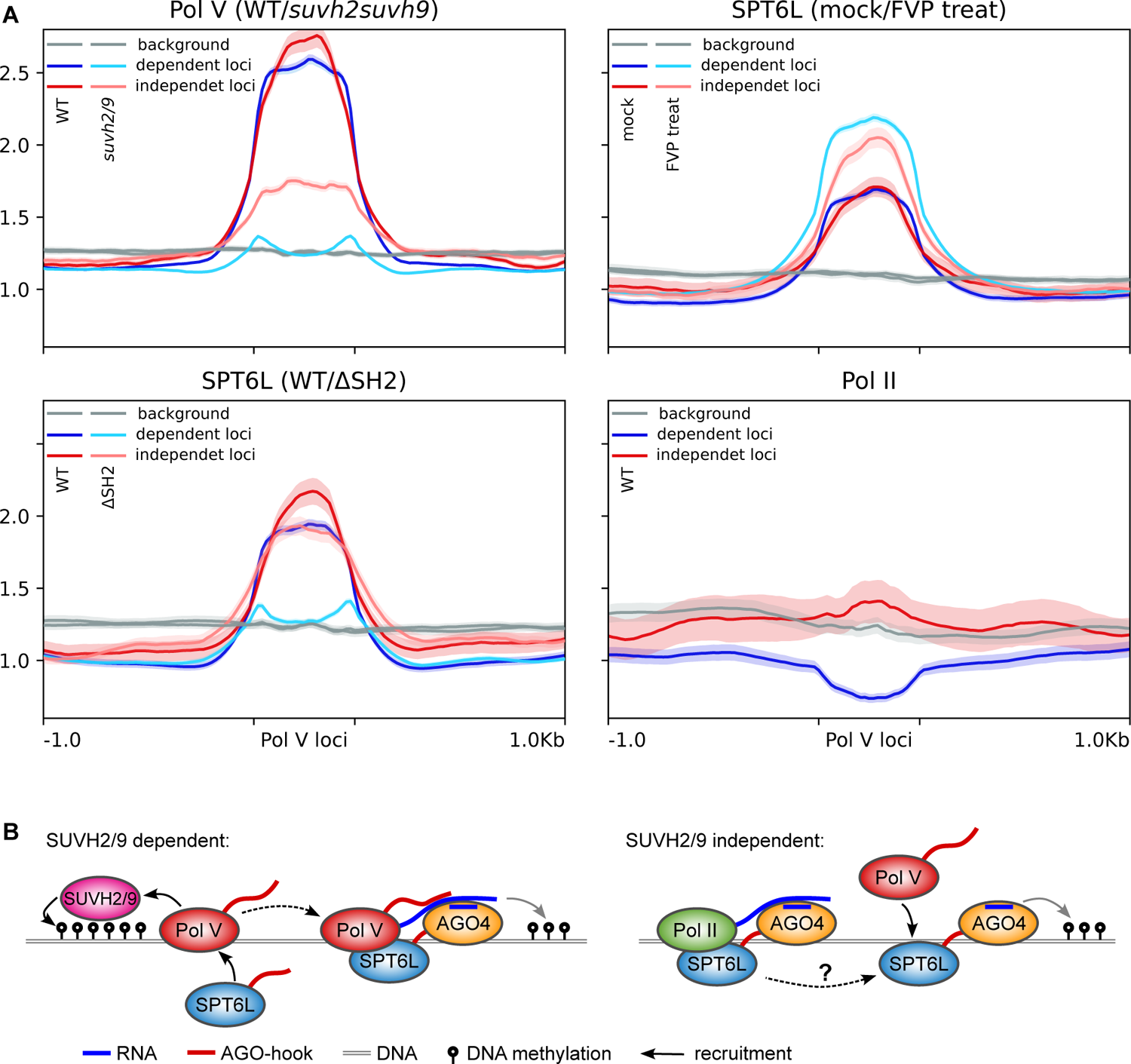
Potential role of SPT6L in tethering Pol V to chromatin (A) Metaplots showing the ChIP-seq data for Pol V loci categorized based on the dependence of Pol V targeting on SUVH2/9 (at “dependent loci” Pol V binding is completely lost in the *suvh2/9* mutant while at “independent loci” at least some Pol V is still deposited in the *suvh2/9* mutant; loci adopted from Sigman et al., 2021; ± 1 kbp of adjacent regions). The metaplots show ChIP-seq data of NRPE1 in WT and *suvh2/9* background, SPT6L with/without SH2 domain (WT/ΔSH2) or treated/mock treated with flavopiridol (mock/FVP treat) and Pol II. Background shows coverage at randomly shuffled Pol V loci. Semi-transparent color below/above the plot lines shows standard error. (B) Schematic model of Pol V recruitment to the SUVH2/9-dependent (left panel) and independent loci (model proposed by this study, right panel). At the SUVH2/9-dependent loci, Pol V is recruited exclusively through SUVH2/9 proteins and pre-existing DNA methylation as described earlier (Johnson et al., 2014) with the additional involvement of SPT6L as suggested by this study (Fig. 2A). At SUVH2/9 independent loci (right panel), SPT6L binds independently of Pol V, therefore it may be able to recruit Pol V there. Sigman et al., 2021 showed that AGO4 can bind to chromatin independently of Pol V, which is possibly mediated by Pol II transcripts. Thereafter AGO4 can recruit Pol V. Since SPT6L is a Pol II elongation factor that can independently bind to chromatin (Chen et al., 2019), and interact with AGO proteins and Pol V (this study), it is a good candidate to facilitate this process, but based on our data it is not clear how exactly SPT6L is recruited by Pol II (dashed arrow). At these loci Pol V binding can be subsequently also reinforced by SUVH2/9, which would be the same process as shown in the left panel.

## Discussion

In this study, we searched for the function and evolution of the AGO-hook domain of SPT6L which is a well-known plant homologue of a conserved transcription elongation factor of RNA Pol II. Throughout the genome of *Arabidopsis thaliana* SPT6L colocalizes with both Pol II and Pol V at mostly separate loci. We take this as a strong support for our hypothesis that SPT6L could function as Pol V elongation factor, alongside previously described SPT5L and SPT4 (SPT4 is shared with Pol II, Köllen et al., 2015). Our finding is also supported by a preprint (Liu et al., 2024) published in parallel to this study, where the authors show experimental evidence for the interaction between SPT6L and Pol V, confirming that SPT6L is indeed a part of RdDM. In our study, we further show that SPT6L is ancestral to RdDM and thoroughly characterize the Pol V loci where SPT6L is engaged.

### The first steps of the RdDM evolution

Our phylogenetic data indicate that SPT6L was the first component of any RNA Polymerase complex in the *Viridiplantae* clade that acquired an AGO-hook domain. We found SPT6L with this domain already in the most basal streptophyte algae *Chlorokybus* and *Mesostigma*. *Mesostigma viride* genome contains only about 1% of CHH methylation (Liang et al., 2020), which indicates no (or only negligible) RdDM activity. This corresponds to the fact that these algae contain neither an orthologue of DRM nor NRPD1/E1 (the Pol V subunit), which is considered essential for RdDM. Both these genes first appeared in *Klebsormidiophyceae* and *Charophyceae* (Luo and Hall, 2007; Trujillo, 2019; Yaari et al., 2019, this study). This indicates that the AGO-hook domain of SPT6L in the two most basal groups of streptophyte algae (*Chlorokybophyceae* and *Mesostigmatophyceae*) was unlikely engaged in some ancestral form of RdDM. However, it might be a prerequisite for evolution of the pathway. As the Pol V evolved, it likely contained two AGO-hook components (SPT6L and NRPD/E1) from the very beginning (Fig. 6).

**Fig. 6.**
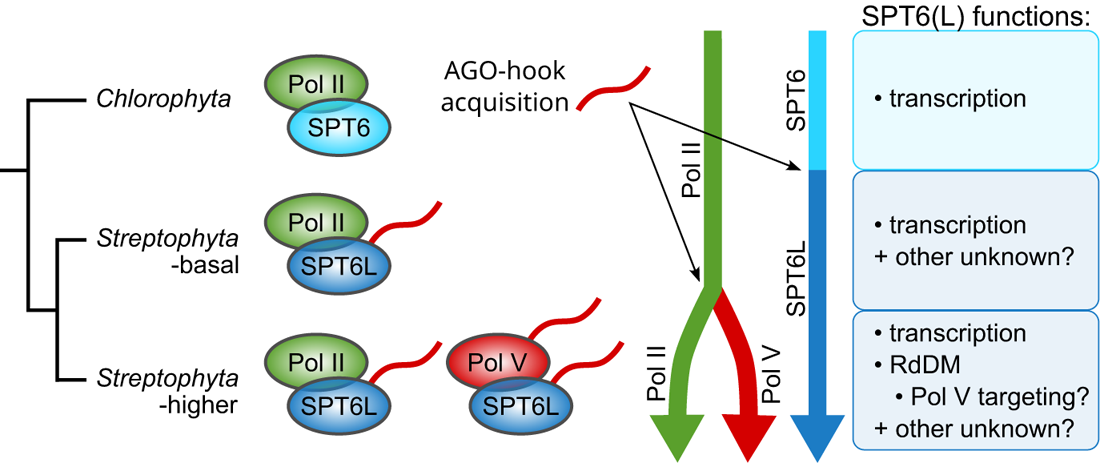
Diagram of the evolution of SPT6L functions and its relation to the RdDM pathway evolution Chlorophyte algae possess SPT6 without the AGO-hook domain, and it functions there as a Pol II elongation factor, the same as in other eukaryotic clades. In the most basal streptophyte algae, SPT6L acquired the AGO-hook domain which has there an unknown function. After the emergence of the RdDM pathway, SPT6L retained its functions in the Pol II complex, but also gained novel functions in the Pol V complex, which may include the recruitment of Pol V to unmethylated naive loci (Fig. 5B).

Functionally specialized Pol IV and Pol V, as we know them from higher plants, likely evolved in the most recent common ancestor of land plants (Wang and Ma, 2015). Although SPT6L was probably well established at this evolutionary stage, our analysis of available ChIP data shows that SPT6L strongly colocalizes only with Pol V but not with Pol IV. It can be related to differences in the processivity of these polymerases. Pol V produces longer transcripts that can span one or more nucleosomes (Blevins et al., 2015; Böhmdorfer et al., 2016). In agreement with this, Pol V loci with SPT6L are generally longer than those without. Although the AGO hook of SPT6L is likely to have some function in the Pol V complex, the mere presence of SPT6L in this complex may also be required for its histone chaperone activity (regardless of the AGO hook).

### Why are there so many AGO-hook proteins in the Pol V complex?

Identification of another AGO-hook component of the Pol V complex raises the question of why the Pol V complex does need so many. When considering that two or three tryptophans are sufficient to bind one AGO protein (Sheu-Gruttadauria and MacRae, 2018) and tens of such motives are present in the typical AGO-hook domain of NRPE1, SPT5L and also SPT6L (Supplemental Fig. S4A,B,E), the Pol V complex with all these components may have the capacity to bind more than ten AGOs at once. Small RNAs that drive methylation are generated from a huge number of genomic loci, so their sequence variability can be extremely high. The high number of AGO-binding sites may be necessary to maximize the likelihood that a nascent transcript of Pol V is recognized by an AGO that is just carrying a complementary sRNA within the short time window of ongoing transcription. In our previous study, we observed that the likelihood of *de novo* methylation of a naïve locus depended on the level of the respective sRNAs (Čermák et al., 2020) which indicates that this interaction can indeed be the limiting step of the whole process. In conclusion, we suggest that increasing the number of AGO-hook domains in plant evolution may be needed to maintain sufficient sensitivity of the RdDM pathway in the face of increasing sRNA and TE complexity.

However, AGO-hook of SPT6L does not necessarily function in attracting AGOs to the Pol V complex. Lahmy et al., 2016 showed that AGO-hooks of NRPE1 and SPT5L are fully redundant and their deletion leads to loss of RdDM dependent DNA methylation. This would either mean that the AGO-hook of SPT6L is not sufficient to attract AGOs on its own or it has other functions. It could function in attracting other proteins than AGOs, like it has been shown for GW182 in attracting CCR4-NOT complex (Chekulaeva et al., 2011). AGO-hook domains are intrinsically disordered, so they likely tend to participate in (or even directly induce) the formation of biomolecular condensates in the nucleus via the liquid-liquid phase separation (LLPS). The overall protein composition of these condensates can be determined by specific combinations of participating intrinsically disordered domains (Laflamme and Mekhail, 2020). Thus various combinations of the three (or even more) AGO-hooks associated with the Pol V complex (NRPE1, SPT5L, SPT6L) might participate in defining specific LLPS condensates that attract/contain a specific wider set of chromatin-binding and chromatin-modifying proteins which directly and indirectly affect the epigenetic landscapes of individual loci.

### Speculative model for targeting RdDM to naïve loci

*A. thaliana* is one of the few exceptional plant species that contains not only SPT6L but also an additional SPT6 paralogue without the AGO-hook domain. Although SPT6 was detected in the Pol II complex (Antosz et al., 2017), nothing is known about its function. The presence of both SPT6 and SPT6L in this plant model was likely the reason for overlooking the fact that the only conserved homolog in *Viridiplantae* is SPT6L that carries the AGO-hook with one exception found, which is *Carica papaya*. This contrasts with SPT5 and SPT5L. These paralogues are regarded to be specialized for separate roles in the Pol II and Pol V complexes, respectively (Bies-Etheve et al., 2009). Different evolutionary trajectories can result from the fact that SPT5L, unlike SPT6L, evolved in the ancestor of vascular plants with already well-defined RdDM machinery. However, it is also possible that there was some evolutionary constraint on the subfunctionalization of SPT6L. Such a reason could be if one protein molecule had to physically interact first with one and then with the other polymerase complex. Chen et al., 2019 documented that SPT6L without SH2 domain (necessary for binding to both Pol II and Pol V loci; Fig. 1E) is still able to bind in gene promoters which indicates its ability to bind to chromatin independently of the Pol II complex. In the follow-up work, SPT6L was shown to participate in attracting chromatin remodeling complexes that open chromatin for initiation of Pol II transcription (Shu et al., 2022). Based on these findings, we speculate that SPT6L might play a similar role for the Pol V complex in *de novo* DNA methylation of naïve loci, where Pol V cannot be tethered by SUVH2/9 that do this at already methylated loci (Johnson et al., 2014). Recently published data by Sigman et al., 2021 indicated that Pol II can recruit AGO4 (and possibly other AGOs) to naïve loci that are targeted by sRNAs and that it might participate in Pol V attraction to these loci. We hypothesize that AGO4 is present at these loci together with SPT6L. They both may be deposited there together by Pol II whose nascent transcripts was just recognised by sRNA loaded on AGO4 attached to SPT6L in the Pol II complex. Thereafter, SPT6L can stay bound to histones in the locus and mediate chromatin re-opening also for Pol V transcription (Fig. 5B).

## Data availability statement

The authors confirm that the data supporting the findings of this study are available within the article [and/or] its supplementary materials. References to all previously published data analyzed in this study are listed in Supplemental Table S1.

## Conflict of Interest

The authors declare that the research was conducted in the absence of any commercial or financial relationships that could be construed as a potential conflict of interest.

## Acknowledgments

This work was supported by Charles University (project GA UK No. 382521) and Czech Science Foundation (project No. 24-12869S). Computational resources were provided by the e-INFRA CZ project (ID:90254), supported by the Ministry of Education, Youth and Sports of the Czech Republic.

## Author contributions

V.C. performed all the genomic data analyses. T.K. performed the phylogenetic analysis. V.C. and L.F. conceived the study and interpreted its results. All authors contributed to the manuscript writing. All authors have read and approved the final manuscript.

## Notes

### Competing Interest Statement

The authors have declared no competing interest.

## References

Antosz, W., Pfab, A., Ehrnsberger, H. F., Holzinger, P., Köllen, K., Mortensen, S. A., et al. (2017). The Composition of the Arabidopsis RNA Polymerase II Transcript Elongation Complex Reveals the Interplay Between Elongation and mRNA Processing Factors. Plant Cell, tpc.00735.2016. doi: 10.1105/tpc.16.00735.

Axtell, M. J. (2013). ShortStack: Comprehensive annotation and quantification of small RNA genes. RNA 19, 740–751. doi: 10.1261/rna.035279.112.

Basu, S., and Wallner, B. (2016). DockQ: A Quality Measure for Protein-Protein Docking Models. PLOS ONE 11, e0161879. doi: 10.1371/journal.pone.0161879.

Bies-Etheve, N., Pontier, D., Lahmy, S., Picart, C., Vega, D., Cooke, R., et al. (2009). RNA-directed DNA methylation requires an AGO4-interacting member of the SPT5 elongation factor family. EMBO Rep 10, 649–654. doi: 10.1038/embor.2009.31.

Blevins, T., Podicheti, R., Mishra, V., Marasco, M., Wang, J., Rusch, D., et al. (2015). Identification of Pol IV and RDR2-dependent precursors of 24 nt siRNAs guiding de novo DNA methylation in Arabidopsis. eLife Sciences 4, e09591. doi: 10.7554/eLife.09591.

Böhmdorfer, G., Sethuraman, S., Rowley, M. J., Krzyszton, M., Rothi, M. H., Bouzit, L., et al. (2016). Long non-coding RNA produced by RNA polymerase V determines boundaries of heterochromatin. eLife 5, e19092. doi: 10.7554/eLife.19092.

Čermák, V., Tyč, D., Přibylová, A., and Fischer, L. (2020). Unexpected variations in posttranscriptional gene silencing induced by differentially produced dsRNAs in tobacco cells. Biochimica et Biophysica Acta (BBA) - Gene Regulatory Mechanisms 1863, 194647. doi: 10.1016/j.bbagrm.2020.194647.

Chekulaeva, M., Mathys, H., Zipprich, J. T., Attig, J., Colic, M., Parker, R., et al. (2011). miRNA repression involves GW182-mediated recruitment of CCR4–NOT through conserved W-containing motifs. Nat Struct Mol Biol 18, 1218–1226. doi: 10.1038/nsmb.2166.

Chen, C., Shu, J., Li, C., Thapa, R. K., Nguyen, V., Yu, K., et al. (2019). RNA polymerase II-independent recruitment of SPT6L at transcription start sites in Arabidopsis. Nucleic Acids Res. doi: 10.1093/nar/gkz465.

Cuerda-Gil, D., and Slotkin, R. K. (2016). Non-canonical RNA-directed DNA methylation. Nature Plants 2, 16163. doi: 10.1038/nplants.2016.163.

Danecek, P., Bonfield, J. K., Liddle, J., Marshall, J., Ohan, V., Pollard, M. O., et al. (2021). Twelve years of SAMtools and BCFtools. GigaScience 10, giab008. doi: 10.1093/gigascience/giab008.

Daxinger, L., Kanno, T., Bucher, E., van der Winden, J., Naumann, U., Matzke, A. J. M., et al. (2009). A stepwise pathway for biogenesis of 24-nt secondary siRNAs and spreading of DNA methylation. EMBO J 28, 48–57. doi: 10.1038/emboj.2008.260.

El-Shami, M., Pontier, D., Lahmy, S., Braun, L., Picart, C., Vega, D., et al. (2007). Reiterated WG/GW motifs form functionally and evolutionarily conserved ARGONAUTE-binding platforms in RNAi-related components. Genes & Development 21, 2539– 2544. doi: 10.1101/gad.451207.

Engel, K. L., French, S. L., Viktorovskaya, O. V., Beyer, A. L., and Schneider, D. A. (2015). Spt6 Is Essential for rRNA Synthesis by RNA Polymerase I. Molecular and Cellular Biology 35, 2321–2331. doi: 10.1128/MCB.01499-14.

Evans, R., O’Neill, M., Pritzel, A., Antropova, N., Senior, A., Green, T., et al. (2022). Protein complex prediction with AlphaFold-Multimer. 2021.10.04.463034. doi: 10.1101/2021.10.04.463034.

Fultz, D., and Slotkin, R. K. (2017). Exogenous Transposable Elements Circumvent Identity-Based Silencing Permitting the Dissection of Expression Dependent Silencing. Plant Cell, tpc.00718.2016. doi: 10.1105/tpc.16.00718.

Gallego-Bartolomé, J., Liu, W., Kuo, P. H., Feng, S., Ghoshal, B., Gardiner, J., et al. (2019). Co-targeting RNA Polymerases IV and V Promotes Efficient De Novo DNA Methylation in Arabidopsis. Cell. doi: 10.1016/j.cell.2019.01.029.

Garcia, D., Garcia, S., Pontier, D., Marchais, A., Renou, J. P., Lagrange, T., et al. (2012). Ago Hook and RNA Helicase Motifs Underpin Dual Roles for SDE3 in Antiviral Defense and Silencing of Nonconserved Intergenic Regions. Molecular Cell 48, 109–120. doi: 10.1016/j.molcel.2012.07.028.

Goll, M. G., and Bestor, T. H. (2005). Eukaryotic Cytosine Methyltransferases. Annual Review of Biochemistry 74, 481–514. doi: 10.1146/annurev.biochem.74.010904.153721.

Havecker, E. R., Wallbridge, L. M., Hardcastle, T. J., Bush, M. S., Kelly, K. A., Dunn, R. M., et al. (2010). The Arabidopsis RNA-Directed DNA Methylation Argonautes Functionally Diverge Based on Their Expression and Interaction with Target Loci. The Plant Cell Online 22, 321–334. doi: 10.1105/tpc.109.072199.

He, L., Huang, H., Bradai, M., Zhao, C., You, Y., Ma, J., et al. (2022). DNA methylation-free Arabidopsis reveals crucial roles of DNA methylation in regulating gene expression and development. Nat Commun 13, 1335. doi: 10.1038/s41467-022-28940-2.

He, L., Zhao, C., Zhang, Q., Zinta, G., Wang, D., Lozano-Durán, R., et al. (2021). Pathway conversion enables a double-lock mechanism to maintain DNA methylation and genome stability. PNAS 118. doi: 10.1073/pnas.2107320118.

Huang, Y., Kendall, T., Forsythe, E. S., Dorantes-Acosta, A., Li, S., Caballero-Perez, J., et al. (2015). Ancient origin and recent innovations of RNA Polymerase IV and V. *Mol Biol Evol*, msv060. doi: 10.1093/molbev/msv060.

Johnson, L. M., Du, J., Hale, C. J., Bischof, S., Feng, S., Chodavarapu, R. K., et al. (2014). SRA- and SET-domain-containing proteins link RNA polymerase V occupancy to DNA methylation. Nature 507, 124–128. doi: 10.1038/nature12931.

Karlowski, W. M., Zielezinski, A., Carrère, J., Pontier, D., Lagrange, T., and Cooke, R. (2010). Genome-wide computational identification of WG/GW Argonaute-binding proteins in Arabidopsis. Nucl. Acids Res. 38, 4231–4245. doi: 10.1093/nar/gkq162.

Köllen, K., Dietz, L., Bies-Etheve, N., Lagrange, T., Grasser, M., and Grasser, K. D. (2015). The zinc-finger protein SPT4 interacts with SPT5L/KTF1 and modulates transcriptional silencing in Arabidopsis. FEBS Letters. doi: 10.1016/j.febslet.2015.09.017.

Kondili, M., Fust, A., Preussner, J., Kuenne, C., Braun, T., and Looso, M. (2017). UROPA: a tool for Universal RObust Peak Annotation. Sci Rep 7, 2593. doi: 10.1038/s41598-017-02464-y.

Laflamme, G., and Mekhail, K. (2020). Biomolecular condensates as arbiters of biochemical reactions inside the nucleus. Commun Biol 3, 1–8. doi: 10.1038/s42003-020-01517-9.

Lahmy, S., Pontier, D., Bies-Etheve, N., Laudié, M., Feng, S., Jobet, E., et al. (2016). Evidence for ARGONAUTE4–DNA interactions in RNA-directed DNA methylation in plants. Genes Dev. 30, 2565–2570. doi: 10.1101/gad.289553.116.

Langmead, B., and Salzberg, S. L. (2012). Fast gapped-read alignment with Bowtie 2. Nat Methods 9, 357–359. doi: 10.1038/nmeth.1923.

Leliaert, F., Smith, D. R., Moreau, H., Herron, M. D., Verbruggen, H., Delwiche, C. F., et al. (2012). Phylogeny and Molecular Evolution of the Green Algae. Critical Reviews in Plant Sciences 31, 1–46. doi: 10.1080/07352689.2011.615705.

Li, Y., Yuan, Y., Fang, X., Lu, X., Lian, B., Zhao, G., et al. (2018). A Role for MINIYO and QUATRE-QUART2 in the Assembly of RNA Polymerases II, IV, and V in Arabidopsis. The Plant Cell 30, 466–480. doi: 10.1105/tpc.17.00380.

Liang, Z., Geng, Y., Ji, C., Du, H., Wong, C. E., Zhang, Q., et al. (2020). Mesostigma viride Genome and Transcriptome Provide Insights into the Origin and Evolution of Streptophyta. Advanced Science 7, 1901850. doi: 10.1002/advs.201901850.

Liu, W., Duttke, S. H., Hetzel, J., Groth, M., Feng, S., Gallego-Bartolome, J., et al. (2018). RNA-directed DNA methylation involves co-transcriptional small-RNA-guided slicing of polymerase V transcripts in Arabidopsis. Nature Plants, 1. doi: 10.1038/s41477-017-0100-y.

Liu, Y., Shu, J., Zhang, Z., Ding, N., Liu, J., Liu, J., et al. (2024). A conserved Pol II elongator SPT6L mediates Pol V transcription elongation to regulate RNA-directed DNA methylation in Arabidopsis. 2024.01.09.574790. doi: 10.1101/2024.01.09.574790.

Luo, C., Sidote, D. J., Zhang, Y., Kerstetter, R. A., Michael, T. P., and Lam, E. (2013). Integrative analysis of chromatin states in Arabidopsis identified potential regulatory mechanisms for natural antisense transcript production. The Plant Journal 73, 77–90. doi: 10.1111/tpj.12017.

Luo, J., and Hall, B. (2007). A Multistep Process Gave Rise to RNA Polymerase IV of Land Plants. Journal of Molecular Evolution 64, 101–112. doi: 10.1007/s00239-006-0093-z.

Ma, Z., Castillo-González, C., Wang, Z., Sun, D., Hu, X., Shen, X., et al. (2018). Arabidopsis Serrate Coordinates Histone Methyltransferases ATXR5/6 and RNA Processing Factor RDR6 to Regulate Transposon Expression. Developmental Cell 45, 769–784.e6. doi: 10.1016/j.devcel.2018.05.023.

Matzke, M. A., Kanno, T., and Matzke, A. J. M. (2015). RNA-Directed DNA Methylation: The Evolution of a Complex Epigenetic Pathway in Flowering Plants. Annual Review of Plant Biology 66, null. doi: 10.1146/annurev-arplant-043014-114633.

Mi, S., Cai, T., Hu, Y., Chen, Y., Hodges, E., Ni, F., et al. (2008). Sorting of Small RNAs into Arabidopsis Argonaute Complexes Is Directed by the 5’ Terminal Nucleotide. Cell 133, 116–127. doi: 10.1016/j.cell.2008.02.034.

Onodera, Y., Haag, J. R., Ream, T., Nunes, P. C., Pontes, O., and Pikaard, C. S. (2005). Plant Nuclear RNA Polymerase IV Mediates siRNA and DNA Methylation-Dependent Heterochromatin Formation. Cell 120, 613–622. doi: 10.1016/j.cell.2005.02.007.

Panda, K., Ji, L., Neumann, D. A., Daron, J., Schmitz, R. J., and Slotkin, R. K. (2016). Full-length autonomous transposable elements are preferentially targeted by expression-dependent forms of RNA-directed DNA methylation. Genome Biology 17, 170. doi: 10.1186/s13059-016-1032-y.

Pettersen, E. F., Goddard, T. D., Huang, C. C., Meng, E. C., Couch, G. S., Croll, T. I., et al. (2021). UCSF ChimeraX: Structure visualization for researchers, educators, and developers. Protein Science 30, 70–82. doi: 10.1002/pro.3943.

Pontier, D., Picart, C., Roudier, F., Garcia, D., Lahmy, S., Azevedo, J., et al. (2012). NERD, a Plant-Specific GW Protein, Defines an Additional RNAi-Dependent Chromatin-Based Pathway in Arabidopsis. Molecular Cell 48, 121–132. doi: 10.1016/j.molcel.2012.07.027.

Quinlan, A. R., and Hall, I. M. (2010). BEDTools: a flexible suite of utilities for comparing genomic features. Bioinformatics 26, 841–842. doi: 10.1093/bioinformatics/btq033.

Ramírez, F., Ryan, D. P., Grüning, B., Bhardwaj, V., Kilpert, F., Richter, A. S., et al. (2016). deepTools2: a next generation web server for deep-sequencing data analysis. Nucleic Acids Res 44, W160–W165. doi: 10.1093/nar/gkw257.

Sdano, M. A., Fulcher, J. M., Palani, S., Chandrasekharan, M. B., Parnell, T. J., Whitby, F. G., et al. (2017). A novel SH2 recognition mechanism recruits Spt6 to the doubly phosphorylated RNA polymerase II linker at sites of transcription. eLife 6, e28723. doi: 10.7554/eLife.28723.

Sheu-Gruttadauria, J., and MacRae, I. J. (2018). Phase Transitions in the Assembly and Function of Human miRISC. Cell 173, 946–957.e16. doi: 10.1016/j.cell.2018.02.051.

Shu, J., Chen, C., Thapa, R. K., Bian, S., Nguyen, V., Yu, K., et al. (2019). Genome-wide occupancy of histone H3K27 methyltransferases CURLY LEAF and SWINGER in Arabidopsis seedlings. Plant Direct 3, e00100. doi: 10.1002/pld3.100.

Shu, J., Ding, N., Liu, J., Cui, Y., and Chen, C. (2022). Transcription elongator SPT6L regulates the occupancies of the SWI2/SNF2 chromatin remodelers SYD/BRM and nucleosomes at transcription start sites in Arabidopsis. *Nucleic Acids Research*, gkac1126. doi: 10.1093/nar/gkac1126.

Sigman, M. J., Panda, K., Kirchner, R., McLain, L. L., Payne, H., Peasari, J. R., et al. (2021). An siRNA-guided ARGONAUTE protein directs RNA polymerase V to initiate DNA methylation. Nature Plants 7, 1461–1474. doi: 10.1038/s41477-021-01008-7.

Stamatakis, A. (2014). RAxML version 8: a tool for phylogenetic analysis and post-analysis of large phylogenies. Bioinformatics 30, 1312–1313. doi: 10.1093/bioinformatics/btu033.

Stroud, H., Greenberg, M. V. C., Feng, S., Bernatavichute, Y. V., and Jacobsen, S. E. (2013). Comprehensive Analysis of Silencing Mutants Reveals Complex Regulation of the Arabidopsis Methylome. Cell 152, 352–364. doi: 10.1016/j.cell.2012.10.054.

Trujillo, J. T. (2019). The Origin and Evolution of Plant-Specific RNA Polymerases and Genes Involved in RNA-Directed DNA Methylation. Available at: https://repository.arizona.edu/handle/10150/632985 [Accessed September 18, 2023].

Tsuzuki, M., Sethuraman, S., Coke, A. N., Rothi, M. H., Boyle, A. P., and Wierzbicki, A. T. (2020). Broad noncoding transcription suggests genome surveillance by RNA polymerase V. PNAS. doi: 10.1073/pnas.2014419117.

Vos, S. M., Farnung, L., Boehning, M., Wigge, C., Linden, A., Urlaub, H., et al. (2018). Structure of activated transcription complex Pol II–DSIF–PAF–SPT6. Nature 560, 607–612. doi: 10.1038/s41586-018-0440-4.

Wang, Y., and Ma, H. (2015). Step-wise and lineage-specific diversification of plant RNA polymerase genes and origin of the largest plant-specific subunits. New Phytol 207, 1198–1212. doi: 10.1111/nph.13432.

Wierzbicki, A. T., Haag, J. R., and Pikaard, C. S. (2008). Noncoding Transcription by RNA Polymerase Pol IVb/Pol V Mediates Transcriptional Silencing of Overlapping and Adjacent Genes. Cell 135, 635–648. doi: 10.1016/j.cell.2008.09.035.

Yaari, R., Katz, A., Domb, K., Harris, K. D., Zemach, A., and Ohad, N. (2019). RdDM - independent de novo and heterochromatin DNA methylation by plant CMT and DNMT3 orthologs. Nat Commun 10, 1–10. doi: 10.1038/s41467-019-09496-0.

Yang, L., Wu, G., and Poethig, R. S. (2012). Mutations in the GW-repeat protein SUO reveal a developmental function for microRNA-mediated translational repression in Arabidopsis. PNAS 109, 315–320. doi: 10.1073/pnas.1114673109.

Yang, R., Zheng, Z., Chen, Q., Yang, L., Huang, H., Miki, D., et al. (2017). The developmental regulator PKL is required to maintain correct DNA methylation patterns at RNA-directed DNA methylation loci. Genome Biol 18. doi: 10.1186/s13059-017-1226-y.

Yelagandula, R., Stroud, H., Holec, S., Zhou, K., Feng, S., Zhong, X., et al. (2014). The Histone Variant H2A.W Defines Heterochromatin and Promotes Chromatin Condensation in Arabidopsis. Cell 158, 98–109. doi: 10.1016/j.cell.2014.06.006.

Zhang, H., Lang, Z., and Zhu, J.-K. (2018a). Dynamics and function of DNA methylation in plants. Nature Reviews Molecular Cell Biology 19, 489. doi: 10.1038/s41580-018-0016-z.

Zhang, Y., Harris, C. J., Liu, Q., Liu, W., Ausin, I., Long, Y., et al. (2018b). Large-scale comparative epigenomics reveals hierarchical regulation of non-CG methylation in Arabidopsis. PNAS, 201716300. doi: 10.1073/pnas.1716300115.

Zhang, Z., Liu, X., Guo, X., Wang, X.-J., and Zhang, X. (2016). Arabidopsis AGO3 predominantly recruits 24-nt small RNAs to regulate epigenetic silencing. Nature Plants 2, nplants201649. doi: 10.1038/nplants.2016.49.

Zheng, Q., Jordan Rowley, M., Böhmdorfer, G., Sandhu, D., Gregory, B. D., and Wierzbicki, A. T. (2012). RNA Polymerase V targets transcriptional silencing components to promoters of protein-coding genes. The Plant Journal 73, 179–189. doi: 10.1111/tpj.12034.

Zhong, X., Du, J., Hale, C. J., Gallego-Bartolome, J., Feng, S., Vashisht, A. A., et al. (2014). Molecular Mechanism of Action of Plant DRM De Novo DNA Methyltransferases. Cell 157, 1050–1060. doi: 10.1016/j.cell.2014.03.056.

Zhong, Z., Feng, S., Duttke, S. H., Potok, M. E., Zhang, Y., Gallego-Bartolomé, J., et al. (2021). DNA methylation-linked chromatin accessibility affects genomic architecture in Arabidopsis. PNAS 118. doi: 10.1073/pnas.2023347118.

Zhou, J.-X., Du, P., Liu, Z.-W., Feng, C., Cai, X.-W., and He, X.-J. (2021). FVE promotes RNA-directed DNA methylation by facilitating the association of RNA polymerase V with chromatin. The Plant Journal 107, 467–479. doi: 10.1111/tpj.15302.

Zielezinski, A., and Karlowski, W. M. (2015). Integrative data analysis indicates an intrinsic disordered domain character of Argonaute-binding motifs. Bioinformatics 31, 332–339. doi: 10.1093/bioinformatics/btu666.

