## Supplemental_Figures for "SPT6L, a newly discovered ancestral component of the plant RNA-directed DNA methylation pathway"

### Supplemental materials

#### Supplemental Figures

**Supplemental Fig. S1**: SPT6L is part of the Pol V complex

**Supplemental Fig. S2**: Characterization of Pol V loci occupied by SPT6L

**Supplemental Fig. S3**: Mode of DNA methylation, types of sRNAs and genomic elements at Pol V loci occupied by SPT6L

**Supplemental Fig S4**: Evolution of SPT6L, other relevant AGO-hook proteins and DNA methyltransferases in *Viridiplantae*

#### Supplemental Tables

**Supplemental Table S1**: Datasets used in this study

**Supplemental Table S2**: AlphaFold-Multimer (AFM) prediction scores (ipTM) for complex formation between SPT6L and relevant Pol II subunits and SPT6L and all NRPB7 paralogues. Scores for all five models generated by AFM are shown. List of used UniProt chain IDs. It should be noted that the two RdDM-related paralogues of NRPB7 are annotated in the TAIR10/Araport11 as NRPD7 and NRPE7, but NRPE7 is present in both Pol IV and Pol V complexes (Law et al. 2011) and more importantly their diversification is specific for the *Brassicaceae* family, other species have only one NRPD/E7 protein (Wang and Ma, 2015).

**Supplemental Table S3**: Table with underlying data for Fig. 3A,B and Supplemental Fig. S3A,B.

**Supplemental Table S4**: Table with underlying data for Fig. 3C and Supplemental Fig. S3C.

#### Supplemental Fig. S1: SPT6L is part of the Pol V complex

(A) Three representative genomic regions with SPT6L ChIP-seq and sRNA-seq data with indicated loci which are significantly enriched for these features and categorisation of these loci based on their overlaps. Pol V ChIP-seq data are shown to illustrate its coverage at the SPT6L and sRNA loci. (B) Boxplots with z-score of ChIP-seq coverage for various chromatin marks at loci presented in Fig. 1A. (C) Metaplots showing the ChIP-seq data of selected chromatin marks, WGBS data and ATAC-seq data (all in WT) for loci presented in Fig. 1A (± 1 kbp of adjacent regions). Randomized SPT6L∩sRNA loci are used as background. This is extended characterization of data from Fig. 1B and Supplemental Fig. S1B. (D) Clustered heatmap for SPT6L∩sRNA loci of data presented in Fig. 1B and Supplemental Fig. S1B (z-score is plotted in the graph). (E) Metaplots showing the ChIP-seq data of several RdDM proteins and Pol II for loci presented in Fig. 1A (± 1 kbp of adjacent regions). Randomized SPT6L∩sRNA loci are used as background. This is extended characterization of data from Fig. 1C. (E) Clustered heatmap for SPT6L∩sRNA loci of data presented in Fig. 1C (z-score is plotted in the graph). (G) Metaplot showing ChIP-seq data for SPT6L at Pol IV loci with overlap with Pol V loci (Pol IV∩Pol V) or without it (Pol IV(-)Pol V) and Pol V loci without overlap with Pol IV loci (Pol V(-)Pol IV). Randomized Pol IV∩Pol V loci are used as background. (H) Venn diagram showing overlaps between Pol IV, Pol V and SPT6L loci. (I) Overlay of the AlphaFold-Multimer predictions for the SPT6LxNRPB7 (light blue/green) and SPT6LxNRPE7 (dark blue/orange) complexes. Note that the SH2 domain of SPT6L is connected with a flexible linker to the rest of the SPT6L and therefore its position is not fixed in the structure, as can be seen also from Fig. 1F. The unstructured parts of SPT6L are not shown (300 aa residues from N-terminus and 210 aa residues from C-terminus).

Fig. S1A

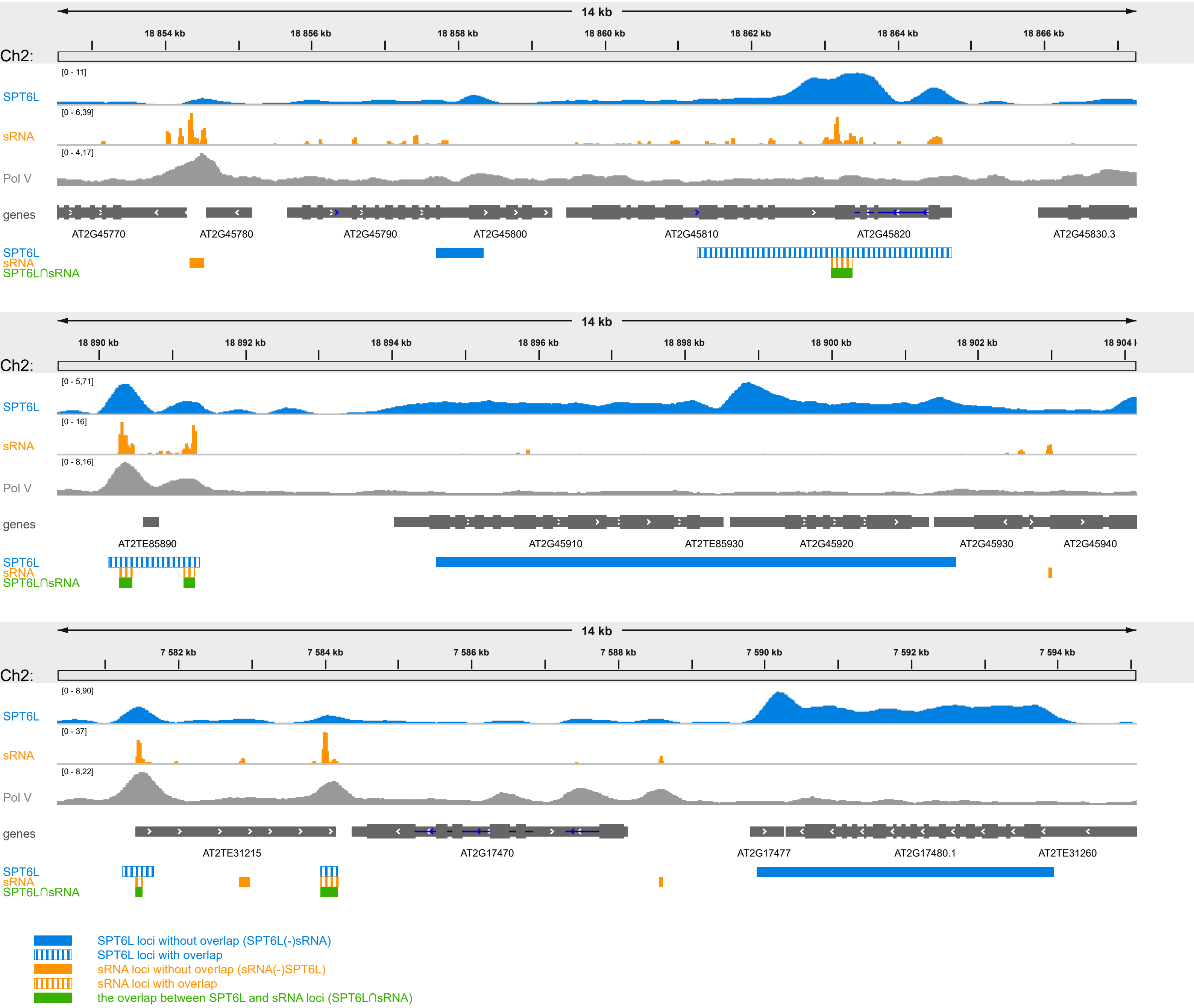


Fig. S1B

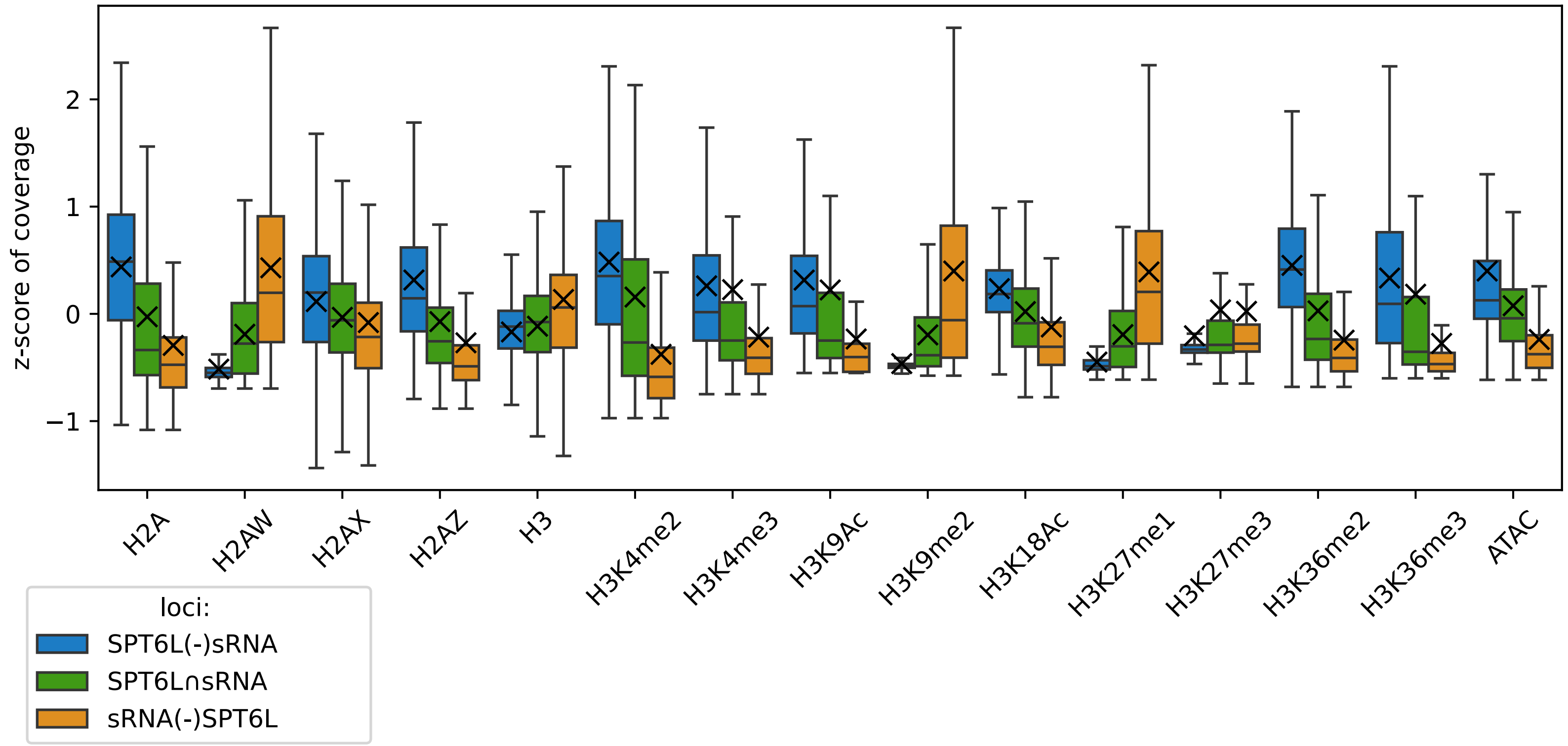

Fig. S1C

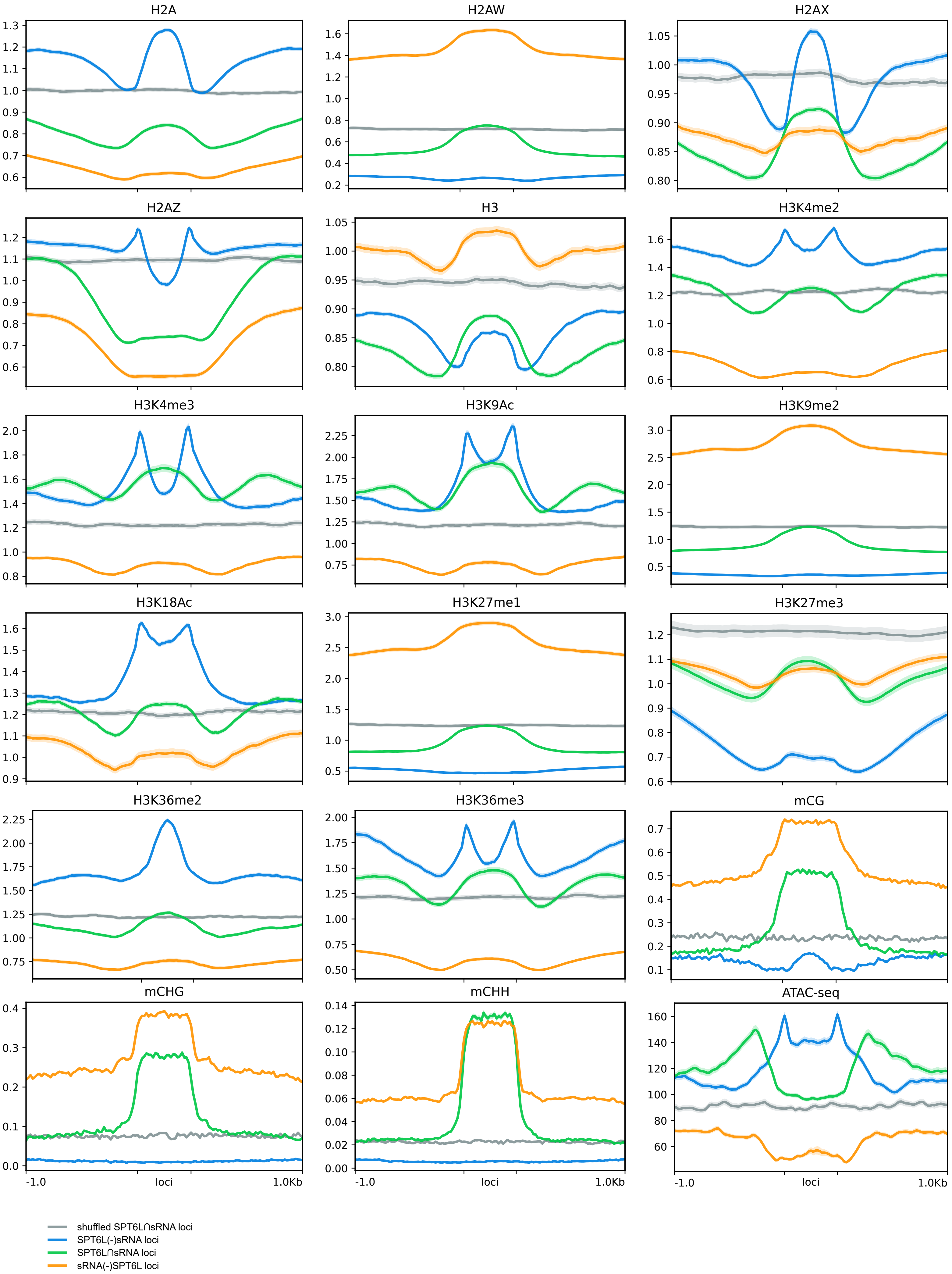
H2A
H2AW
H2AX
H2AZ
H3
H3K4me2
H3K4me3
H3K9Ac
H3K9me2
H3K18Ac
H3K27me1
H3K27me3
H3K36me2
H3K36me3
mCG
mCHG
mCHH
ATAC-seq
-1.0
loci
1.0Kb
shuffled SPT6L∩sRNA loci
SPT6L(-)sRNA loci
SPT6L∩sRNA loci
sRNA(-)SPT6L loci

Fig. S1D

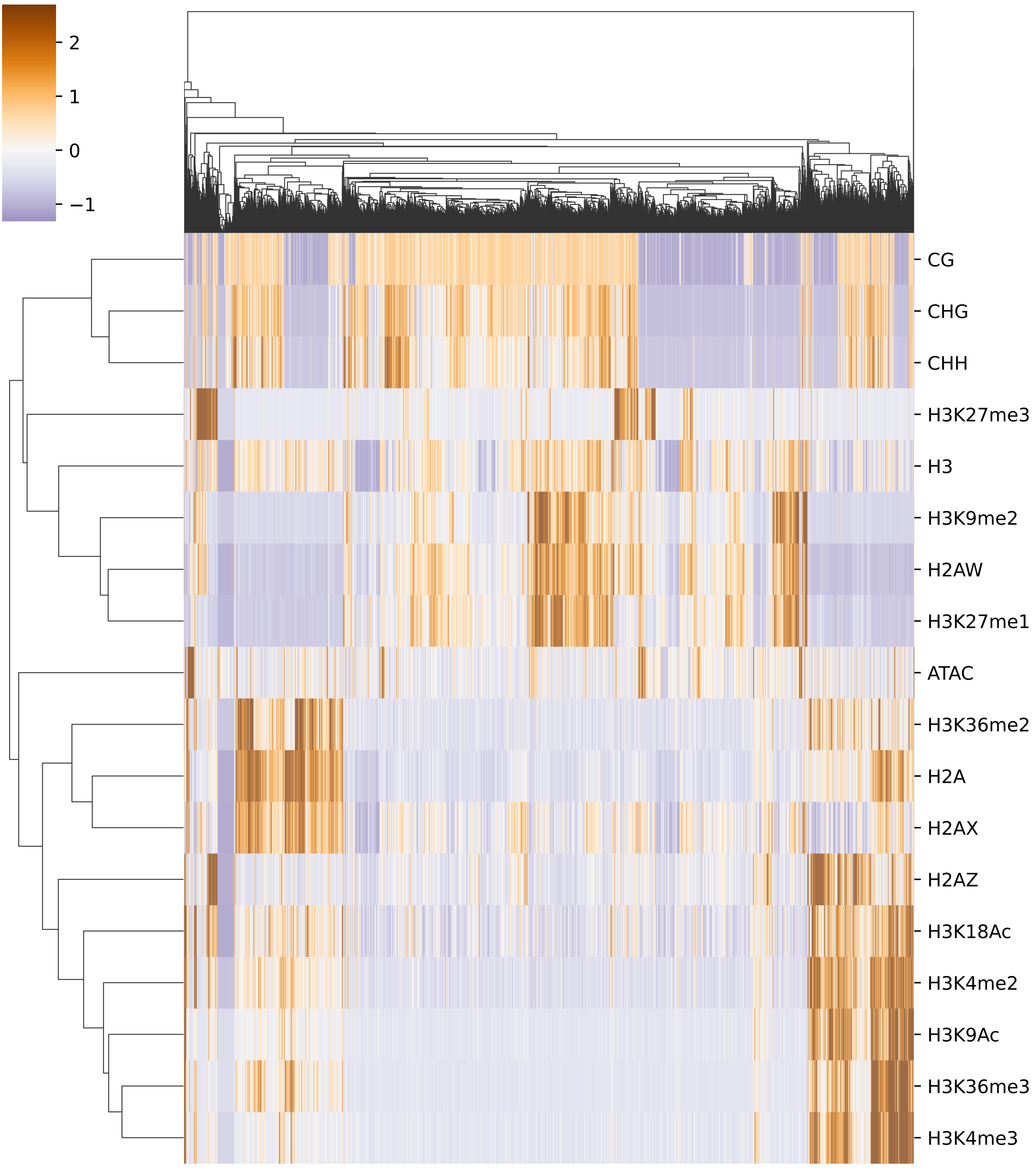
2
1
0
−1
CG
CHG
CHH
H3K27me3
H3
H3K9me2
H2AW
H3K27me1
ATAC
H3K36me2
H2A
H2AX
H2AZ
H3K18Ac
H3K4me2
H3K9Ac
H3K36me3
H3K4me3

Fig. S1E

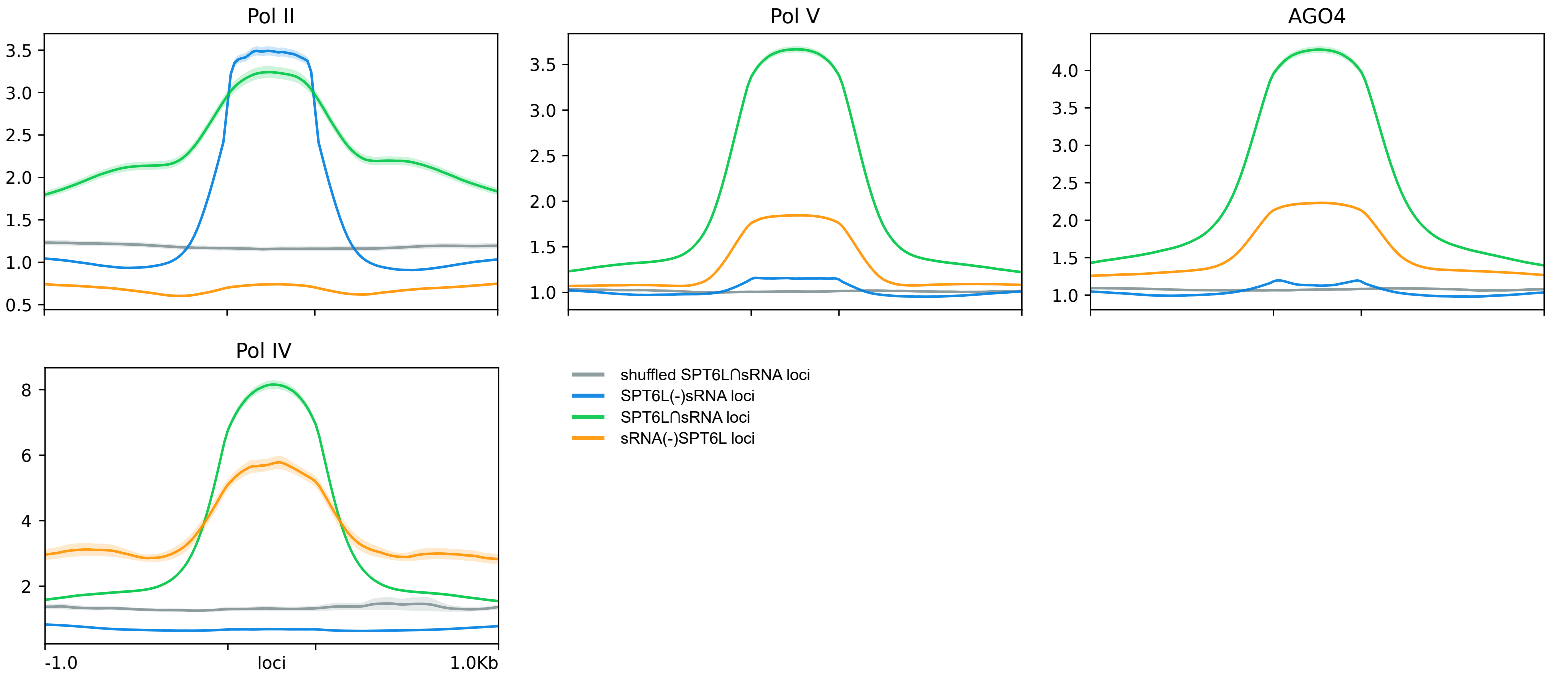


Fig. S1F

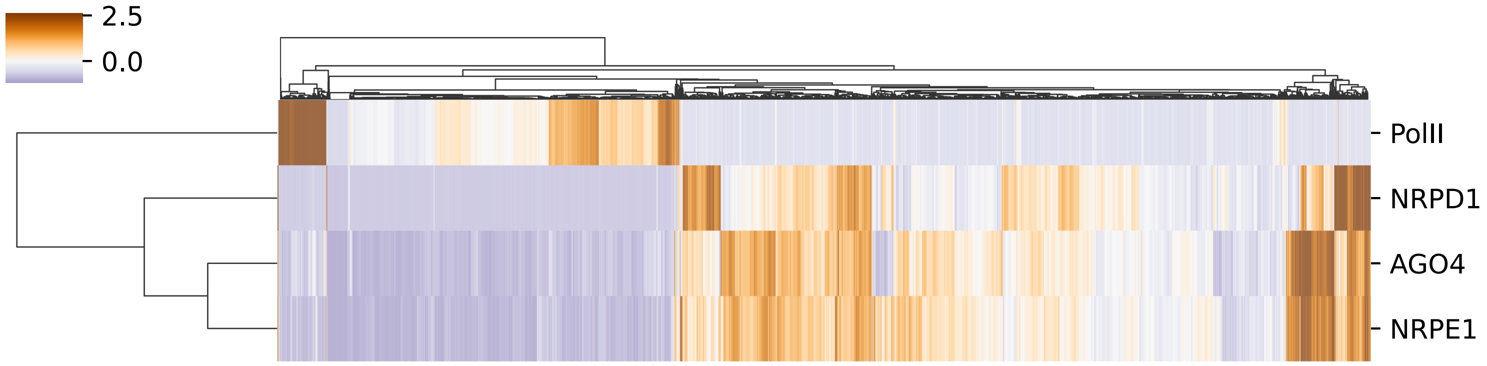


Fig. S1G

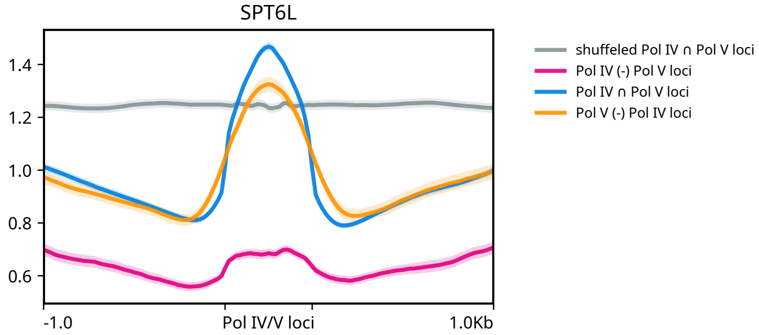


Fig. S1H

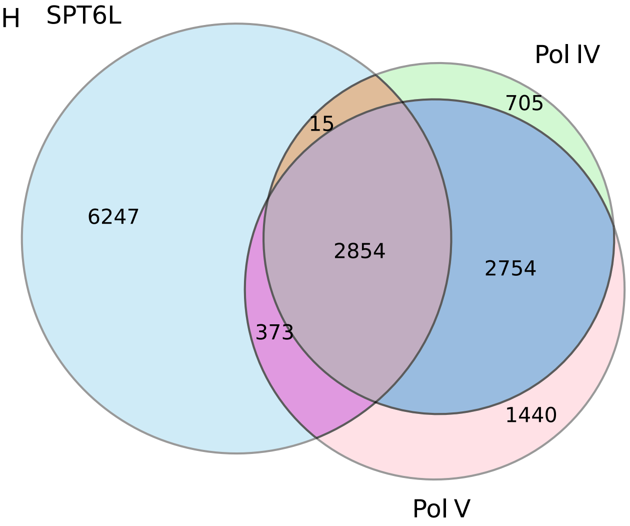


Fig. S1I

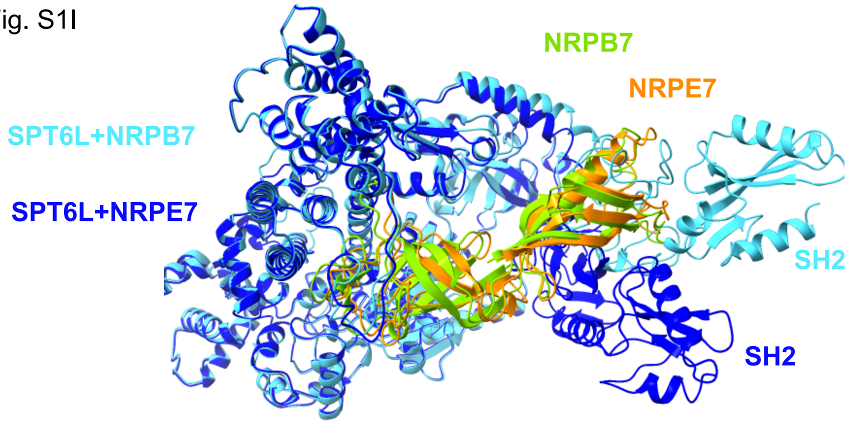

#### Supplemental Fig. S2: Characterization of Pol V loci occupied by SPT6L

(A) Distribution of Pol II, SPT6L, Pol V and subcategories of Pol V loci along chromosomes of *A. thaliana*. (B) Three representative genomic regions with Pol II (NRPB1), SPT6L and Pol V (NRPE1) ChIP-seq data with indicated loci which are significantly enriched for these proteins. Pol V loci are categorized according to Fig. 2A. (C,D) Extended version of Fig. 2 to show how the analyzed loci are affected by Pol II. (C) Boxplots showing the lengths and GC content of the four subcategories of Pol V loci. Note generally lower GC content at Pol V loci than at Pol II loci that are shown for comparison. (D) Enrichment/depletion of chromatin marks and RdDM components at P5(+)SPT6L loci calculated as log2(P5(+)SPT6L/P5(-)SPT6L) and P5(+)SPT6L(-)P2 loci calculated as log2(P5(+)SPT6L(-)P2/P5(-)SPT6L(-)P2). (E,F) Metaplots showing ChIP-seq data (underlying data for Fig. 2C,E) for (E) various chromatin marks and (F) RdDM components at the four subcategories of Pol V loci. Metaplots contain ± 1 kbp of adjacent regions, randomized P5 loci are used for background. Significance is indicated by asterisks: $p < 0.05$ = *; $p < 0.005$ = **; $p < 0.0005$ = ***; NS = not significant; in Supplemental Fig. S2D the statistics were calculated from the normalized coverages at the P5(+)SPT6L and P5(-)SPT6L or P5(+)SPT6L(-)P2 and P5(-)SPT6L(-)P2 loci from which the log2 ratio was calculated.

Fig. S2A

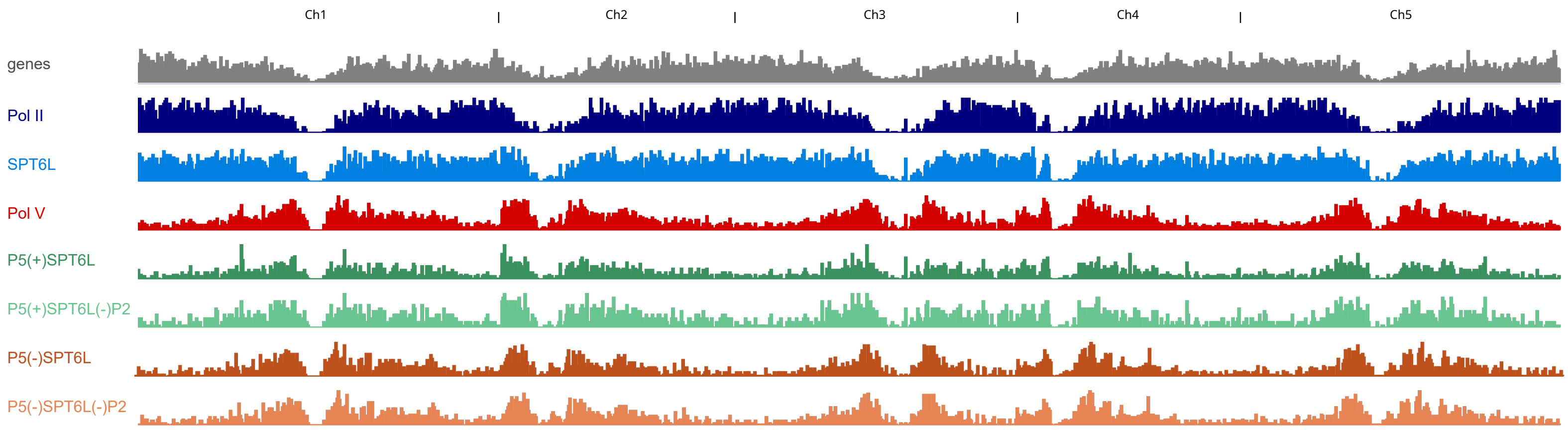


Fig. S2B

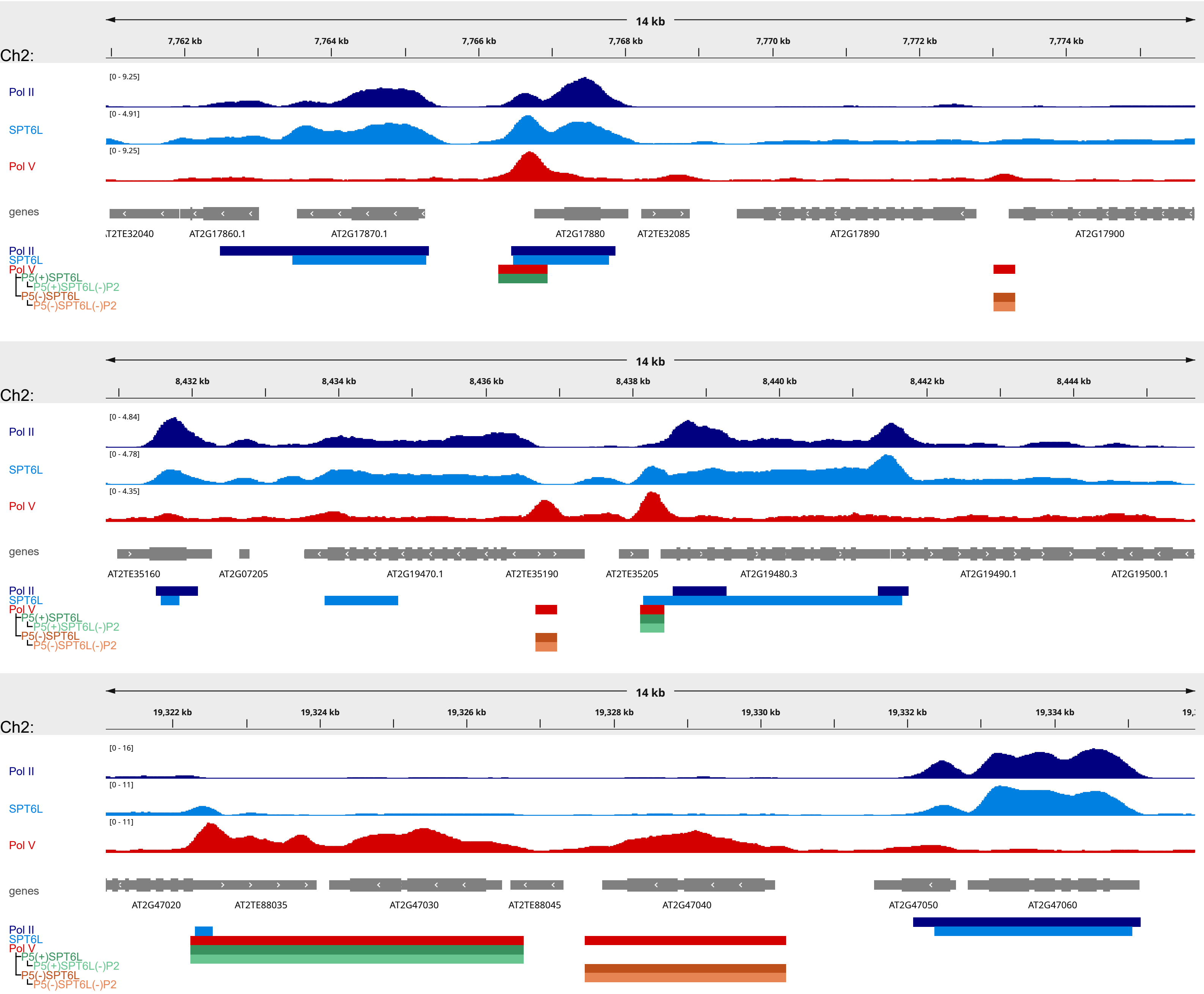

Fig. S2C

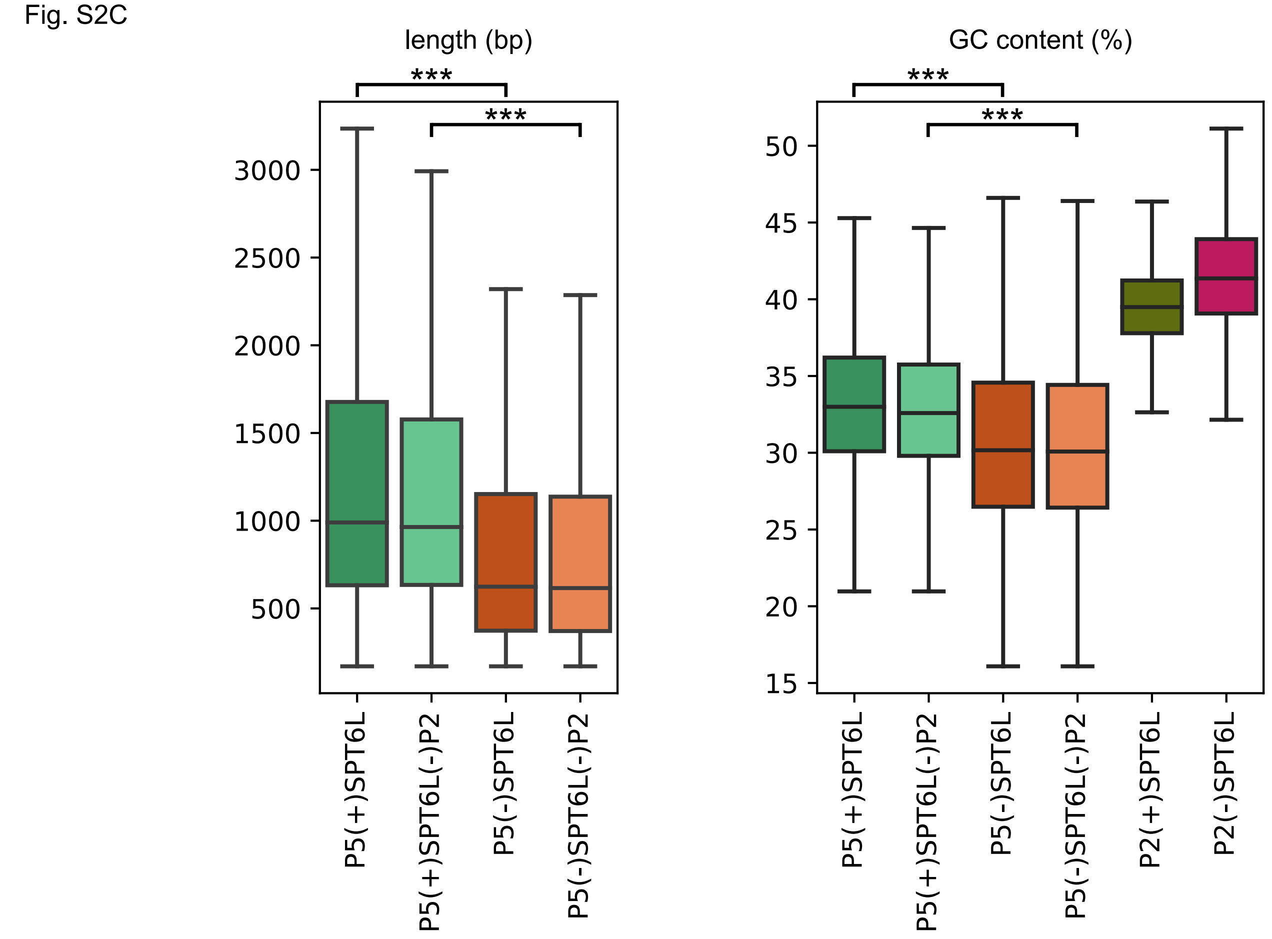


Fig. S2D

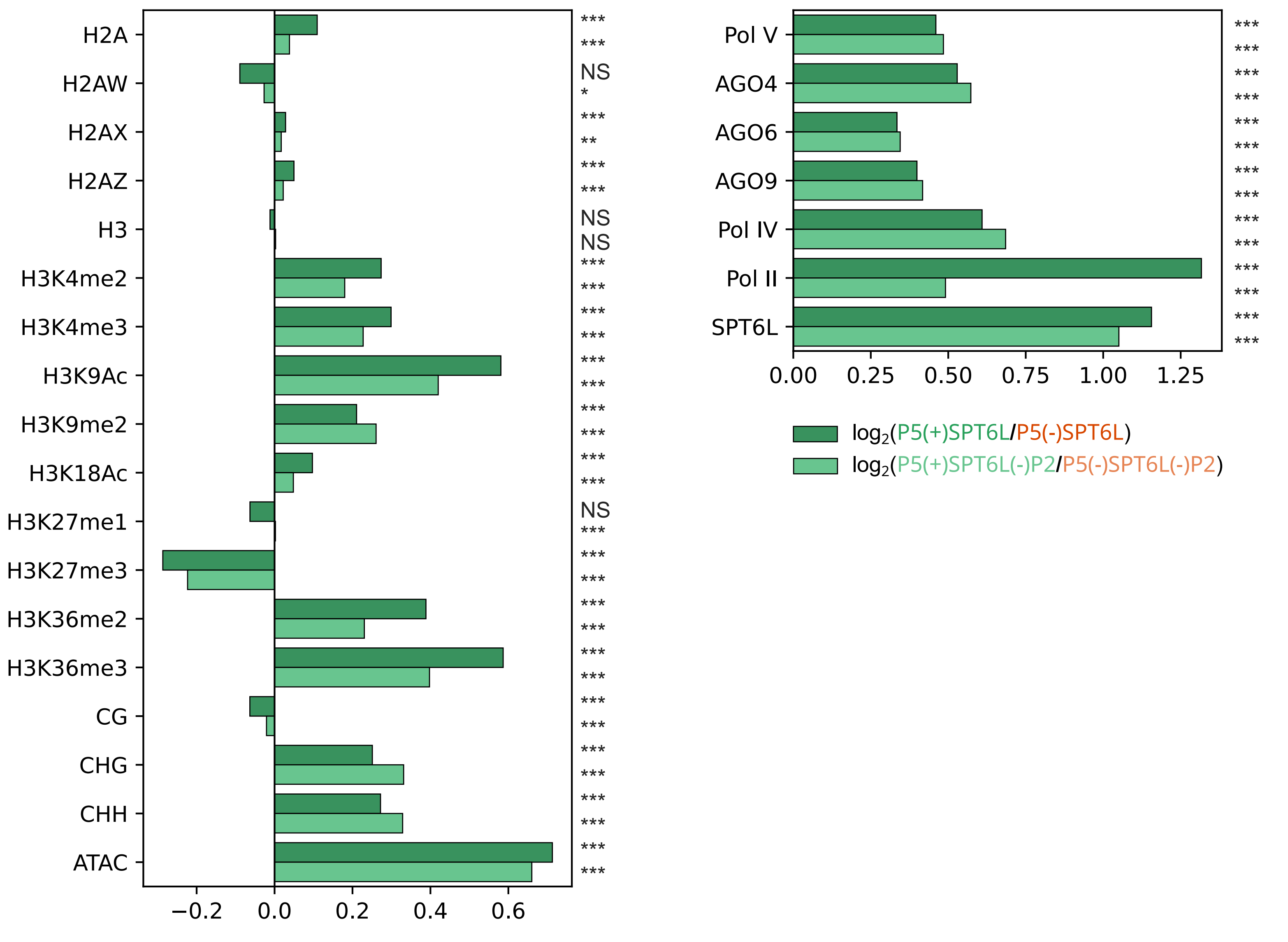

Fig. S2E

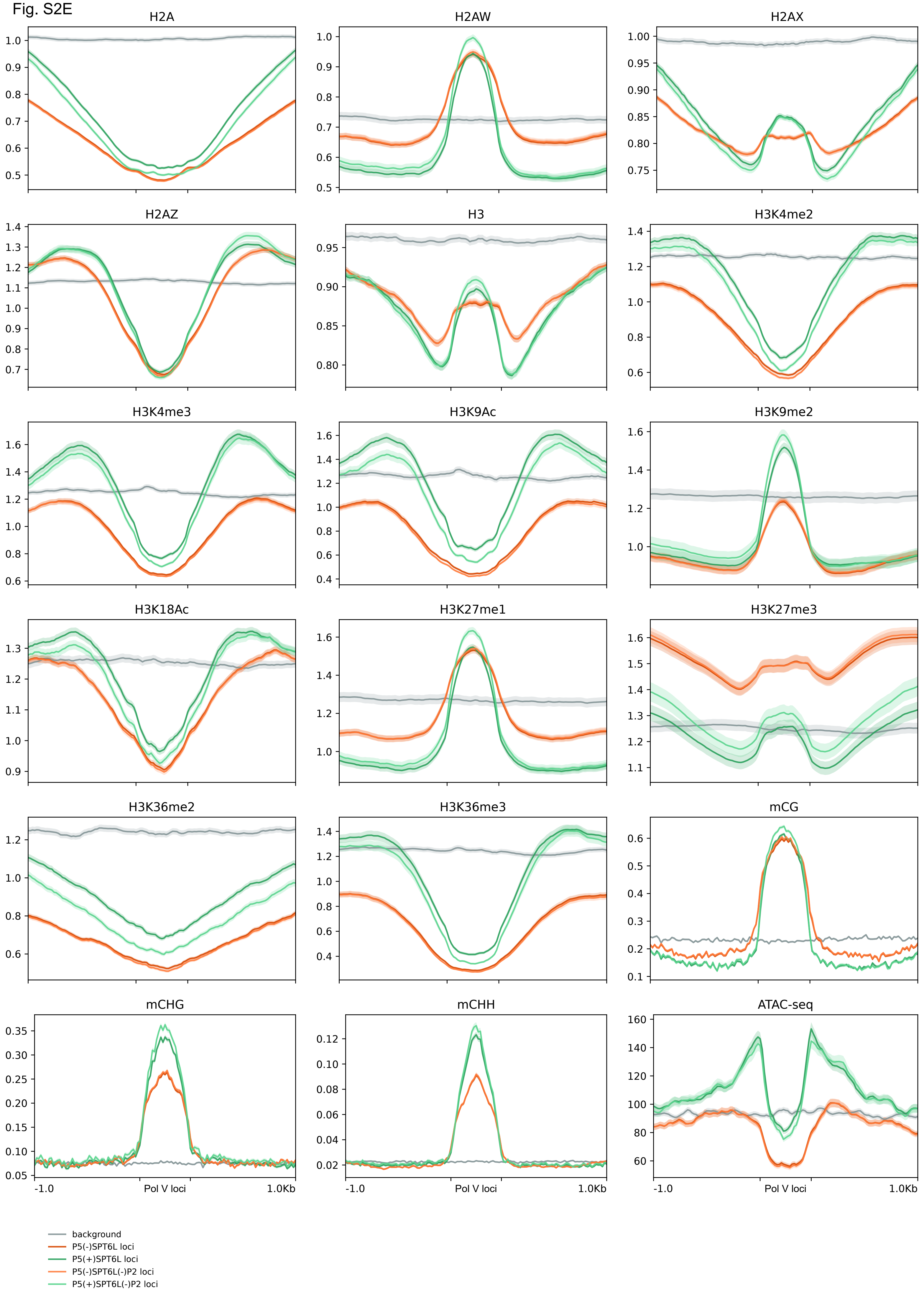
H2A
H2AW
H2AX
H2AZ
H3
H3K4me2
H3K4me3
H3K9Ac
H3K9me2
H3K18Ac
H3K27me1
H3K27me3
H3K36me2
H3K36me3
mCG
mCHG
mCHH
ATAC-seq
-1.0
Pol V loci
1.0Kb
background
P5(-)SPT6L loci
P5(+)SPT6L loci
P5(-)SPT6L(-)P2 loci
P5(+)SPT6L(-)P2 loci

Fig. S2F

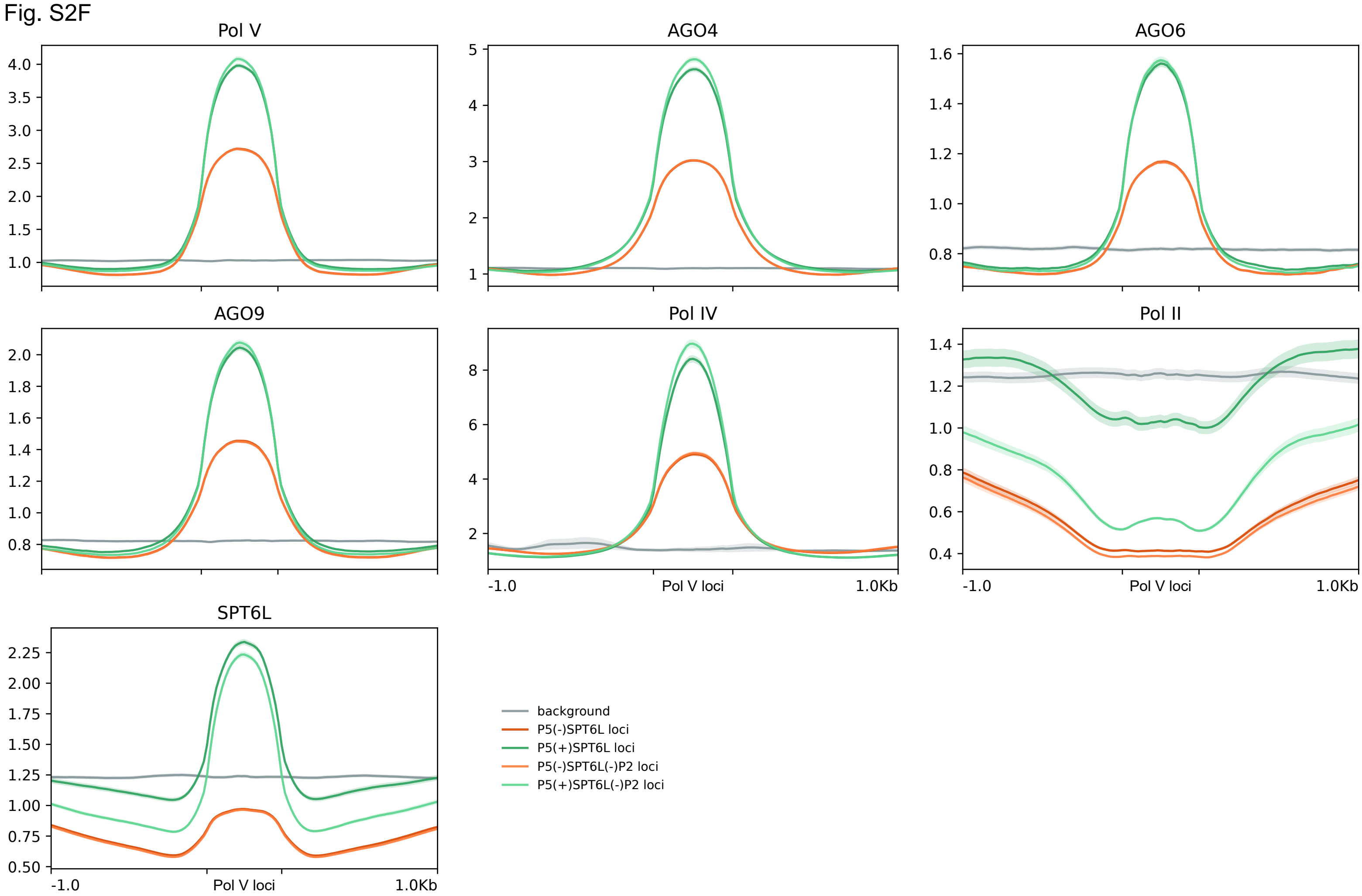
Pol V
AGO4
AGO6
AGO9
Pol IV
Pol II
SPT6L
-1.0
Pol V loci
1.0Kb
background
P5(-)SPT6L loci
P5(+)SPT6L loci
P5(-)SPT6L(-)P2 loci
P5(+)SPT6L(-)P2 loci

#### Supplemental Fig. S3: Mode of DNA methylation, types of sRNAs and genomic elements at Pol V loci occupied by SPT6L

Extended version of Fig. 3 to show how the analyzed loci are affected by Pol II. (A,B) Enrichment/depletion of differentially methylated regions (DMRs) at P5(+)SPT6L(-)P2 loci in datasets/samples from Zhang et al. 2018 containing whole genome methylation data for various mutants and some tissues. Violin plots show how the individual samples are enriched at P5(+)SPT6L(-)P2 loci (i.e. whether DMRs from the sample tend to overlap the P5(+)SPT6L(-)P2 loci more than the P5(-)SPT6L(-)P2 loci): (A) sample assessment based on DMR type (cytosine context and hyper-/hypomethylation) and by (B) the sample category that is shown only for hypo-CHH DMRs in the sample. Only datasets significantly enriched/depleted for the P5(+)SPT6(-)P2 loci are shown. In panel A the width of the violin plot is scaled to the number of samples and vertical lines indicate median and quartiles, in panel B the plots are not scaled and vertical lines indicate individual datasets; position of *nrpe1* mutant datasets is outlined below by the red rectangle. Data for individual datasets and their classification can be found in Supplemental Table S3. (C) Heatmap with enrichment of sRNA at P5(+)SPT6L(-)P2 loci for indicated samples and sRNA lengths calculated as log2(P5(+)SPT6L(-)P2/P5(-)SPT6L(-)P2). Analyzed sRNA-seq datasets originate either from AGO immunoprecipitation or from total sRNA samples in case of "input" and WT, and *nrpe1* and *nrpd1* mutants. Samples outlined within the same rectangle come from the same experiment (these are well comparable with each other). Individual replicates are shown in Supplemental Table S4. (D) Enrichment/depletion of overlap of P5(+)SPT6L and P5(+)SPT6L(-)P2 loci at different types of genomic elements (see Methods for genomic elements definition) and TE categories as defined by Panda et al. 2016: TE family, TE expression, Type of TE silencing in WT (either canonical PolIV-RdDM or various non-canonical RdDM pathways, non-RdDM pathways or not determined, ND), Type of TE silencing in *ddm1*. The "n" indicates the total number of genomic elements or TEs in a given category. (E) Characteristic features of TEs overlapping P5(+)SPT6L, P5(-)SPT6L, P5(+)SPT6L(-)P2 and P5(-)SPT6L(-)P2 loci, features adapted from Panda et al. 2016: TE copy number, TE length, TE gene distance and TE centromere distance. Significance is indicated by asterisks: $p < 0.05$ = *; $p < 0.005$ = **; $p < 0.0005$ = ***; NS = not significant. The significance indicates either enrichment of P5(+)SPT6L or P5(+)SPT6L(-)P2 in panel D, or significant difference between P5(+)SPT6L and P5(-)SPT6L or P5(+)SPT6L(-)P2 and P5(-)SPT6L(-)P2 in panel E.

Fig. S3A

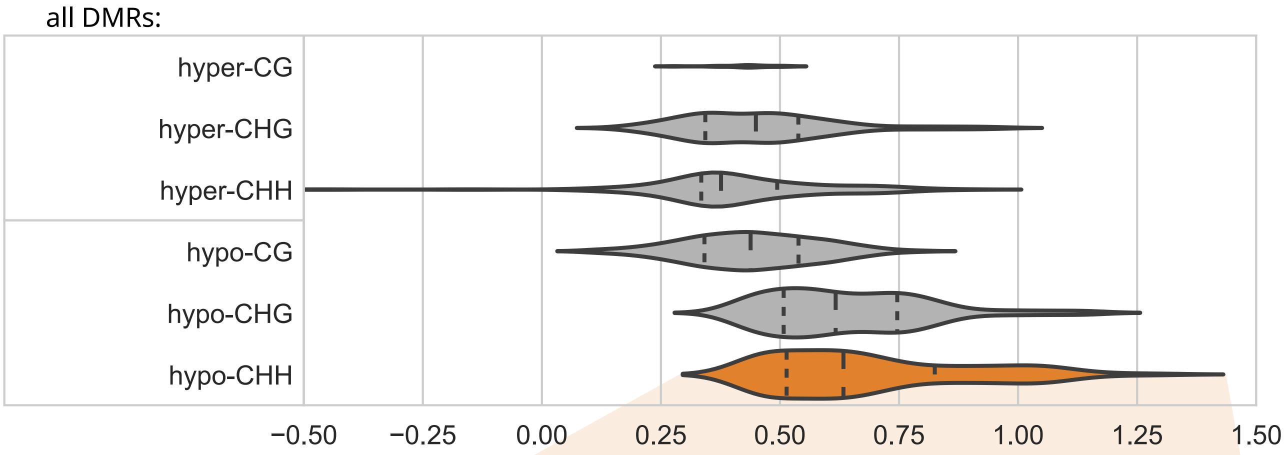


Fig. S3B

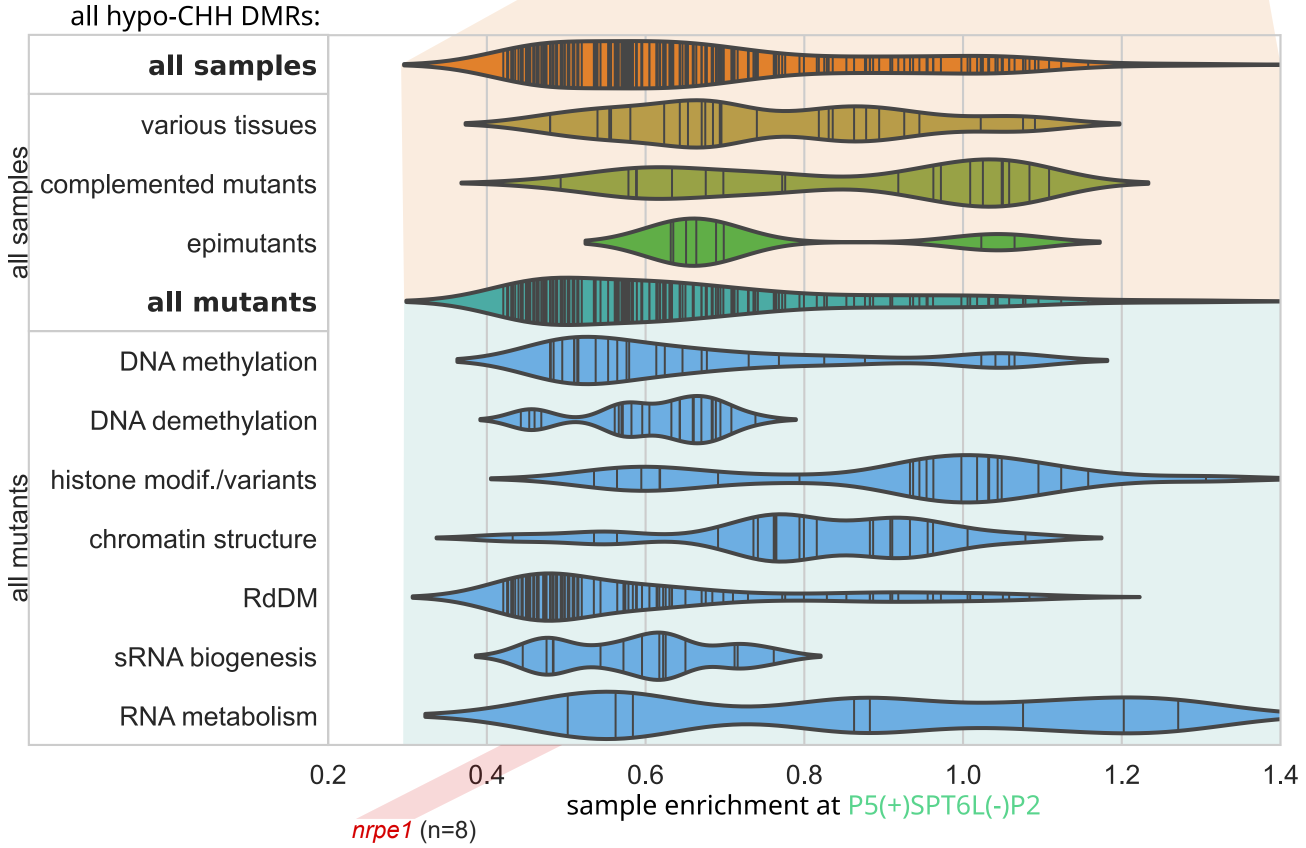


Fig. S3C

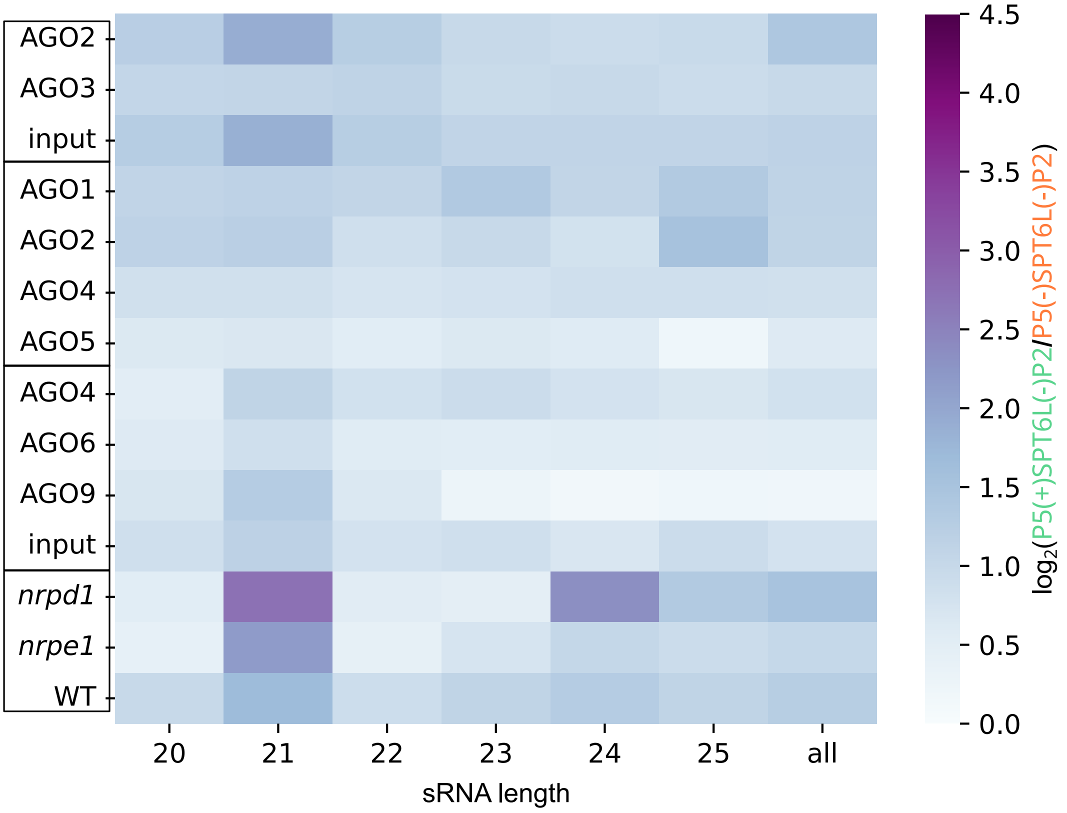

Fig. S3D

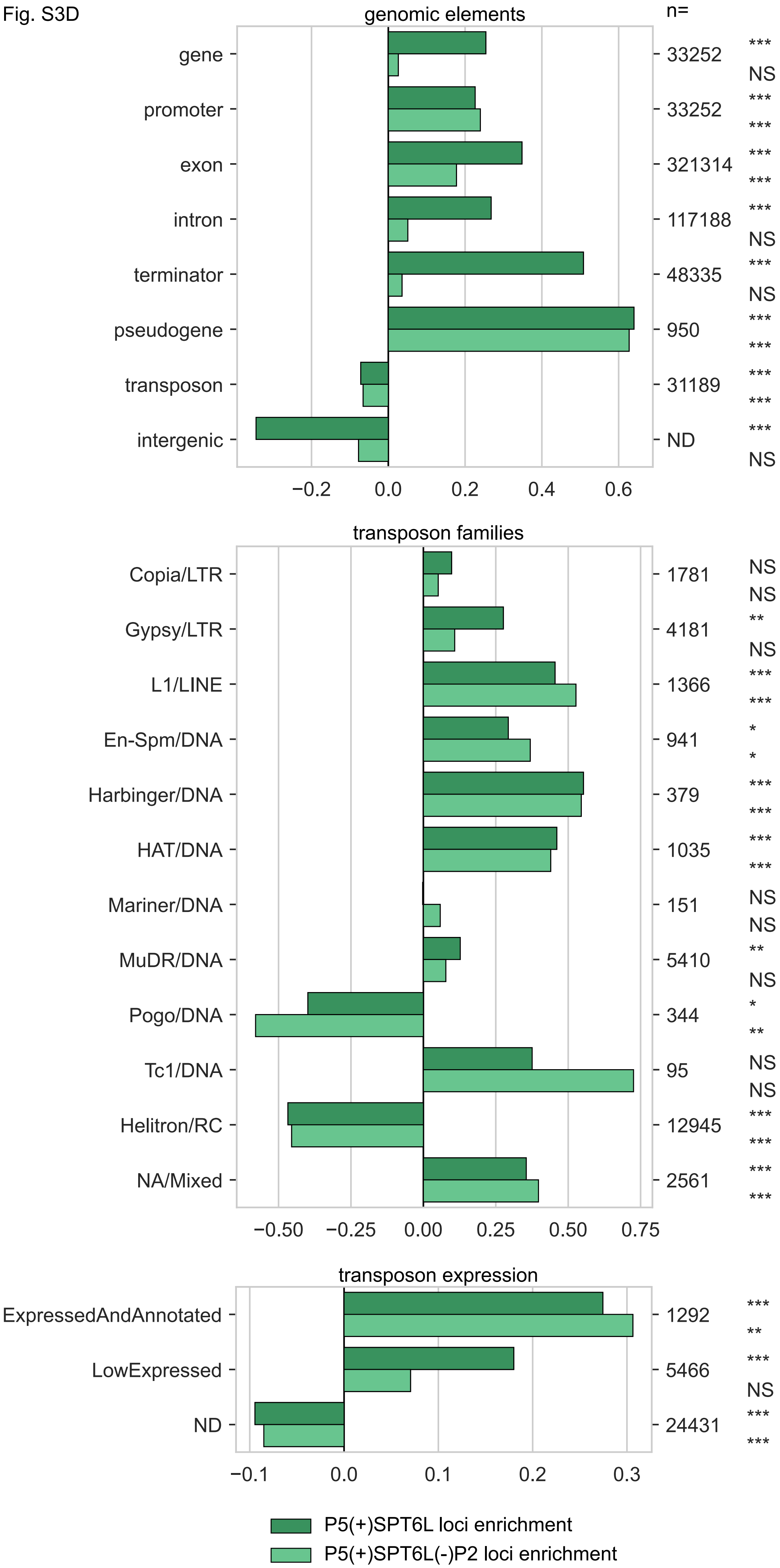

genomic elements
n=
gene
promoter
exon
intron
terminator
pseudogene
transposon
intergenic
33252
33252
321314
117188
48335
950
31189
ND
***
NS
***
***
***
***
***
NS
***
NS
***
***
***
***
***
NS
−0.2
0.0
0.2
0.4
0.6
transposon families
Copia/LTR
Gypsy/LTR
L1/LINE
En-Spm/DNA
Harbinger/DNA
HAT/DNA
Mariner/DNA
MuDR/DNA
Pogo/DNA
Tc1/DNA
Helitron/RC
NA/Mixed
1781
4181
1366
941
379
1035
151
5410
344
95
12945
2561
NS
NS
**
NS
***
***
*
*
***
***
***
***
NS
NS
**
NS
*
**
NS
NS
***
***
***
***
−0.50
−0.25
0.00
0.25
0.50
0.75
transposon expression
ExpressedAndAnnotated
LowExpressed
ND
1292
5466
24431
***
**
***
NS
***
***
−0.1
0.0
0.1
0.2
0.3
P5(+)SPT6L loci enrichment
P5(+)SPT6L(-)P2 loci enrichment

Fig. S3D

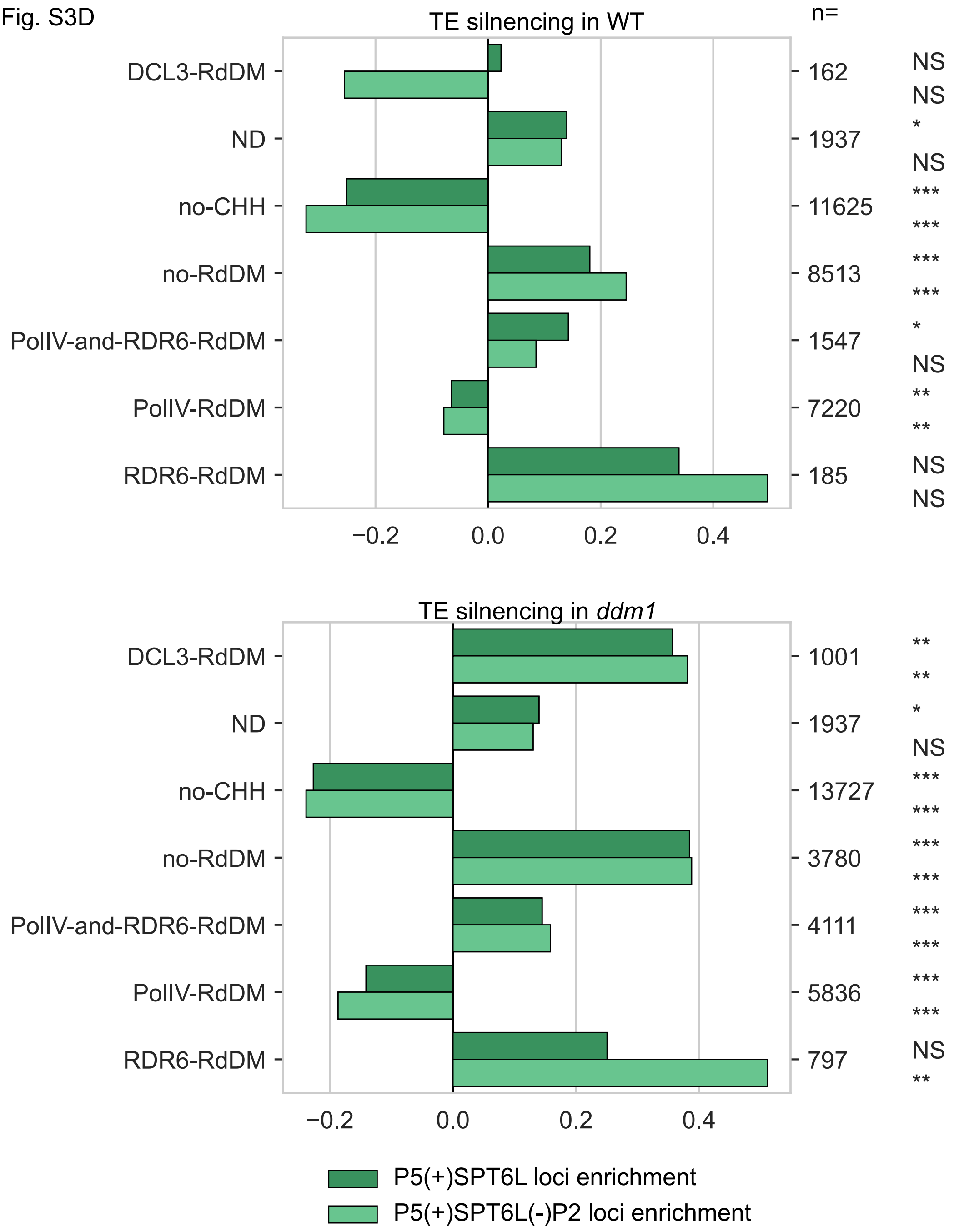

TE silnencing in WT
n=
DCL3-RdDM
ND
no-CHH
no-RdDM
PolIV-and-RDR6-RdDM
PolIV-RdDM
RDR6-RdDM
162
1937
11625
8513
1547
7220
185
NS
NS
*
NS
***
***
***
***
*
NS
**
**
NS
NS
−0.2
0.0
0.2
0.4
TE silnencing in ddm1
DCL3-RdDM
ND
no-CHH
no-RdDM
PolIV-and-RDR6-RdDM
PolIV-RdDM
RDR6-RdDM
1001
1937
13727
3780
4111
5836
797
**
**
*
NS
***
***
***
***
***
***
***
***
NS
**
−0.2
0.0
0.2
0.4
P5(+)SPT6L loci enrichment
P5(+)SPT6L(-)P2 loci enrichment

Fig. S3E

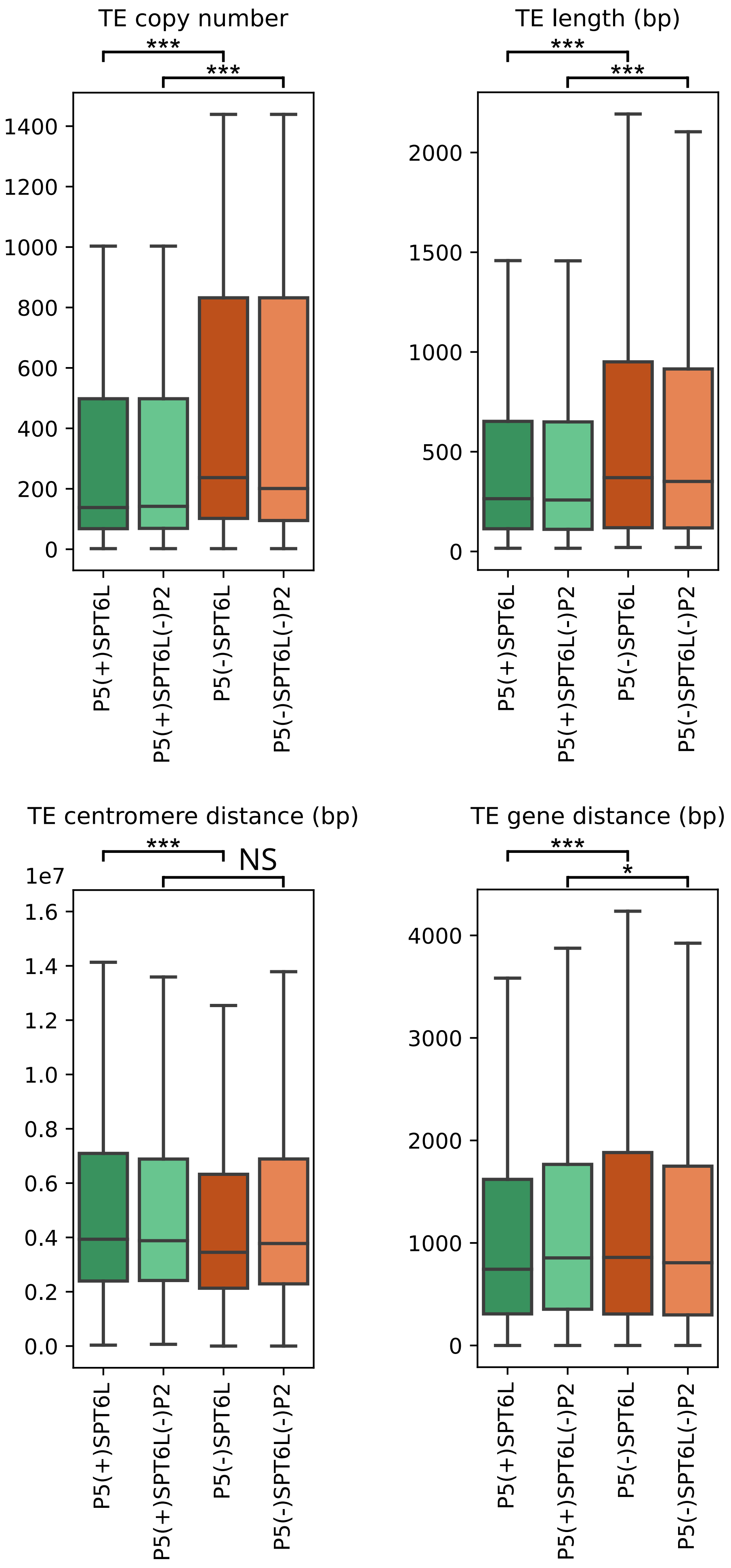

TE copy number
***
***
1400
1200
1000
800
600
400
200
0
P5(+)SPT6L
P5(+)SPT6L(-)P2
P5(-)SPT6L
P5(-)SPT6L(-)P2
TE length (bp)
***
***
2000
1500
1000
500
0
TE centromere distance (bp)
***
NS
1e7
1.6
1.4
1.2
1.0
0.8
0.6
0.4
0.2
0.0
TE gene distance (bp)
***
*
4000
3000
2000
1000
0

#### Supplemental Fig S4: Evolution of SPT6L, other relevant AGO-hook proteins and DNA methyltransferases in Viridiplantae

(A-G) Rooted maximum-likelihood trees depict the evolution of (A) SPT6/SPT6L, (B) SPT5/SPT5L, (C) NERD, (D) SDE3 and its homologues, (E) largest subunit of RNA polymerases II, IV and V, (F) DNMT3 and DRMs, (G) DNMT1 and CMTs proteins in the *Viridiplantae* clade. Bootstrap values are shown in the branching points of the tree. (A-D) Red coloring depicts proteins that carry an AGO-hook domain (containing at least 3 WG/GW motives), whereas blue coloring indicates those not possessing this domain. The number of GW/WG motifs indicates the length of this domain. (C) Tree for NERD is unrooted. (E) Yellow coloring depicts the largest subunit of Pol II, blue of Pol IV, light green of Pol V and darker green of Pol IV/V (ancestor of Pols IV and V). The scale bar in a RAxML trees is an estimation of the number of nucleotide substitutions per site. (H-I) Analysis of the C-terminal domain of SPT6(L) in the *Klebsormidium* genus. *Klebsormidium* seems to be the only streptophyte algae that doesn't possess SPT6L with an AGO-hook domain. The C-terminal domain of SPT6(L) in neither *K. flaccidum* nor *K. nitens* did comply with our stringent filter, even though these domains resemble AGO-hook domains in other SPT6L proteins. It remains unclear whether even the truncated AGO-hook-like domain of SPT6(L) in the *Klebsormidium* genus is functional, or if this domain was lost. (H) Sashimi plot of contig kfl00507 of *Klebsormidium nitens* NIES-2285 genome with aligned RNA-seq reads from sequencing project GSM5397554. Reads were aligned with RNA-STAR with default options allowing long introns. There seems to be no introns spanning further than the annotated sequence for SPT6L which rules out the possibility that the domain was originally not found due to alternative splicing. (I) Scheme of CTDs of obtained sequences for SPT6(L) protein from Arabidopsis thaliana and *Klebsormidium nitens*. Colors from blue (low) to red (high) denote the probability of AGO-hook motifs. Remark: when the eukaryotic matrix for W-search is used instead of the plant matrix, then the SPT6(L) passes our criteria as the third tryptophan residue passes our threshold.

Fig. S4A

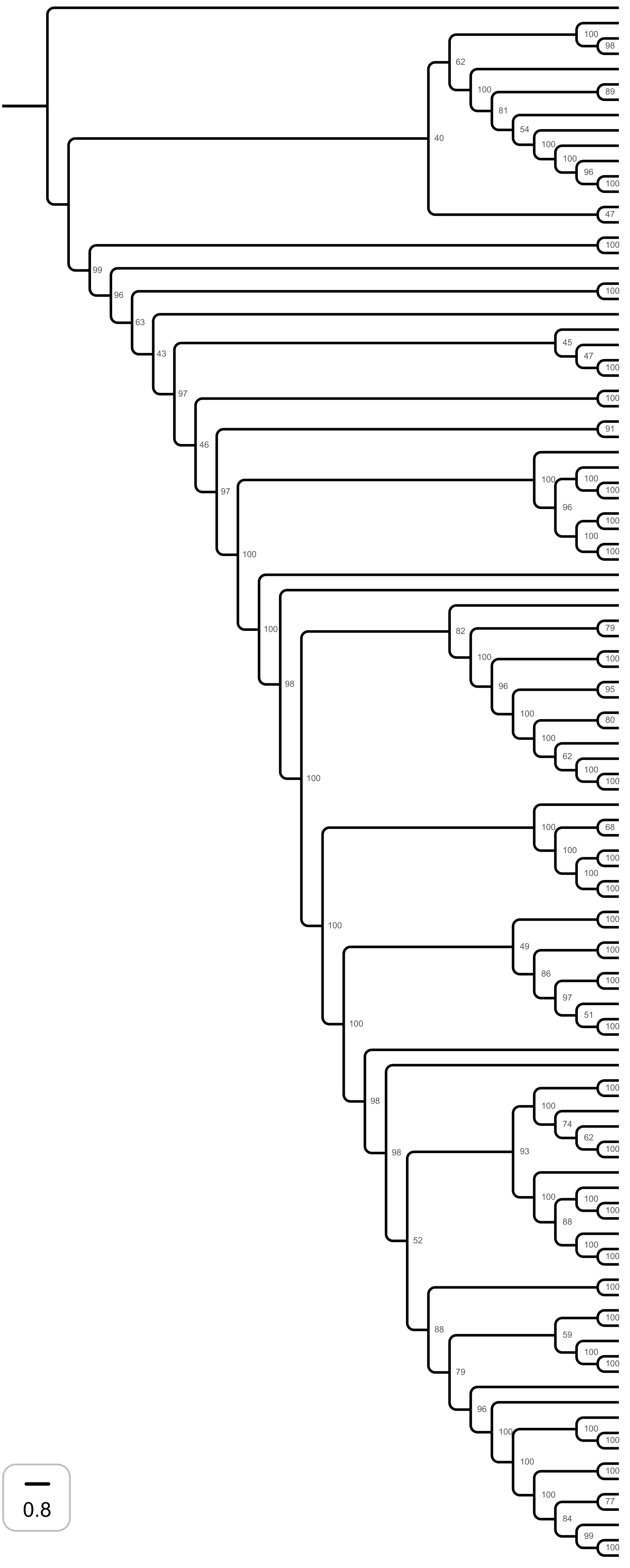


<table>
<tr><th>Species</th><th>Sequence ID</th><th># GW/WG</th></tr>
<tr><td>Galdieria sulphuraria</td><td>XM_005707307.1</td><td>0</td></tr>
<tr><td>Micromonas pusilla</td><td>XP_003057540.1</td><td>1</td></tr>
<tr><td>Ostreococcus lucimarinus</td><td>XP_001420894.1</td><td>1</td></tr>
<tr><td>Bathycoccus prasinos</td><td>XP_007514822.1</td><td>0</td></tr>
<tr><td>Ostreobium quekettii</td><td>CAD7702599.1</td><td>2</td></tr>
<tr><td>Coccomyxa subellipsoidea</td><td>C_169_XP_005647292.1</td><td>1</td></tr>
<tr><td>Trebouxia sp.</td><td>A1_2_KAA6428789.1</td><td>0</td></tr>
<tr><td>Auxenochlorella protothecoides</td><td>XP_011397843.1</td><td>0</td></tr>
<tr><td>Scenedesmus sp.</td><td>KAF6260805.1</td><td>0</td></tr>
<tr><td>Chlamydomonas reinhardtii</td><td>XP_042921444.1</td><td>0</td></tr>
<tr><td>Edaphochlamys debaryana</td><td>KAG2488784.1</td><td>1</td></tr>
<tr><td>Volvox carteri</td><td>XP_002958667.1</td><td>1</td></tr>
<tr><td>Astrephomene gubernaculifera</td><td>GFR40315.1</td><td>0</td></tr>
<tr><td>Chlorokybus atmophyticus</td><td>GFZF01006451.1</td><td>7</td></tr>
<tr><td>Mesostigma viride</td><td>jgi|Mesvir1|20970|Mv08037_RA.1</td><td>5</td></tr>
<tr><td>Interfilum paradoxum</td><td>FPCO_scaffold_2030348</td><td>3</td></tr>
<tr><td>Klebsormidium nitens</td><td>kfl00507_0030_v1.1</td><td>2</td></tr>
<tr><td>Coleochaete scutata</td><td>GFXZ01004166.1</td><td>4</td></tr>
<tr><td>Mesotaenium kramstae</td><td>jgi|Meskra657_3|2180028</td><td>3</td></tr>
<tr><td>Mougeotia calospora</td><td>GJZN01001526.1</td><td>5</td></tr>
<tr><td>Chara braunii</td><td>GBG67752.1</td><td>26</td></tr>
<tr><td>Anthoceros angustus</td><td>MU008166.1</td><td>6</td></tr>
<tr><td>Marchantia polymorpha</td><td>PTQ43424</td><td>34</td></tr>
<tr><td>Physcomitrium patens</td><td>XM_024520549.1</td><td>9</td></tr>
<tr><td>Physcomitrium patens</td><td>XM_024519963.1</td><td>11</td></tr>
<tr><td>Selaginella moellendorfii</td><td>XP_024519823</td><td>10</td></tr>
<tr><td>Selaginella moellendorffii</td><td>XM_024664055.1</td><td>7</td></tr>
<tr><td>Ceratopteris richardii</td><td>KAH7290621</td><td>21</td></tr>
<tr><td>Ceratopteris richardii</td><td>KAH7314514</td><td>14</td></tr>
<tr><td>Cycas revoluta</td><td>GBJU01001637.1</td><td>16</td></tr>
<tr><td>Cryptomeria japonica</td><td>ICQT01009851.1</td><td>12</td></tr>
<tr><td>Thuja plicata</td><td>Thupl.29382020s0009.1.p</td><td>11</td></tr>
<tr><td>Thuja occidentalis</td><td>GGJL01009513.1</td><td>13</td></tr>
<tr><td>Pinus sylvestris</td><td>GILO01692244.1</td><td>13</td></tr>
<tr><td>Pinus sylvestris</td><td>GILP01759947.1</td><td>13</td></tr>
<tr><td>Abies pinsapo</td><td>GCZN01015930.1</td><td>13</td></tr>
<tr><td>Abies pinsapo</td><td>GCZN01015928.1</td><td>13</td></tr>
<tr><td>Amborella trichopoda</td><td>XM_020672699.1</td><td>21</td></tr>
<tr><td>Nymphaea colorata</td><td>v1.2|Nycol.A02963.1.p</td><td>7</td></tr>
<tr><td>Liriodendron tulipifera</td><td>GHTS01012009.1</td><td>12</td></tr>
<tr><td>Dendrobium catenatum</td><td>XM_020816634.2</td><td>13</td></tr>
<tr><td>Asparagus officinalis</td><td>XM_020385771.1</td><td>6</td></tr>
<tr><td>Musa acuminata</td><td>v1|GSMUA_Achr2P00830_001</td><td>10</td></tr>
<tr><td>Musa acuminata</td><td>v1|GSMUA_Achr11P15640_001</td><td>2</td></tr>
<tr><td>Ananas comosus</td><td>XM_020244307.1</td><td>46</td></tr>
<tr><td>Ananas comosus</td><td>XM_020234201.1</td><td>4</td></tr>
<tr><td>Brachypodium distachyon</td><td>XM_014897374.2</td><td>1</td></tr>
<tr><td>Brachypodium distachyon</td><td>XM_010232918.3</td><td>11</td></tr>
<tr><td>Oryza sativa</td><td>XM_015783221.2</td><td>12</td></tr>
<tr><td>Zea mays</td><td>XM_008651595.2</td><td>9</td></tr>
<tr><td>Setaria viridis</td><td>XM_034732657.1</td><td>15</td></tr>
<tr><td>Panicum hallii</td><td>XM_025950289.1</td><td>12</td></tr>
<tr><td>Vitis vinifera</td><td>XM_010647114.2</td><td>17</td></tr>
<tr><td>Theobroma cacao</td><td>XM_007010677.2</td><td>7</td></tr>
<tr><td>Populus trichocarpa</td><td>XM_024593835.1</td><td>7</td></tr>
<tr><td>Glycine max</td><td>Wm82.a2.v1|Glyma.06G312200.1.p</td><td>7</td></tr>
<tr><td>Vigna unguiculata</td><td>XM_028063641.1</td><td>18</td></tr>
<tr><td>Medicago truncatula</td><td>XM_024779411.2</td><td>24</td></tr>
<tr><td>Cicer arietinum</td><td>XM_004499712.3</td><td>24</td></tr>
<tr><td>Spinacia oleracea</td><td>Spov3|Spov3_chr4.02535</td><td>9</td></tr>
<tr><td>Beta vulgaris</td><td>XM_010674818.2</td><td>10</td></tr>
<tr><td>Lactuca sativa</td><td>V8|Lsat_1_v5_gn_8_83860.1</td><td>3</td></tr>
<tr><td>Lactuca sativa</td><td>V8|Lsat_1_v5_gn_9_32200.1</td><td>24</td></tr>
<tr><td>Nicotiana tabacum</td><td>XM_016614012.1</td><td>8</td></tr>
<tr><td>Solanum tuberosum</td><td>XM_006339186.2</td><td>7</td></tr>
<tr><td>Olea europaea</td><td>XM_022990656.1</td><td>10</td></tr>
<tr><td>Coffea arabica</td><td>XM_027215630.1</td><td>0</td></tr>
<tr><td>Coffea arabica</td><td>XM_027215633.1</td><td>12</td></tr>
<tr><td>Vitis vinifera</td><td>XM_010655357.2</td><td>8</td></tr>
<tr><td>Eucalyptus grandis</td><td>XM_039304186.1</td><td>8</td></tr>
<tr><td>Prunus persica</td><td>XM_020569586.1</td><td>2</td></tr>
<tr><td>Malus domestica</td><td>v1.1|MD10G1173400</td><td>5</td></tr>
<tr><td>Fragaria vesca</td><td>XM_011472667.1</td><td>9</td></tr>
<tr><td>Prunus persica</td><td>XM_020567728.1</td><td>8</td></tr>
<tr><td>Malus domestica</td><td>v1.1|MD00G1108000</td><td>10</td></tr>
<tr><td>Malus domestica</td><td>v1.1|MD00G1025500</td><td>10</td></tr>
<tr><td>Arachis duranensis</td><td>XM_016115189.2</td><td>10</td></tr>
<tr><td>Vigna unguiculata</td><td>XM_028069878.1</td><td>8</td></tr>
<tr><td>Glycine max</td><td>Wm82.a2.v1|Glyma.19G125600.1.p</td><td>8</td></tr>
<tr><td>Glycine max</td><td>Wm82.a2.v1|Glyma.03G121000.1.p</td><td>9</td></tr>
<tr><td>Cicer arietinum</td><td>XM_004493257.3</td><td>7</td></tr>
<tr><td>Medicago truncatula</td><td>XM_039827554.1</td><td>11</td></tr>
<tr><td>Medicago truncatula</td><td>XM_039827995.1</td><td>1</td></tr>
<tr><td>Citrus clementina</td><td>XM_024182253.1</td><td>7</td></tr>
<tr><td>Citrus clementina</td><td>v1.0|Ciclev10013566m</td><td>7</td></tr>
<tr><td>Populus trichocarpa</td><td>XM_002322561.3</td><td>11</td></tr>
<tr><td>Populus trichocarpa</td><td>XM_024604608.1</td><td>7</td></tr>
<tr><td>Theobroma cacao</td><td>XM_007010649.2</td><td>7</td></tr>
<tr><td>Gossypium raimondii</td><td>XM_012599158.1</td><td>2</td></tr>
<tr><td>Gossypium raimondii</td><td>XM_012593495.1</td><td>9</td></tr>
<tr><td>Carica papaya</td><td>XM_022035839.1</td><td>2</td></tr>
<tr><td>Tarenaya hassleriana</td><td>XP_010528633.1</td><td>9</td></tr>
<tr><td>Capsella rubella</td><td>XM_023788783.1</td><td>0</td></tr>
<tr><td>Arabidopsis thaliana</td><td>NM_105000.1</td><td>0</td></tr>
<tr><td>Capsella rubella</td><td>XM_023788580.1</td><td>0</td></tr>
<tr><td>Capsella rubella</td><td>XM_006300582.2</td><td>13</td></tr>
<tr><td>Arabidopsis thaliana</td><td>NM_105218.5</td><td>12</td></tr>
<tr><td>Eutrema salsugineum</td><td>XM_024156252.1</td><td>11</td></tr>
<tr><td>Eutrema salsugineum</td><td>XM_024150584.1</td><td>2</td></tr>
<tr><td>Brassica rapa</td><td>v1.3|Brara.B01624.1.p</td><td>8</td></tr>
<tr><td>Brassica oleracea</td><td>XM_013736047.1</td><td>9</td></tr>
<tr><td>Brassica rapa</td><td>v1.3|Brara.G02597.1.p</td><td>8</td></tr>
</table>

### Fig. S4B

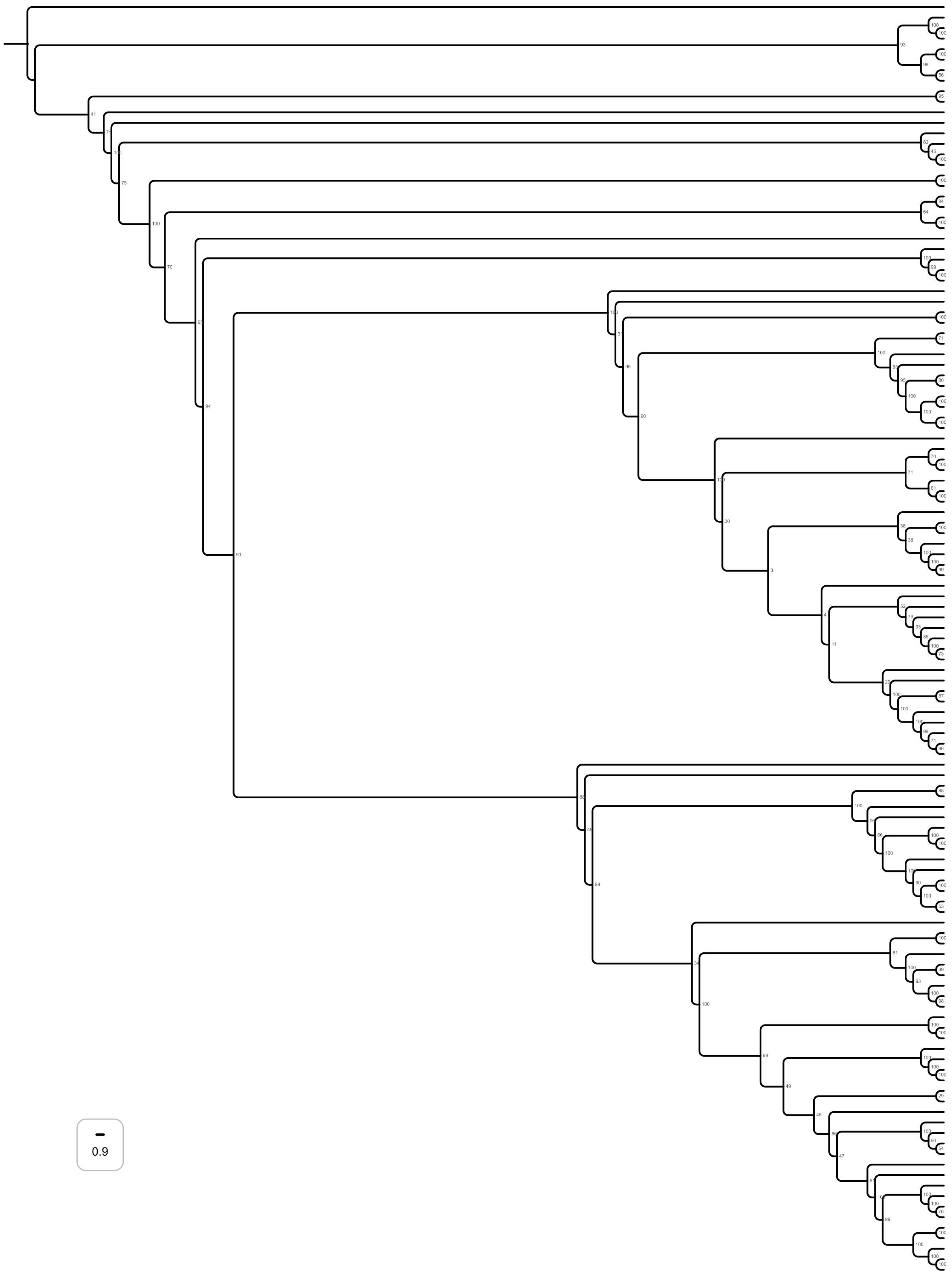


| Species | Sequence ID | # GW/WG |
|---|---|---|
| *Galdieria sulphuraria* | GsulpMS1_G3830.1 | **0** |
| *Ostreococcus lucimarinus* | XM_001418238.1 | **1** |
| *Micromonas commoda* | XM_002499716.1 | **0** |
| *Micromonas pusilla* | XM_003062105.1 | **1** |
| *Volvox carteri* | v2.1\|Vocar.0001s0920.1.p | **1** |
| *Chlamydomonas reinhardtii* | XM_001690005.2 | **0** |
| *Coccomyxa subellipsoidea* | XM_005645479.1 | **0** |
| *Chlorella variabilis* | XM_005842754.1 | **0** |
| *Chlorokybus atmophyticus* | Chrsp234S03564 | **2** |
| *Mesostigma viride* | GBSK01044010.1 | **0** |
| *Klebsormidium flaccidum* | GBSO01024704.1 | **1** |
| *Chara braunii* | GGXX01117120.1 | **2** |
| *Coleochaete orbicularis* | GBSL01013226.1 | **1** |
| *Spiroglea muscicola* | jgi\|Spimu1\|12625\|SM000026S08975 | **1** |
| *Mesotaenium kramstae* | jgi\|Meskra657_3\|2701821 | **2** |
| *Mougeotia sp.* | MZCH_SVCK_240 | **1** |
| *Selaginella bryopteris* | GEMU01096050.1 | **1** |
| *Selaginella denticulata* | GIMH01012008.1 | **1** |
| *Anthoceros angustus* | VJWM01000025.1 | **1** |
| *Marchantia polymorpha* | PTQ42993.1 | **1** |
| *Physcomitrium patens* | XM_024543534 | **1** |
| *Physcomitrium patens* | XM_024509952.1 | **1** |
| *Ceratopteris richardii* | KAH7297507.1 | **2** |
| *Cycas revoluta* | GBJU01014550.1 | **1** |
| *Thuja plicata* | v3.1c\|Thupl.29380412s0002.1.p | **2** |
| *Pinus tecunumanii* | GFZL01027662.1 | **2** |
| *Abies pinsapo* | GCZN01028779.1 | **2** |
| *Amborella trichopoda* | XM_020664155 | **60** |
| *Nymphaea colorata* | XM_031631806 | **43** |
| *Liriodendron tulipifera* | v1.1\|Litul.01G132600.1.p | **49** |
| *Liriodendron tulipifera* | v1.1\|Litul.15G046300.1.p | **35** |
| *Asparagus officinalis* | V1.1\|evm.model.AsparagusV1_03.1993 | **54** |
| *Phalaenopsis equestris* | XM_020731481 | **50** |
| *Musa acuminata* | XP_009409884.1 | **36** |
| *Ananas comosus* | v3\|Aco005272.1 | **50** |
| *Oryza sativa* | Japonica_Group_XM_015783842 | **40** |
| *Brachypodium distachyon* | v3.2\|Bradi2g20517.2.p | **44** |
| *Zea mays* | XM_008658209 | **49** |
| *Zea mays* | XM_020539610 | **51** |
| *Setaria viridis* | v2.1\|Sevir.3G191100.2.p | **57** |
| *Panicum hallii* | v3.2\|Pahal.3G222100.1.p | **54** |
| *Eucalyptus grandis* | XM_010033834 | **64** |
| *Fragaria vesca* | XM_011471308 | **31** |
| *Spinacia oleracea* | Spov3\|Spov3_C0003.00068 | **22** |
| *Beta vulgaris* | EL10_1.0\|EL10Ac7g16663.1 | **42** |
| *Prunus persica* | XM_020557016 | **60** |
| *Malus domestica* | v1.1\|MD01G1100800 | **42** |
| *Malus domestica* | v1.1\|MD07G1167800 | **38** |
| *Populus trichocarpa* | XM_024607844 | **73** |
| *Gossypium raimondii* | v2.1\|Gorai.011G234600.1 | **49** |
| *Theobroma cacao* | v2.1\|Thecc.10G086900.1.p | **58** |
| *Medicago truncatula* | Mt4.0v1\|Medtr1g111560.1 | **32** |
| *Vigna unguiculata* | v1.2\|Vigun07g272900.1.p | **58** |
| *Glycine max* | Wm82.a2.v1\|Glyma.20G112800.1.p | **52** |
| *Glyma.10G276900.1.p* | Wm82.a2.v1\|Glyma.10G276900.1.p | **46** |
| *Carica papaya* | XM_022042070 | **13** |
| *Vitis vinifera* | XM_019224314 | **56** |
| *Lactuca sativa* | V8\|Lsat_1_v5_gn_2_112001.1 | **23** |
| *Solanum tuberosum* | XM_006361632 | **62** |
| *Olea europaea* | XP_022854124.1 | **50** |
| *Coffea arabica* | XM_027249107 | **61** |
| *Coffea eugenioides* | XM_027303492 | **65** |
| *Coffea arabica* | XM_027255656 | **63** |
| *Citrus clementina* | v1.0\|Ciclev10030480m | **70** |
| *Tarenaya hassleriana* | XP_010543566.1 | **67** |
| *Capsella rubella* | XM_006286833 | **51** |
| *Arabidopsis thaliana* | NM_001342760 | **54** |
| *Brassica oleracea* | XM_013755907 | **51** |
| *Brassica napus* | XM_022713865 | **21** |
| *Brassica rapa* | XM_009124368 | **57** |
| *Brassica napus* | XM_013812761 | **54** |
| *Brassica napus* | XM_013812954 | **56** |
| *Nymphaea colorata* | XM_031632769.1 | **1** |
| *Amborella trichopoda* | XM_011623260 | **2** |
| *Asparagus officinalis* | V1.1\|evm.model.AsparagusV1_05.376 | **2** |
| *Phalaenopsis equestris* | XM_020725506 | **1** |
| *Musa acuminata* | XP_009391480.1 | **1** |
| *Ananas comosus* | v3\|Aco002639.1 | **1** |
| *Brachypodium distachyon* | v3.2\|Bradi3g57590.2.p | **2** |
| *Panicum hallii* | v3.2\|Pahal.1G415200.2.p | **2** |
| *Setaria viridis* | v2.1\|Sevir.1G342100.1.p | **2** |
| *Oryza sativa* | XM_015788355 | **0** |
| *Brachypodium distachyon* | v3.2\|Bradi1g46230.4.p | **1** |
| *Zea mays* | XM_008660913 | **0** |
| *Zea mays* | XM_020539445 | **0** |
| *Panicum hallii* | v3.2\|Pahal.4G286300.1.p | **0** |
| *Setaria viridis* | v2.1\|Sevir.4G083700.1.p | **0** |
| *Liriodendron tulipifera* | v1.1\|Litul.15G046700.1.p | **2** |
| *Beta vulgaris* | EL10_1.0\|EL10Ac1g01470.1 | **1** |
| *Spinacia oleracea* | Spov3\|Spov3_chr5.00481 | **1** |
| *Lactuca sativa* | V8\|Lsat_1_v5_gn_3_132801.1 | **2** |
| *Olea europaea* | XP_022851445.1 | **2** |
| *Solanum tuberosum* | XM_006356238 | **2** |
| *Coffea arabica* | XM_027238315 | **1** |
| *Coffea arabica* | XM_027235696 | **1** |
| *Coffea eugenioides* | XM_027292439 | **1** |
| *Fragaria vesca* | XM_004292500 | **2** |
| *Malus domestica* | v1.1\|MD10G1013600 | **1** |
| *Prunus persica* | XM_007201697 | **1** |
| *Medicago truncatula* | Mt4.0v1\|Medtr3g089910.1 | **1** |
| *Vigna unguiculata* | v1.2\|Vigun09g155900.1.p | **1** |
| *Glycine max* | Wm82.a2.v1\|Glyma.06G122500.1.p | **1** |
| *Glycine max* | Wm82.a2.v1\|Glyma.04G240900.1.p | **0** |
| *Citrus clementina* | v1.0\|Ciclev10000121m | **2** |
| *Eucalyptus grandis* | XM_010063865 | **2** |
| *Populus trichocarpa* | XM_002305428 | **2** |
| *Theobroma cacao* | v2.1\|Thecc.08G143700.1.p | **2** |
| *Gossypium raimondii* | v2.1\|Gorai.009G252200.1 | **2** |
| *Gossypium raimondii* | v2.1\|Gorai.009G284500.1 | **2** |
| *Gossypium raimondii* | v2.1\|Gorai.005G082600.1 | **2** |
| *Carica papaya* | XM_022032025 | **2** |
| *Tarenaya hassleriana* | XP_010530229.1 | **1** |
| *Capsella rubella* | XM_023785155 | **2** |
| *Brassica rapa* | XM_009134730 | **2** |
| *Brassica napus* | XM_013843537 | **2** |
| *Brassica oleracea* | XM_013767266 | **2** |
| *Capsella rubella* | XM_006289130 | **2** |
| *Arabidopsis thaliana* | NM_116904 | **2** |
| *Brassica oleracea* | XM_013751925 | **0** |
| *Brassica rapa* | XM_009135838 | **2** |
| *Brassica napus* | XM_013828515 | **2** |

Fig. S4C

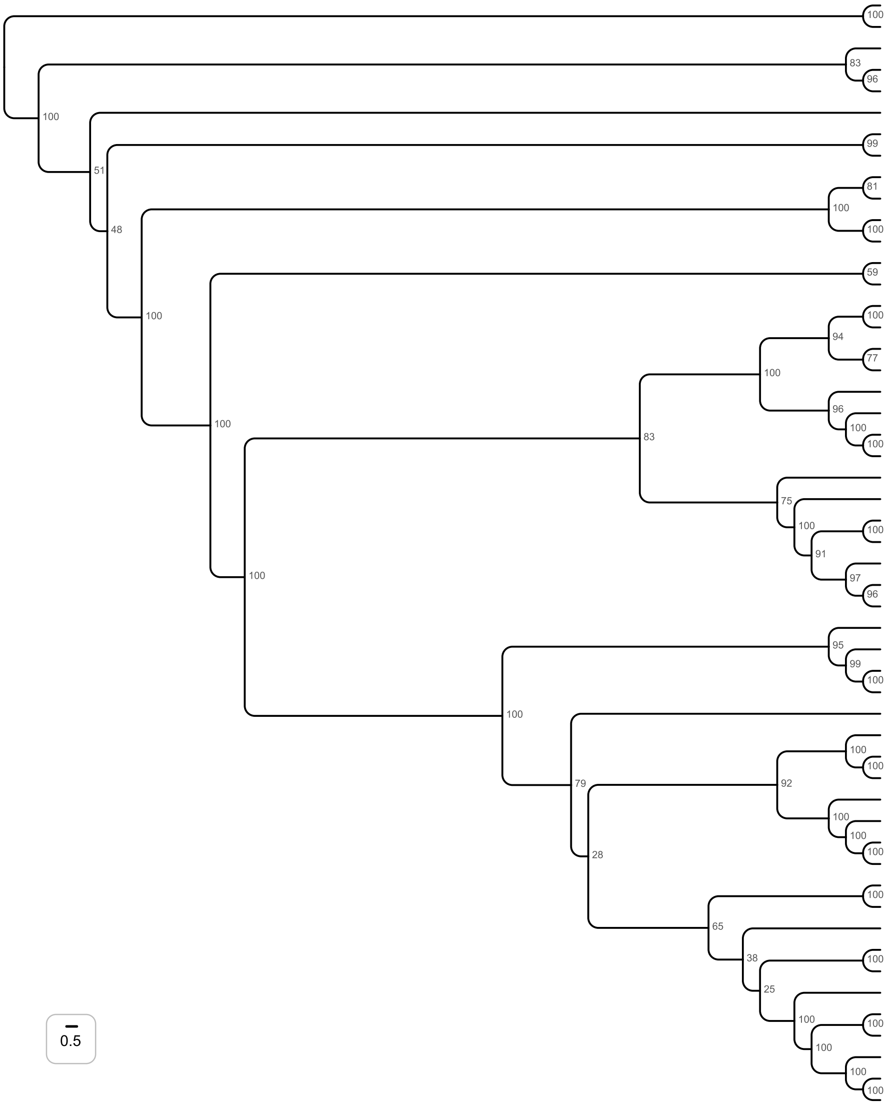


| Species | Sequence ID | # GW/WG |
|---|---|---|
| *Mesotaenium endlicherianum* | Mesen1\|5796\|ME000293S04950 | **5** |
| *Mesotaenium kramstae* | Meskra657_3\|204237\|CE203456_10532 | **4** |
| *Anthoceros angustus* | MU008185.1 | **4** |
| *Marchantia polymorpha* | GEFO01024060.1 | **5** |
| *Physcomitrium patens* | XM_024540864.1 | **6** |
| *Selaginella bryopteris* | GEMU01103855.1 | **4** |
| *Brainea insignis* | GFUE01030576.1 | **4** |
| *Ceratopteris richardii* | GIXU01028817.1 | **6** |
| *Cycas revoluta* | GGZS01014446.1 | **16** |
| *Ginkgo biloba* | GGBW01011546.1 | **24** |
| *Abies pinsapo* | GCZN01027697.1 | **17** |
| *Pinus sylvestris* | GHKX01012973.1 | **18** |
| *Nymphaea colorata* | XM_031627670.1 | **4** |
| *Amborella trichopoda* | XM_011626888.2 | **17** |
| *Ananas comosus* | XP_020093697.1 | **6** |
| *Oryza sativa* | XM_015758901.2 | **3** |
| *Asparagus officinalis* | V1.1\|AsparagusV1_Unassigned.582 | **11** |
| *Phalaenopsis equestris* | XM_020727459.1 | **7** |
| *Musa acuminata* | XP_009395393.1 | **3** |
| *Brachypodium distachyon* | v3.1\|Bradi4g35977.3.p | **4** |
| *Zea mays* | XM_008659006.4 | **6** |
| *Panicum hallii* | v3.2\|Pahal.5G162300.1.p | **6** |
| *Liriodendron tulipifera* | v1.1\|Litul.04G072000.1.p | **16** |
| *Vitis vinifera* | v2.1\|VIT_204s0023g02950.2 | **8** |
| *Coffea arabica* | XM_027221728.1 | **3** |
| *Coffea arabica* | XM_027217781.1 | **3** |
| *Malus domestica* | XM_008346037.3 | **3** |
| *Manihot esculenta* | XM_021749076.2 | **3** |
| *Carica papaya* | XM_022052612.1 | **0** |
| *Lactuca sativa* | V8\|Lsat_1_v5_gn_8_128721.1 | **6** |
| *Olea europaea* | XM_023040276.1 | **8** |
| *Solanum lycopersicum* | XM_010314914.3 | **3** |
| *Solanum tuberosum* | XM_006356320.2 | **3** |
| *Eucalyptus grandis* | XM_010067912.3 | **7** |
| *Fragaria vesca* | XM_004292388.2 | **11** |
| *Prunus persica* | v2.1\|Prupe.8G085700.1.p | **9** |
| *Prunus persica* | XP_020409470.1 | **9** |
| *Medicago truncatula* | XM_024779664.2 | **5** |
| *Vigna unguiculata* | v1.2\|Vigun09g269600.1.p | **5** |
| *Glycine max* | Wm82.a4.v1\|Glyma.06G008300.1.p | **5** |
| *Glycine soja* | XM_028378725.1 | **5** |
| *Populus alba* | XM_035071778.1 | **11** |
| *Manihot esculenta* | XM_021755439.2 | **7** |
| *Citrus clementina* | XM_006423406.2 | **5** |
| *Theobroma cacao* | v2.1\|Thecc.02G076000.1.p | **9** |
| *Gossypium raimondii* | XM_012624939.1 | **5** |
| *Tarenaya hassleriana* | XP_019056896.1 | **5** |
| *Capsella rubella* | XM_006296755.2 | **6** |
| *Arabidopsis thaliana* | NM_127202.5 | **6** |
| *Brassica oleracea* | v1.0\|Bol012694 | **6** |
| *Brassica rapa* | v1.3\|Brara.I00996.1.p | **6** |
| *Brassica napus* | XM_013800386.2 | **6** |

Fig. S4D

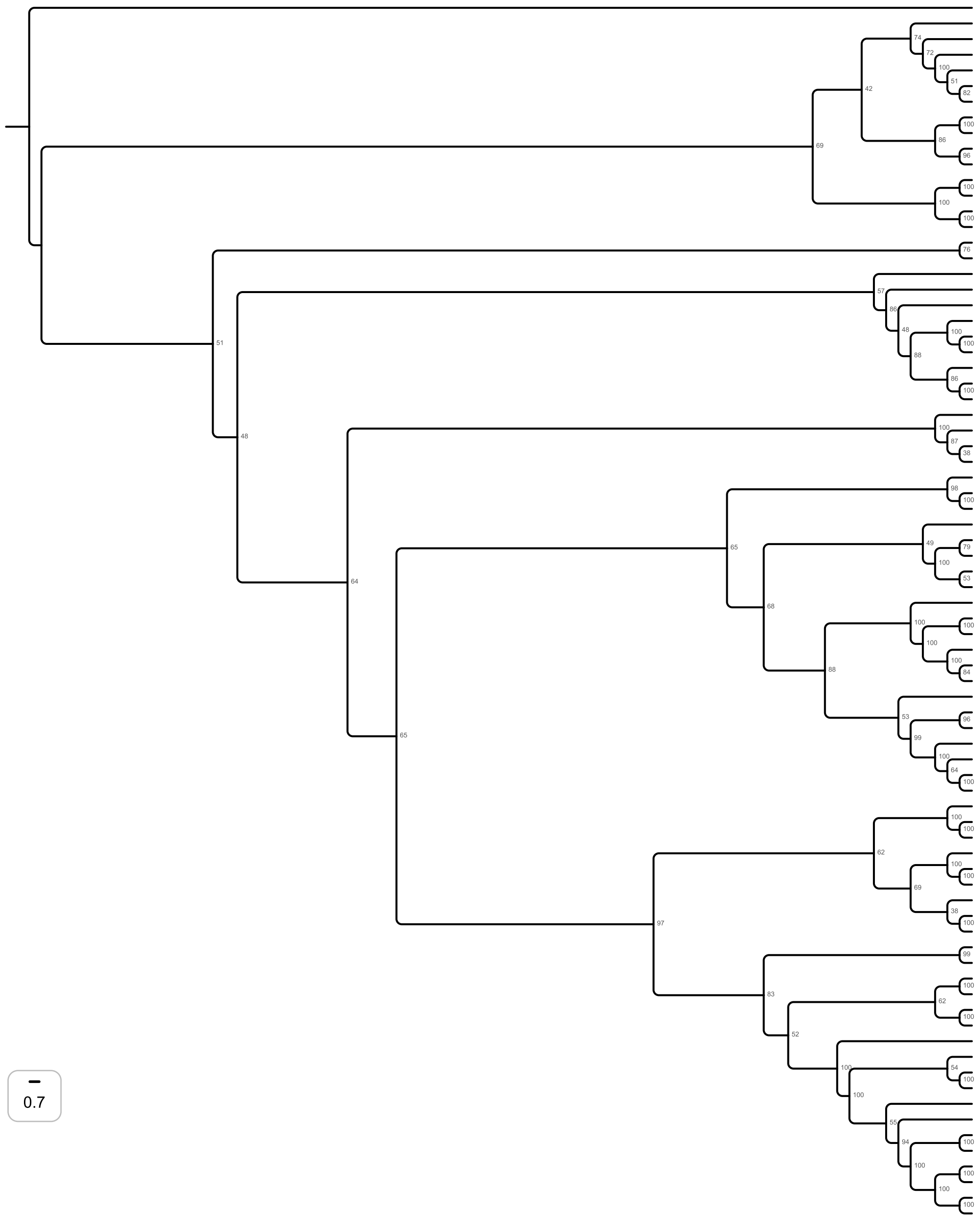


| Species | Sequence ID | # GW/WG |
|---|---|---|
| *Galdieria sulphuraria* | Gsulp5572_G6141.1_UPF1 | 1 |
| *Klebsormidium flaccidum* | kfl00006_0480_v1.1 | 0 |
| *Klebsormidium flaccidum* | kfl00077_0020_v1.1 | 0 |
| *Spiroglea muscicola* | jgi\|Spimu1\|23425 | 0 |
| *Mesotaenium kramstae* | jgi\|Meskra657_3\|2582996 | 0 |
| *Klebsormidium flaccidum* | kfl00120_0100_v1.1 | 0 |
| *Chara braunii* | CHBRA412g00140 | 1 |
| *Sellaginella moellendorffii* | v1.0\|236589 | 0 |
| *Sellaginella moellendorffii* | v1.0\|430895 | 0 |
| *Botrychium lunaria* | GIJZ01000402.1 | 0 |
| *Physcomitrella patens* | Pp3c12_8920V3.1 | 0 |
| *Thuja occidentalis* | GGJL01020185.1 | 0 |
| *Thuja plicata* | v3.1c\|Thupl.29380709s0004.1.p | 1 |
| *Pinus lambertiana* | GEUZ01002387.1 | 0 |
| *Pinus flexilis* | GHWE01188158.1 | 0 |
| *Amborella trichopoda* | XM_020664199.1 | 0 |
| *Nymphaea colorata* | XM_031619418.1 | 0 |
| *Liriodendron tulipifera* | v1.1\|Litul.16G005300.1.p | 0 |
| *Asparagus officinalis* | V1.1\|AsparagusV1_08.1887 | 0 |
| *Phalaenopsis equestris* | XM_020734797.1 | 0 |
| *Musa acuminata* | v1\|GSMUA_Achr3P25460_001 | 1 |
| *Zingiber officinale* | XM_042617426.1 | 1 |
| *Zingiber officinale* | XM_042612330.1 | 1 |
| *Ananas comosus* | v3\|Aco000226.1 | 1 |
| *Zea mays* | XM_008653011.3 | 0 |
| *Oryza sativa* | XM_015772278.2 | 0 |
| *Carica papaya* | XM_022035582.1 | 0 |
| *Populus trichocarpa* | XM_006382673.2 | 0 |
| *Eucalyptus grandis* | XM_039300975.1 | 0 |
| *Theobroma cacao* | v2.1\|Thecc.05G291200.1.p | 1 |
| *Camellia sinensis* | XM_028224285.1 | 0 |
| *Impatiens glandulifera* | XM_047477804.1 | 0 |
| *Impatiens glandulifera* | XM_047461405.1 | 0 |
| *Daucus carota* | XM_017400410.1 | 3 |
| *Cynara cardunculus* | XM_025112441.1 | 4 |
| *Lactuca sativa* | XM_023902859.2 | 7 |
| *Erigeron canadensis* | XM_043763207.1 | 5 |
| *Helianthus annuus* | XM_022154715.2 | 3 |
| *Olea europaea* | v1.0\|Oeu018974.1 | 1 |
| *Salvia hispanica* | XM_048096010.1 | 3 |
| *Salvia splendens* | XM_042200070.1 | 2 |
| *Salvia hispanica* | XM_048127576.1 | 2 |
| *Salvia splendens* | XM_042182159.1 | 3 |
| *Salvia splendens* | XM_042209340.1 | 3 |
| *Coffea arabica* | XM_027261613.1 | 0 |
| *Solanum lycopersicum* | XM_004240994.4 | 1 |
| *Capsicum annuum* | XM_016719860.2 | 6 |
| *Nicotiana attenuata* | XM_019410890.1 | 4 |
| *Nicotiana tabacum* | XM_016647526.1 | 2 |
| *Nicotiana tabacum* | XM_016608630.1 | 3 |
| *Nicotiana tomentosiformis* | XM_009619903.3 | 3 |
| *Glycine soja* | XM_028324683.1 | 1 |
| *Medicago truncatula* | XM_003595558.4 | 2 |
| *Medicago truncatula* | XM_003610728.3 | 2 |
| *Fragaria vesca* | XM_004297068.2 | 0 |
| *Prunus persica* | XM_020564494.1 | 1 |
| *Malus domestica* | v1.1\|MD13G1196500 | 0 |
| *Morus notabilis* | XM_010110024.2 | 1 |
| *Ziziphus jujuba* | XM_016032673.2 | 1 |
| *Ziziphus jujuba* | XM_025069116.1 | 1 |
| *Manihot esculenta* | XM_021751855.2 | 0 |
| *Populus trichocarpa* | XM_006379784.2 | 0 |
| *Gossypium raimondii* | XM_012631726.1 | 0 |
| *Gossypium hirsutum* | XM_041101358.1 | 0 |
| *Citrus clementina* | XM_024179974.1 | 2 |
| *Citrus sinensis* | XM_006489365.2 | 2 |
| *Tarenaya hassleriana* | XM_010523987.2 | 0 |
| *Capsella rubella* | XM_006306592.2 | 4 |
| *Camelina sativa* | XM_010486686.2 | 5 |
| *Camelina sativa* | XM_010482052.1 | 2 |
| *Arabidopsis thaliana* | NM_100425.4 | 5 |
| *Eutrema salsugineum* | XM_006417961.2 | 7 |
| *Raphanus sativus* | XM_018610486.1 | 0 |
| *Raphanus sativus* | XM_018580283.1 | 9 |
| *Brassica oleracea* | XM_013730784.1 | 5 |
| *Brassica napus* | XM_022702907.1 | 5 |
| *Brassica rapa* | XM_009120906.3 | 4 |
| *Brassica napus* | XM_022693149.1 | 4 |

Fig. S4E

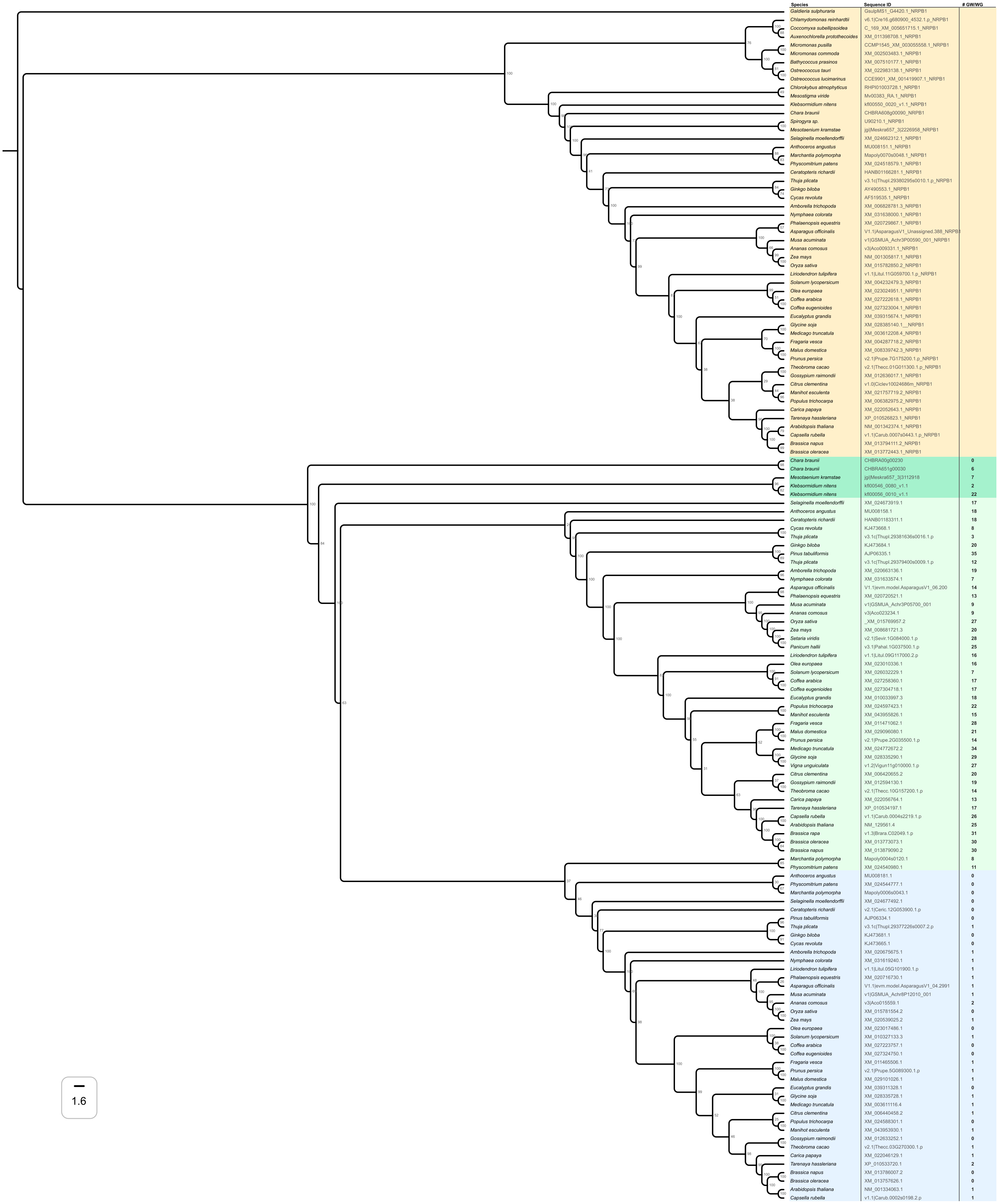
Species Sequence ID # GW/WG
Galdieria sulphuraria GsulpMS1_G4420.1_NRPB1
Chlamydomonas reinhardtii v6.1|Cre16.g680900_4532.1.p_NRPB1
Coccomyxa subellipsoidea C_169_XM_005651715.1_NRPB1
Auxenochlorella protothecoides XM_011398708.1_NRPB1
Micromonas pusilla CCMP1545_XM_003055558.1_NRPB1
Micromonas commoda XM_002503483.1_NRPB1
Bathycoccus prasinos XM_007510177.1_NRPB1
Ostreococcus tauri XM_022983138.1_NRPB1
Ostreococcus lucimarinus CCE9901_XM_001419907.1_NRPB1
Chlorokybus atmophyticus RHPI01003728.1_NRPB1
Mesostigma viride Mv00383_RA.1_NRPB1
Klebsormidium nitens kfl00550_0020_v1.1_NRPB1
Chara braunii CHBRA608g00090_NRPB1
Spirogyra sp. U90210.1_NRPB1
Mesotaenium kramstae jgi|Meskra657_3|2226958_NRPB1
Selaginella moellendorffii XM_024662312.1_NRPB1
Anthoceros angustus MU008151.1_NRPB1
Marchantia polymorpha Mapoly0070s0048.1_NRPB1
Physcomitrium patens XM_024518579.1_NRPB1
Ceratopteris richardii HANB01166281.1_NRPB1
Thuja plicata v3.1c|Thupl.29380295s0010.1.p_NRPB1
Ginkgo biloba AY490553.1_NRPB1
Cycas revoluta AF519535.1_NRPB1
Amborella trichopoda XM_006828781.3_NRPB1
Nymphaea colorata XM_031638000.1_NRPB1
Phalaenopsis equestris XM_020729867.1_NRPB1
Asparagus officinalis V1.1|AsparagusV1_Unassigned.388_NRPB1
Musa acuminata v1|GSMUA_Achr3P00590_001_NRPB1
Ananas comosus v3|Aco009331.1_NRPB1
Zea mays NM_001305817.1_NRPB1
Oryza sativa XM_015782850.2_NRPB1
Liriodendron tulipifera v1.1|Litul.11G059700.1.p_NRPB1
Solanum lycopersicum XM_004232479.3_NRPB1
Olea europaea XM_023024951.1_NRPB1
Coffea arabica XM_027222618.1_NRPB1
Coffea eugenioides XM_027323004.1_NRPB1
Eucalyptus grandis XM_039315674.1_NRPB1
Glycine soja XM_028385140.1__NRPB1
Medicago truncatula XM_003612208.4_NRPB1
Fragaria vesca XM_004287718.2_NRPB1
Malus domestica XM_008339742.3_NRPB1
Prunus persica v2.1|Prupe.7G175200.1.p_NRPB1
Theobroma cacao v2.1|Thecc.01G011300.1.p_NRPB1
Gossypium raimondii XM_012636017.1_NRPB1
Citrus clementina v1.0|Ciclev10024686m_NRPB1
Manihot esculenta XM_021757719.2_NRPB1
Populus trichocarpa XM_006382975.2_NRPB1
Carica papaya XM_022052643.1_NRPB1
Tarenaya hassleriana XP_010526823.1_NRPB1
Arabidopsis thaliana NM_001342374.1_NRPB1
Capsella rubella v1.1|Carub.0007s0443.1.p_NRPB1
Brassica napus XM_013794111.2_NRPB1
Brassica oleracea XM_013772443.1_NRPB1
Chara braunii CHBRA00g00230 0
Chara braunii CHBRA651g00030 6
Mesotaenium kramstae jgi|Meskra657_3|3112918 7
Klebsormidium nitens kfl00546_0080_v1.1 2
Klebsormidium nitens kfl00056_0010_v1.1 22
Selaginella moellendorffii XM_024673919.1 17
Anthoceros angustus MU008158.1 18
Ceratopteris richardii HANB01183311.1 18
Cycas revoluta KJ473668.1 8
Thuja plicata v3.1c|Thupl.29381636s0016.1.p 3
Ginkgo biloba KJ473684.1 20
Pinus tabuliformis AJP06335.1 35
Thuja plicata v3.1c|Thupl.29379400s0009.1.p 12
Amborella trichopoda XM_020663136.1 19
Nymphaea colorata XM_031633574.1 7
Asparagus officinalis V1.1|evm.model.AsparagusV1_06.200 14
Phalaenopsis equestris XM_020720521.1 13
Musa acuminata v1|GSMUA_Achr3P05700_001 9
Ananas comosus v3|Aco023234.1 9
Oryza sativa _XM_015769957.2 27
Zea mays XM_008681721.3 20
Setaria viridis v2.1|Sevir.1G084000.1.p 28
Panicum hallii v3.1|Pahal.1G037500.1.p 25
Liriodendron tulipifera v1.1|Litul.09G117000.2.p 16
Olea europaea XM_023010336.1 16
Solanum lycopersicum XM_026032229.1 7
Coffea arabica XM_027258360.1 17
Coffea eugenioides XM_027304718.1 17
Eucalyptus grandis XM_010033997.3 18
Populus trichocarpa XM_024597423.1 22
Manihot esculenta XM_043955826.1 15
Fragaria vesca XM_011471062.1 28
Malus domestica XM_029096080.1 21
Prunus persica v2.1|Prupe.2G035500.1.p 14
Medicago truncatula XM_024772672.2 34
Glycine soja XM_028335290.1 29
Vigna unguiculata v1.2|Vigun11g010000.1.p 27
Citrus clementina XM_006420655.2 20
Gossypium raimondii XM_012594130.1 19
Theobroma cacao v2.1|Thecc.10G157200.1.p 14
Carica papaya XM_022056764.1 13
Tarenaya hassleriana XP_010534197.1 17
Capsella rubella v1.1|Carub.0004s2219.1.p 26
Arabidopsis thaliana NM_129561.4 25
Brassica rapa v1.3|Brara.C02049.1.p 31
Brassica oleracea XM_013773073.1 30
Brassica napus XM_013879090.2 30
Marchantia polymorpha Mapoly0004s0120.1 8
Physcomitrium patens XM_024540980.1 11
Anthoceros angustus MU008181.1 0
Physcomitrium patens XM_024544777.1 0
Marchantia polymorpha Mapoly0006s0043.1 0
Selaginella moellendorffii XM_024677492.1 0
Ceratopteris richardii v2.1|Ceric.12G053900.1.p 0
Pinus tabuliformis AJP06334.1 0
Thuja plicata v3.1c|Thupl.29377226s0007.2.p 1
Ginkgo biloba KJ473681.1 0
Cycas revoluta KJ473665.1 0
Amborella trichopoda XM_020675675.1 1
Nymphaea colorata XM_031619240.1 1
Liriodendron tulipifera v1.1|Litul.05G101900.1.p 1
Phalaenopsis equestris XM_020716730.1 1
Asparagus officinalis V1.1|evm.model.AsparagusV1_04.2991 1
Musa acuminata v1|GSMUA_Achr8P12010_001 1
Ananas comosus v3|Aco015559.1 2
Oryza sativa XM_015781554.2 0
Zea mays XM_020539025.2 1
Olea europaea XM_023017486.1 0
Solanum lycopersicum XM_010327133.3 1
Coffea arabica XM_027223757.1 0
Coffea eugenioides XM_027324750.1 0
Fragaria vesca XM_011465506.1 1
Prunus persica v2.1|Prupe.5G089300.1.p 1
Malus domestica XM_029101026.1 1
Eucalyptus grandis XM_039311328.1 0
Glycine soja XM_028335728.1 1
Medicago truncatula XM_003611116.4 1
Citrus clementina XM_006440458.2 1
Populus trichocarpa XM_024588301.1 0
Manihot esculenta XM_043953930.1 1
Gossypium raimondii XM_012633252.1 0
Theobroma cacao v2.1|Thecc.03G270300.1.p 1
Carica papaya XM_022046129.1 1
Tarenaya hassleriana XP_010533720.1 2
Brassica napus XM_013786007.2 0
Brassica oleracea XM_013757626.1 0
Arabidopsis thaliana NM_001334063.1 1
Capsella rubella v1.1|Carub.0002s0198.2.p 1
1.6

Fig. S4F

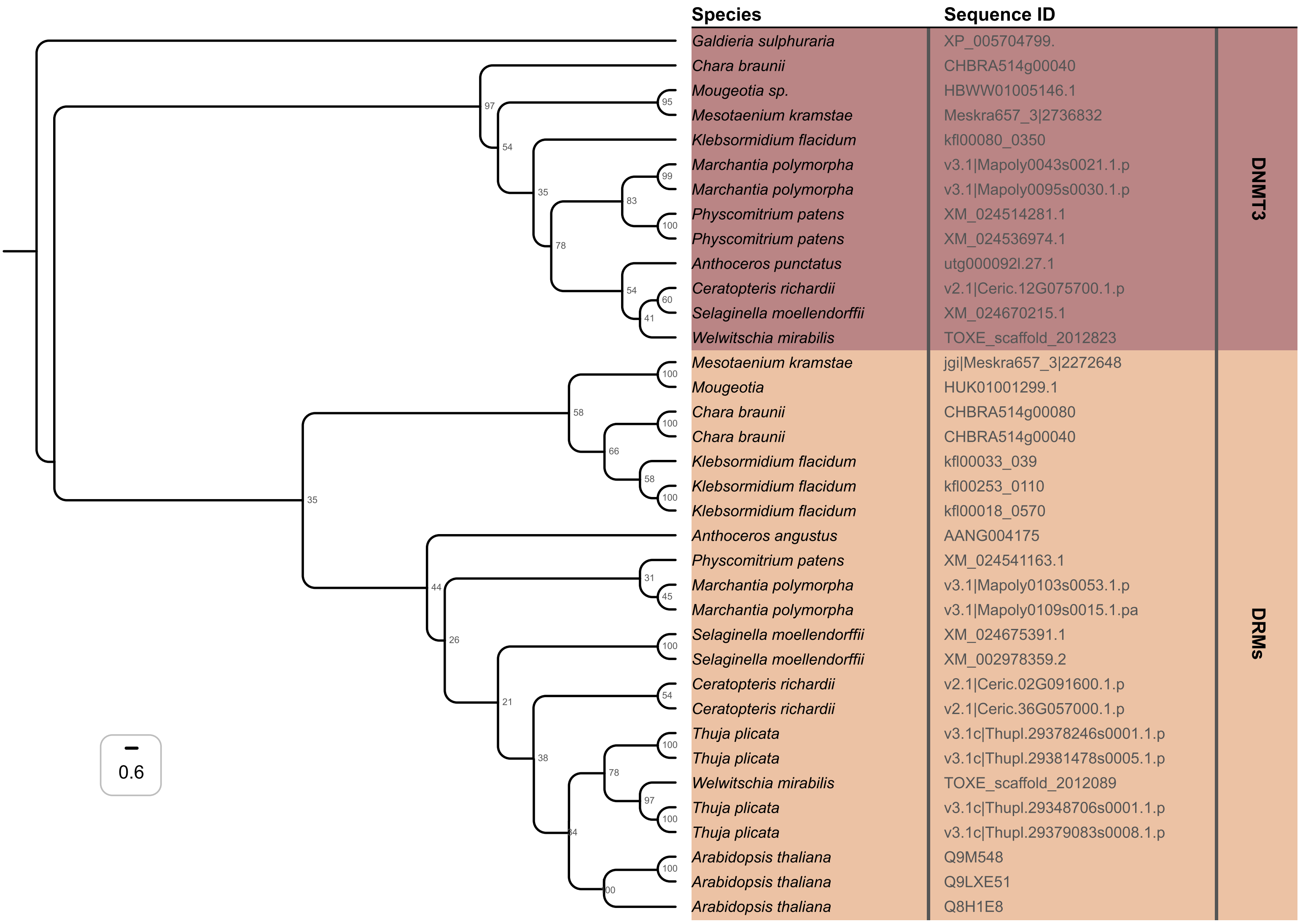
Species
Sequence ID
Galdieria sulphuraria XP_005704799.
Chara braunii CHBRA514g00040
Mougeotia sp. HBWW01005146.1
Mesotaenium kramstae Meskra657_3|2736832
Klebsormidium flacidum kfl00080_0350
Marchantia polymorpha v3.1|Mapoly0043s0021.1.p
Marchantia polymorpha v3.1|Mapoly0095s0030.1.p
Physcomitrium patens XM_024514281.1
Physcomitrium patens XM_024536974.1
Anthoceros punctatus utg000092l.27.1
Ceratopteris richardii v2.1|Ceric.12G075700.1.p
Selaginella moellendorffii XM_024670215.1
Welwitschia mirabilis TOXE_scaffold_2012823
Mesotaenium kramstae jgi|Meskra657_3|2272648
Mougeotia HUK01001299.1
Chara braunii CHBRA514g00080
Chara braunii CHBRA514g00040
Klebsormidium flacidum kfl00033_039
Klebsormidium flacidum kfl00253_0110
Klebsormidium flacidum kfl00018_0570
Anthoceros angustus AANG004175
Physcomitrium patens XM_024541163.1
Marchantia polymorpha v3.1|Mapoly0103s0053.1.p
Marchantia polymorpha v3.1|Mapoly0109s0015.1.pa
Selaginella moellendorffii XM_024675391.1
Selaginella moellendorffii XM_002978359.2
Ceratopteris richardii v2.1|Ceric.02G091600.1.p
Ceratopteris richardii v2.1|Ceric.36G057000.1.p
Thuja plicata v3.1c|Thupl.29378246s0001.1.p
Thuja plicata v3.1c|Thupl.29381478s0005.1.p
Welwitschia mirabilis TOXE_scaffold_2012089
Thuja plicata v3.1c|Thupl.29348706s0001.1.p
Thuja plicata v3.1c|Thupl.29379083s0008.1.p
Arabidopsis thaliana Q9M548
Arabidopsis thaliana Q9LXE51
Arabidopsis thaliana Q8H1E8
DNMT3
DRMs
0.6

Fig. S4G

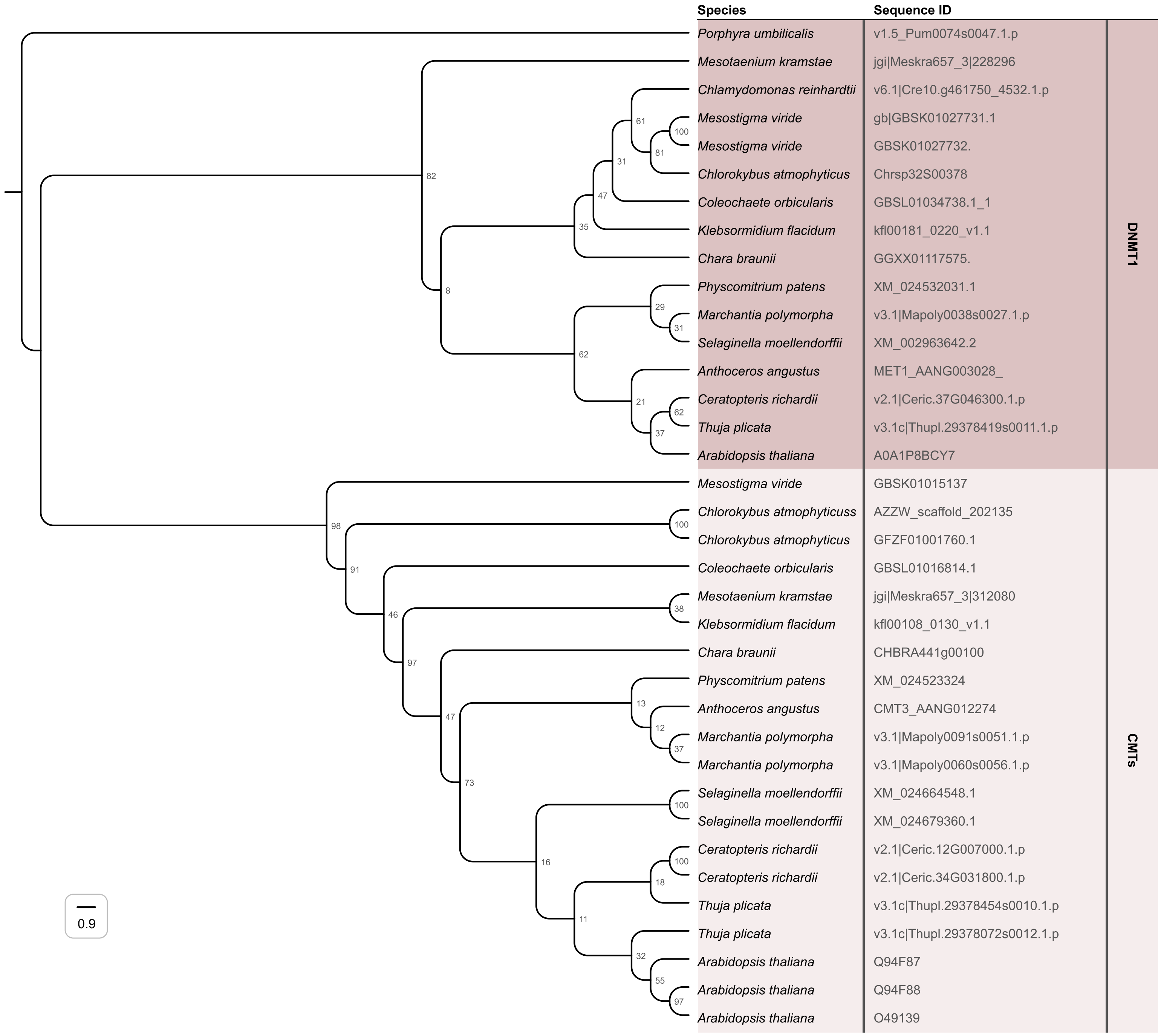

Species
Sequence ID
Porphyra umbilicalis
v1.5_Pum0074s0047.1.p
Mesotaenium kramstae
jgi|Meskra657_3|228296
Chlamydomonas reinhardtii
v6.1|Cre10.g461750_4532.1.p
Mesostigma viride
gb|GBSK01027731.1
Mesostigma viride
GBSK01027732.
Chlorokybus atmophyticus
Chrsp32S00378
Coleochaete orbicularis
GBSL01034738.1_1
Klebsormidium flacidum
kfl00181_0220_v1.1
Chara braunii
GGXX01117575.
Physcomitrium patens
XM_024532031.1
Marchantia polymorpha
v3.1|Mapoly0038s0027.1.p
Selaginella moellendorffii
XM_002963642.2
Anthoceros angustus
MET1_AANG003028_
Ceratopteris richardii
v2.1|Ceric.37G046300.1.p
Thuja plicata
v3.1c|Thupl.29378419s0011.1.p
Arabidopsis thaliana
A0A1P8BCY7
Mesostigma viride
GBSK01015137
Chlorokybus atmophyticuss
AZZW_scaffold_202135
Chlorokybus atmophyticus
GFZF01001760.1
Coleochaete orbicularis
GBSL01016814.1
Mesotaenium kramstae
jgi|Meskra657_3|312080
Klebsormidium flacidum
kfl00108_0130_v1.1
Chara braunii
CHBRA441g00100
Physcomitrium patens
XM_024523324
Anthoceros angustus
CMT3_AANG012274
Marchantia polymorpha
v3.1|Mapoly0091s0051.1.p
Marchantia polymorpha
v3.1|Mapoly0060s0056.1.p
Selaginella moellendorffii
XM_024664548.1
Selaginella moellendorffii
XM_024679360.1
Ceratopteris richardii
v2.1|Ceric.12G007000.1.p
Ceratopteris richardii
v2.1|Ceric.34G031800.1.p
Thuja plicata
v3.1c|Thupl.29378454s0010.1.p
Thuja plicata
v3.1c|Thupl.29378072s0012.1.p
Arabidopsis thaliana
Q94F87
Arabidopsis thaliana
Q94F88
Arabidopsis thaliana
O49139
DNMT1
CMTs
0.9

Fig. S4H

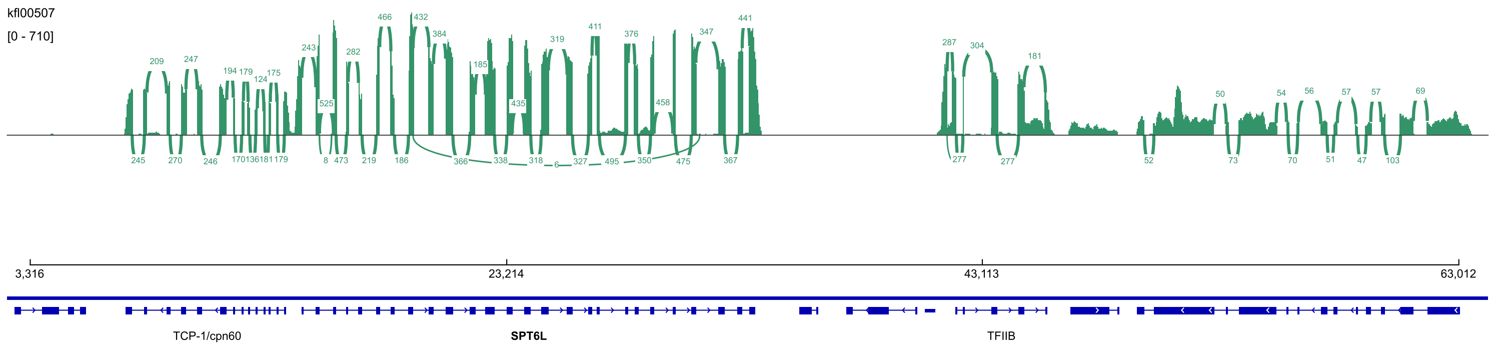


Fig. S4I

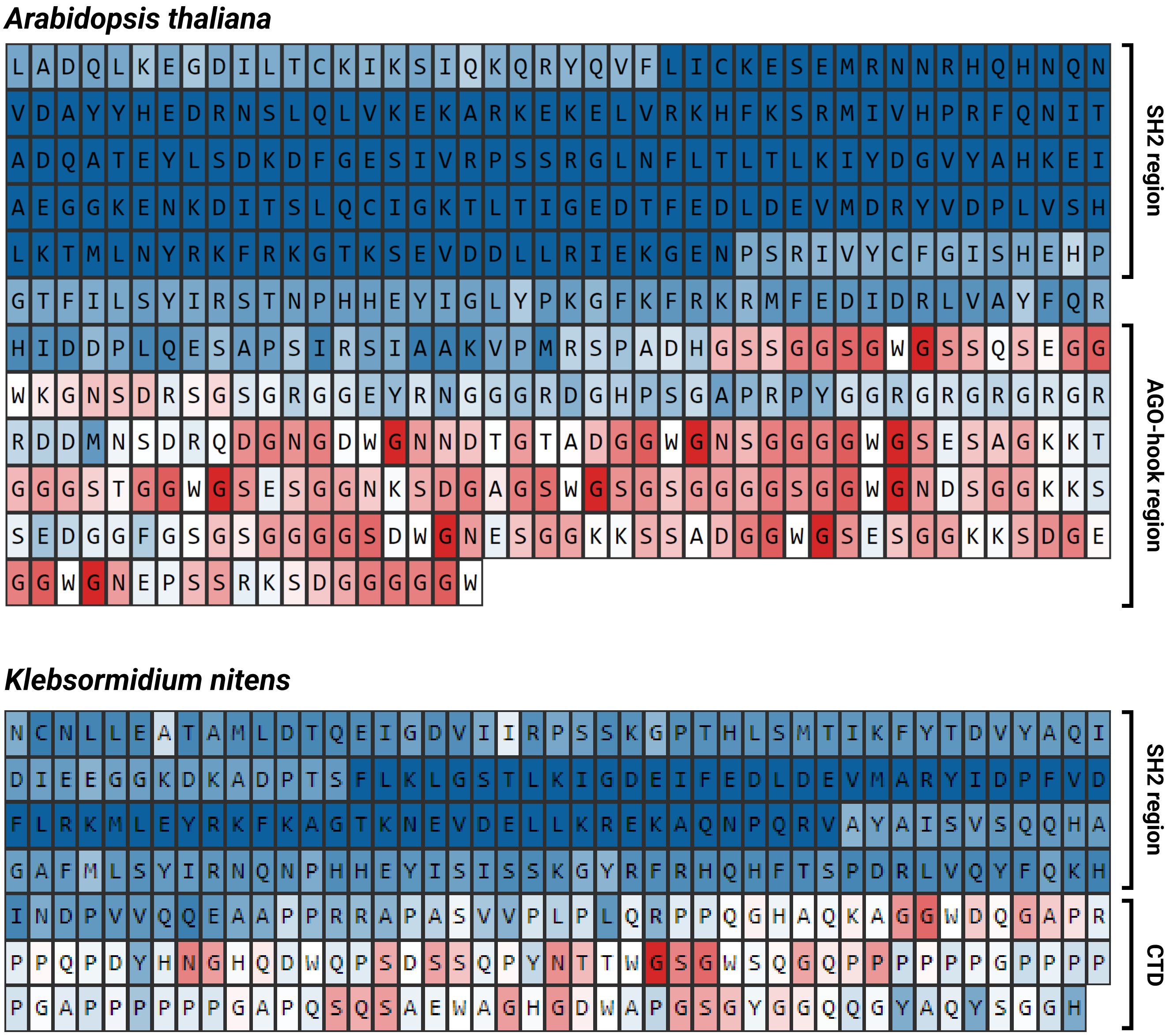
